# Dona Flor and her two husbands: Discovery of novel HDAC6/AKT2 inhibitors for myeloid cancer treatment

**DOI:** 10.1101/2024.11.30.626092

**Authors:** Karoline B. Waitman, Holli-Joi Martin, Jorge A. E. G. Carlos, Rodolpho C. Braga, Vinícius A. M. Souza, Cleber C. Melo-Filho, Sebastian Hilscher, Mônica F. Z. J. Toledo, Maurício T. Tavares, Letícia V. Costa-Lotufo, João A. Machado-Neto, Mike Schutkowski, Wolfgang Sippl, Thales Kronenberger, Vinicius M. Alves, Roberto Parise-Filho, Eugene N. Muratov

## Abstract

Hematological cancer treatment with hybrid kinase/HDAC inhibitors is a novel strategy to overcome the challenge of acquired resistance to drugs. We collected IC_50_ datasets from the ChEMBL database for 13 cancer cell lines (72 h cytotoxicity, measured by MTT), known inhibitors for 38 kinases, and 10 HDACs isoforms, that we identified by target fishing and literature review. The data was subjected to rigorous biological and chemical curation leaving the final datasets ranging from 76 to 8173 compounds depending on the target. We generated Random Forest classification models, whereby 14 showed greater than 80% predictability after 5-fold external cross-validation. We screened 30 hybrid kinase/HDAC inhibitor analogs through each of these models. Fragment-contribution maps were constructed to aid the understanding of SARs and the optimization of these compounds as selective kinase/HDAC inhibitors for cancer treatment. Among the predicted compounds, 9 representative hybrids were synthesized and subjected to biological evaluation to validate the models. We observed high hit rates after biological testing for the following models: K562 (62.5%), MV4-11 (75.0%), MM1S (100%), NB-4 (62.5%), U937 (75.0), and HDAC6 (86.0%). This aided the identification of **6b** and **6k** as potent anticancer inhibitors with IC_50_ of 0.2-0.8 µM in three cancer cell lines, linked to HDAC6 inhibition below 2 nM, and blockade of AKT2 phosphorylation at 2 μM, validating the ability of our models to predict novel drug candidates.

**Graphical Abstract:** 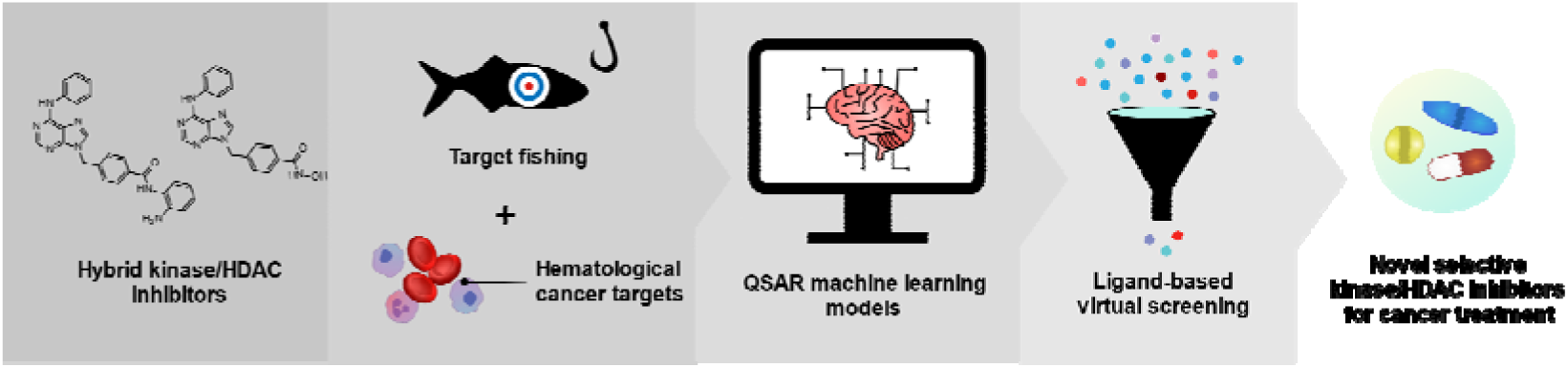

**Highlights:**

- Novel kinase/HDAC inhibitors for cancer treatment were found using machine learning
- 61 QSAR models for hematological cancers and its targets were built and validated
- K562, MV4-11, MM1S, NB-4, U937, and HDAC6 models had hit rates above 62.5% in tests
- **6b** and **6k** presented potent IC_50_ of 0.2-0.8 µM in three cancer cell lines
- **6b** and **6k** inhibited HDAC6 below 2 nM, and blockade of AKT2 phosphorylation at 2 μM

## Introduction

> “To be happy, you need us both.”[1]

The global burden of cancer is immense, accounting for one in six deaths worldwide as of 2020 [2]. There is an urgent need for new therapeutic and technological approaches to overcome it. Among those is the advent of precision medicine, with targeted therapies that block specific survival pathways for cancer cells, increasing treatment efficacy [3]. Such treatments are slowly improving cancer statistics, with slight increases in five-year survival rates for many cancers and declines in global age-standardized death rates in the last decade [4]. Among a myriad of validated targets, kinases have been established among the most promising ones, with over 70 approved inhibitors in clinical use [5]. Imatinib embodied most of the approved kinase inhibitors, and under-development drugs for oncological indications (**Figure 1A**), due to their role in inhibiting cell growth and replication in tumors bearing Philadelphia chromosome [6]. Epigenetic targets, as histone deacetylases (HDACs) are emerging as promising targets, since their inhibitors can reestablish the expression of tumor suppressor genes. For example, treatment with vorinostat improved patient prognosis, especially in hematological malignancies (**Figure 1B**) [7].

**Figure 1.**
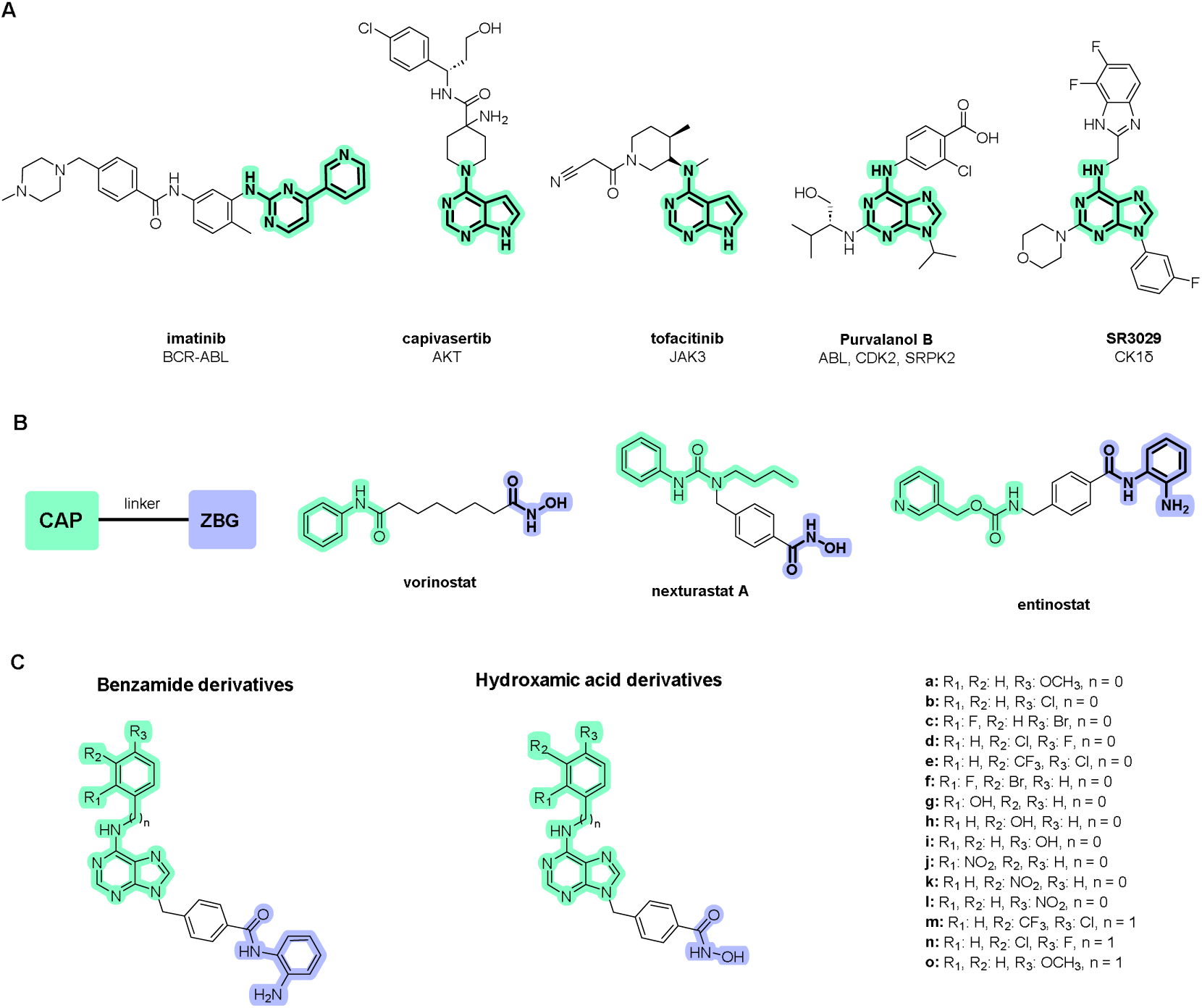
Chemical structures of kinase inhibitors, HDAC inhibitors, and hybrid kinase/HDAC inhibitors. A) Chemical structures of kinase inhibitors with hinge-binding groups (HBG) are highlighted in light green. B) General pharmacophore and chemical structures of HDAC inhibitors, depicting the CAP in light green, the linker in black, and the zinc-binding group (ZBG) in purple. C) Library of hybrid kinase/HDAC inhibitors and their identification. In green is depicted the area for kinase interaction/CAP in HDAC, in black is the linker portion, and in purple, is the ZBG.

However, acquired treatment resistance remains a challenge [8]. One counter strategy is the development of dual hybrid inhibitors, *i.e.* inhibitors able to act on two different targets, which reduces the resistance onset since mutations on two distinct signaling pathways would be needed for that [9]. Like Dona Flor, in Jorge Amado’s book, that only finds fulfilment with her two husbands, one at a time [1], these inhibitor’s dual targeting allow they to fulfill their role by leading to potent, synergistic response in cancer cells, sometimes able to induce apoptosis even in resistant lineages [10] and achieve clinical trials [11–13].

So far, no dual kinase/HDAC inhibitors have been clinically approved, but the conventional drug discovery pipeline is resource intensive [14]. The incorporation of computational approaches, such as machine learning (ML) can use available data to speed up this process and mitigate failure rates [15,16]. For example, quantitative structure-activity relationships (QSARs) models can use information from available chemical databases such as ChEMBL [17] to predict the activity of novel sets of compounds within the applicability domain[18,19] and generate similarity maps to distinguish between structural fragments contributing to the activity [20].

There are few published studies employing integrative approaches that combine both cell and target-based data to develop novel hybrid kinase/HDAC inhibitors for hematological cancer treatment. In most cases, QSAR models have been built to predict the activity in solid cancer cell lines [21–24] or focused solely on one target [25]. However, even among the ones that explored hematological cancers, some models were not subjected to biological evaluation [26–28], and none was curated for specific assays and/or time intervals which would allow biologists not only to easily replicate their findings but also experimentally validate their novel predictions.

To fill this gap, the present study constructed and validated QSAR models, using IC_50_ datasets of cancer cell lines and their putative targets obtained by target fishing, to aid the development of novel selective drugs for hematological cancer treatment. These models were used to screen a small library of hybrid kinase/HDAC inhibitors (**Figure 1C**) designed based on previous studies [29], guiding the synthesis and biological evaluation of these compounds, as well as the biological validation of novel QSAR models feasible for further usage as a prediction tool in drug design programs.

## Results and Discussion

We started gathering biological data from inhibitors tested both on hematological cancer cell lines and its associated targets from the ChEMBL database to construct QSAR models. Additionally, the designed kinase/HDAC hybrids were submitted to target fishing and phenotypic screening to predict its putative kinase/HDAC targets in a less biased approach, which were also modeled. These models were then used to assess the dual activity of the compounds in a ligand-based virtual screening (LBVS), that was later subjected to biological assessment. A general scheme of the workflow for the conduction of this study can be seen in **Figure 2**.

**Figure 2.**
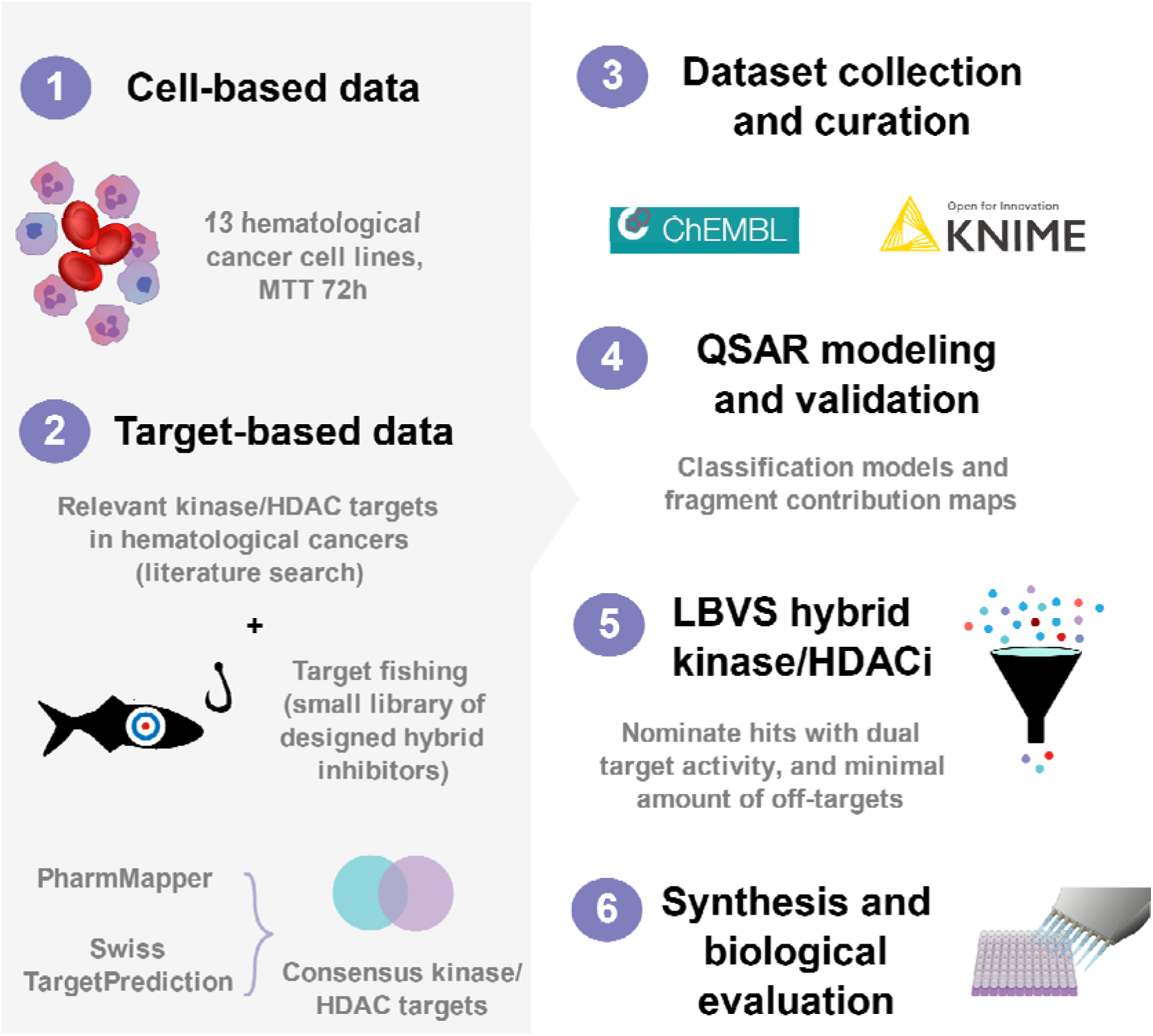
Workflow employed for the study execution. Biological data from inhibitors from hematological cancer cell lines (1) and associated targets (2) were retrieved from the ChEMBL database (3) to construct QSAR models (4). Additionally, the designed kinase/HDAC hybrids were submitted to target fishing (2) and phenotypic screening to predict its putative kinase/HDAC targets in a less biased approach, which were also modeled. These models were then used to assess the dual activity of the compounds in a ligand-based virtual screening (LBVS, 5), that was later subjected to biological assessment (6).

### Cancer cell models, target selection, QSAR modeling, and statistical validation

A set of 13 hematological cancer cell lines was selected to provide biological diversity of cancer models (**Supplementary Table S1**), which could be associated with different pathology mechanisms, and exhibited overexpression of different biological targets [30,31]. Twelve kinases linked to hematological malignancies and HDACs 1-10 were chosen based on literature reviews [30,32–34] (**Table 1**), and to provide a more unbiased approach to enzyme selection, an additional target fishing analysis step was also conducted, using the library compounds as baits (**Figure 1C**).

**Table 1.**
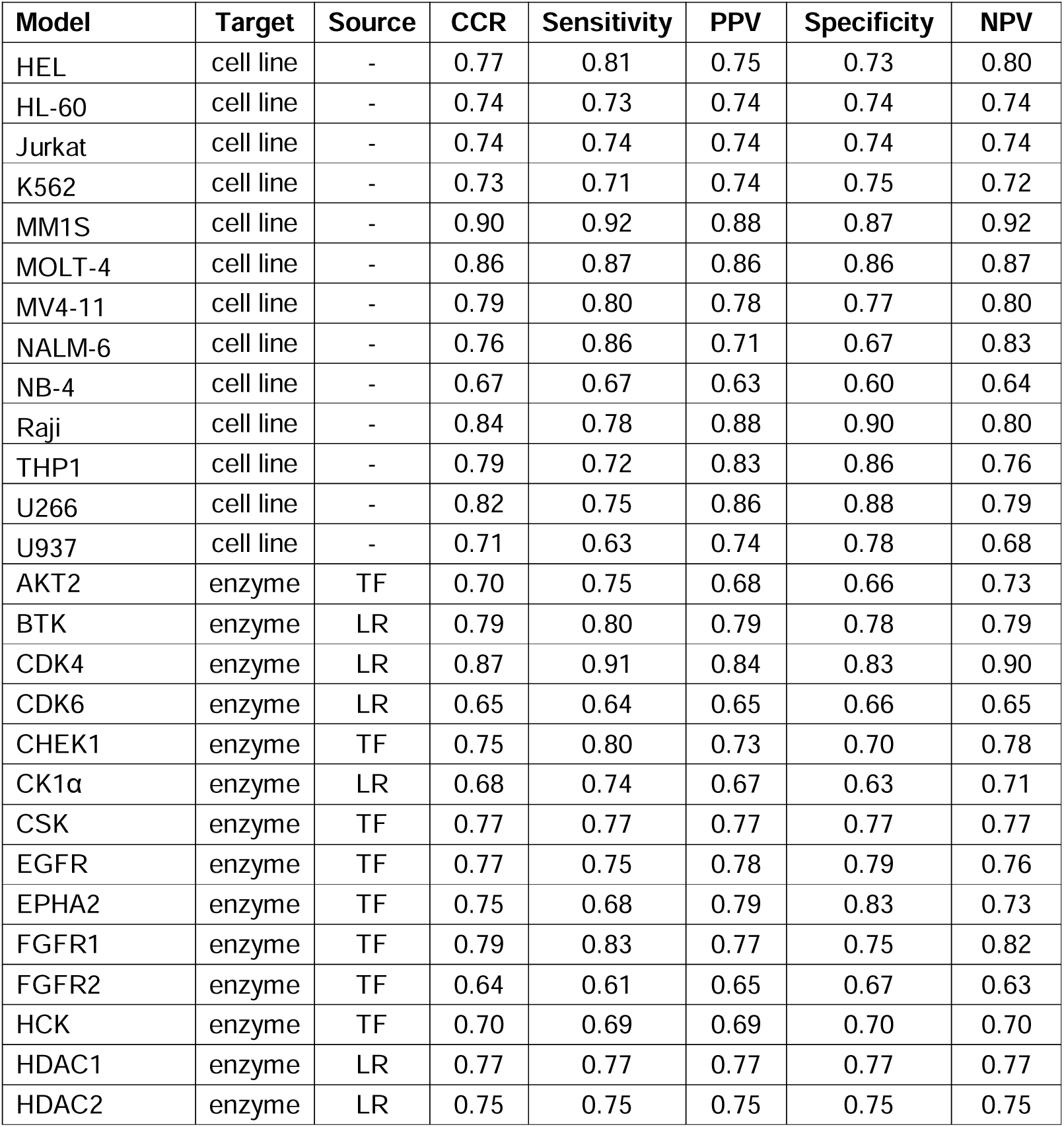
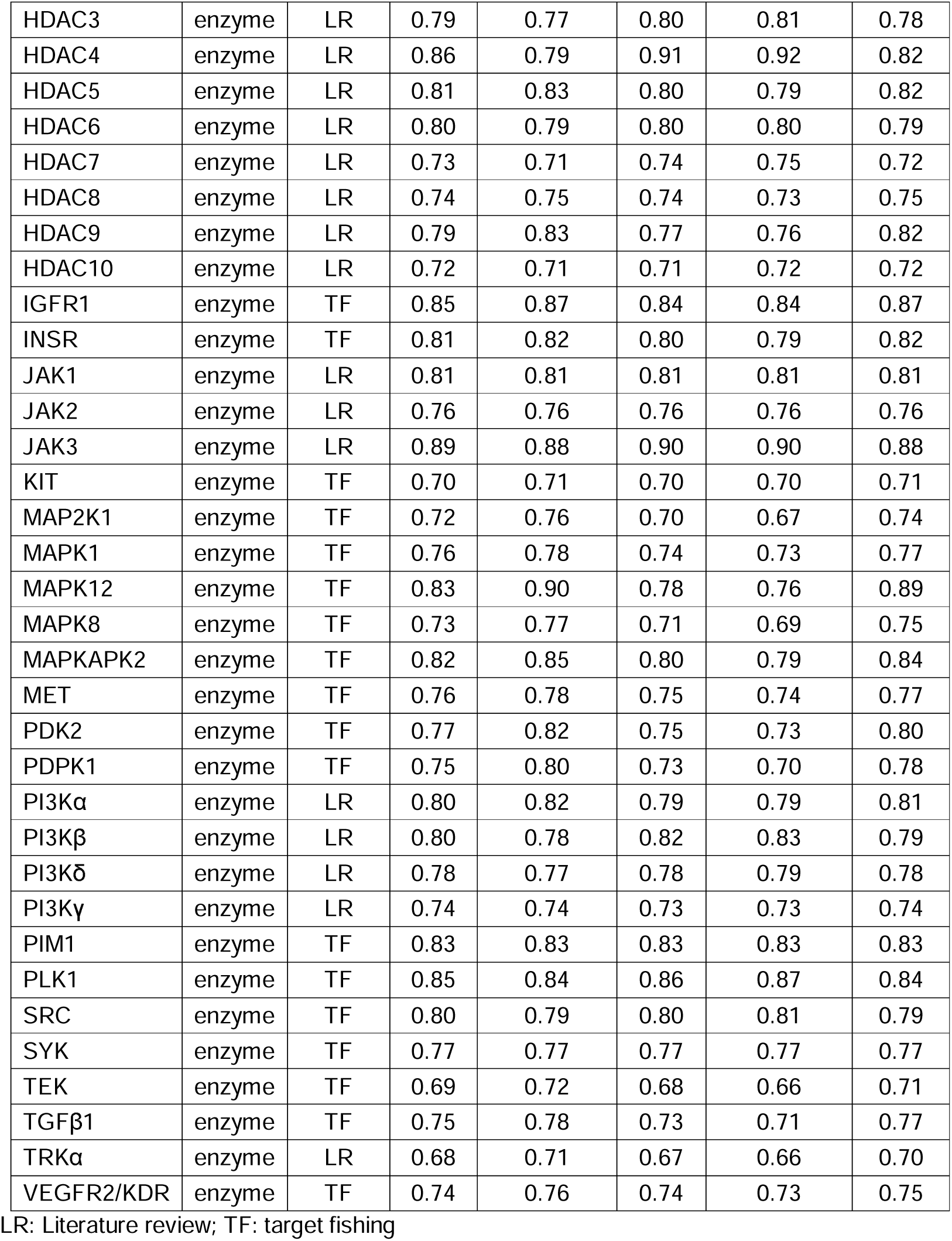
Targets retrieved for the hybrid compounds, and the statistical performances of their corresponding QSAR models assessed by 5-fold external validation.

Two different approaches were employed to predict the putative targets of the hybrid kinase/HDAC compounds, namely PharmMapper and SwissTargetPrediction. PharmMapper relies upon a reverse pharmacophore-mapping strategy [35], while SwissTargetPrediction employs a combination of 2D and 3D similarity values to predict the targets of the studied molecules and is considered one of the most reliable sources for target prediction [36,37]. The combination of both tools through a consensus analysis allows a more robust identification of the putative targets, potentially leading to a more assertive process. A total of 26 kinases were retrieved from the target fishing (**Table 1**). The number of putative targets found for each compound in PharmMapper, SwissTargetPrediction, and their consensus is described in **Supplementary Tables S2** and **S3**.

Following cancer cell and target selection, QSAR models were constructed using biological data retrieved from ChEMBL for each enzyme. The data was thoroughly curated to ensure removal standardization of chemical data, removal of duplicates, and avoid errors in input data [38]. Given the high variability of the cellular environment, an additional manual curation step was performed for cancer cell models, to ensure only IC_50_ measurements by MTT assays in the interval 0-72h were included in the data. On average 16.5% of the initial dataset was removed during data curation, with 5 datasets losing above 30% of the initial data. All the models were constructed employing the KNIME software, and 5-fold external validation with the RF algorithm. The performance characteristics of the 13 cancer cells and 48 enzymatic models are described in **Table 1**, and the activity threshold selected for each enzyme and dataset size are described in **Supplementary Table S4**.

Most models presented good performance with predictive capacity above 70% as evaluated by CCR, sensitivity, specificity, PPV, and NPV values. 19 models presented the best performances with correct classification rates above 80%, following OECD standards [39]. Only CDK6, CK1α, and FGFR2 presented predictability lower than 65%, which can be in part related to the smaller dataset size for these enzymes (**Supplementary Table S4**). Another possible reason for the differences in the performances of the models can be related to inaccurate measures in the experimental biological assays, which is an issue in modeling cheminformatics data [40–42].

### LBVS can identify potential anticancer kinase/HDAC inhibitors

Using the validated models, the LBVS of the 30 hybrid kinase/HDAC compounds was performed to predict their activity in the cancer cell and enzyme models. The results obtained in the screening can be found in **Supplementary Tables S5** and **S6**. It is important to notice that all predicted outcomes depend on the thresholds established for the targets (**Supplementary Table S4**), and on the subjective factors involved in the biological assays from which the data was collected, such as type and concentration of substrate, type of buffer, differences in cel culture medium, among others.

All hybrids were classified as active in at least two cancer cell models, with most being predicted as active in at least 4 different cell types (**Supplementary Table S5**). All hybrids were predicted to be active in the erythroleukemia cell HEL, with benzamides nominated as active in MV4-11, NB-4, and THP1. Hydroxamic acid derivatives were predicted to be active in the erythroleukemia model K562, and HL-60, Jurkat which are acute promyelocytic leukemia and acute T-cell leukemia models. In enzymatic models, all hybrid compounds were predicted active on average against six different targets (**Supplementary Table S6**), however, there were differences in their predicted preferred enzyme target. For instance, benzamides would more often be predicted to target kinases than HDACs, and the opposite happened for hydroxamic acids, which were predicted to be more active in HDACs. That difference can be attributed to the zinc-binding group employed for HDAC interaction, with hydroxamic acids being less selective than benzamides for the isoform inhibition, which agrees with the literature [43].

The increased affinity for kinases seen on benzamide derivatives can be attributed to the interaction of the additional nitrogen group with DFG-out residues of the kinase since this group can be found in FDA-approved type II kinase inhibitors such as imatinib [44]. However, these outcomes vary according to the kinase family which is being targeted benzamides were predicted active in the tyrosine kinase group (TK) more frequently than the other hybrids (58 positive outcomes against 11 for hydroxamic acids). No significant difference was noted for the capacity of targeting serine/threonine kinases, with both hydroxamic acids and benzamides presenting around 20 to 25 positive outcomes. Overall, 70% of the library was predicted active in CK1α, followed by HDAC7, and AKT2 with 60% of the hybrids classified as actives.

Taking the activity of both targets into account it was noticed that most hybrids were predicted to have the desired kinase/HDAC inhibition. However, one of the challenges in developing hybrid inhibitors is to balance the dual activity in the desired targets with the off-target [45]. In this sense, the usage of QSAR allows for a quick and cheap assessment of the expected off-targets to direct the synthesis of compounds [15].

### Synthesis of representative hybrid compounds

To verify whether the hybrids indeed displayed the predicted profile, a set of representative compounds (**5a-c, 6a-d**, and **6k**), prioritized from our models with different selectivity profiles (**Supplementary Table S7)** was synthesized and characterized. Selected compounds were nominated as active for both kinases and HDACs with selective profile were favored but representative of both hydroxamic acids and benzamides were chosen to compare the predictions with the biological activity. **Scheme 1** shows the synthetic route employed to prepare the test compounds. 6-chloropurine (**1**) was reacted with substituted anilines to give the intermediates **2a-d**, and **2k**. These intermediates were reacted with 4-(bromomethyl)benzoate to provide esters **3a-d**, and **3k. 3b-c** were subjected to basic hydrolysis to prepare carboxylic acids **4b-c** [46], and then coupled with 1,2-diaminobenzene to yield benzamides **5b-c** [47]. Alternatively, esters **3a-d**, and **3k** reacted with hydroxylamine to produce hydroxamic acids **6a-d**, and **6k** [29].

**Scheme 1:**
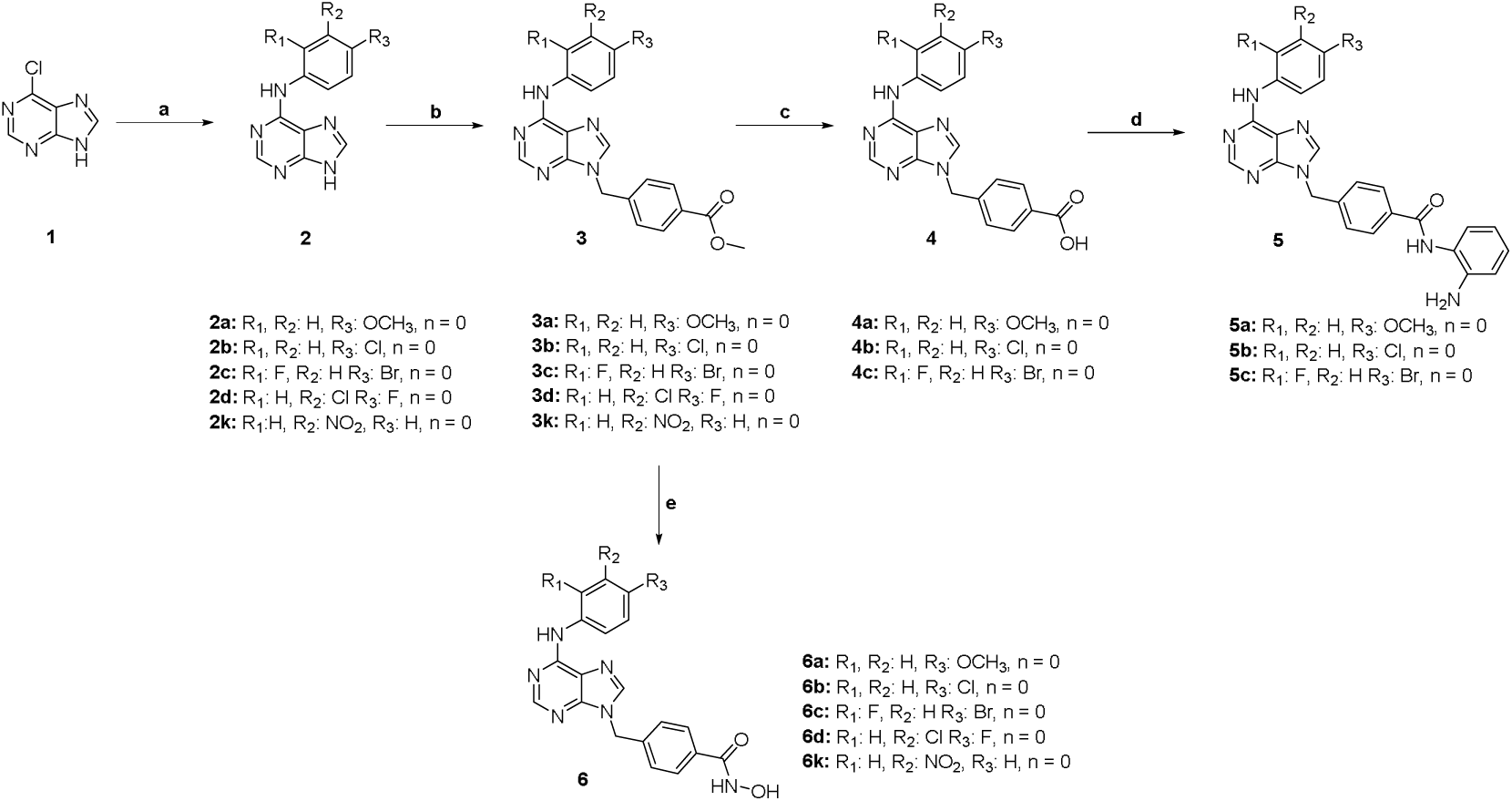
Synthesis of test compounds **5b-c**, **6a-d**, and **6k**. Reactants and conditions: a) anilines (1 eq.), concentrated HCl (0.8 eq.), isopropanol, reflux, 18 h; b) methyl 4-(bromomethyl)benzoate (1 eq.), K_2_CO_3_ (3 eq.), acetone, reflux, 18 h; c) KOH 2M (6.6 eq.), THF/H_2_O (2 mL/mmol), 24-48h; [46] d) Intermediate 4 (1 eq.), 1,2-diaminobenzene (1 eq.), 1-ethyl-3-(3-dimethylaminopropil)carbodiimide - EDC (2 eq.), 1-hydroxybenzotriazole - HOBt (cat.), triethylamine (2.5 eq.), 0 °C – r.t., 18 h [47]; e) NH_2_OH (50 eq.), NaOH (8 eq.), THF-MeOH-dioxane (1:1:1), 0 °C – r.t., 2h [29].

### Nominated hit compounds displayed cytotoxicity in hematological cancer cell lines

The synthesized hybrids were subjected to IC_50_ determination in the 13 hematological cancer cells modeled. **Table 2** shows the results of the biological assessment of the hybrids in the cancer cell lines and their predictability rates. All hybrids presented the desired anticancer activity below 0.8 µM in at least one cancer cell line, with **6b** showing the highest potency, with IC_50_ of 60 nM in the acute myeloid leukemia cell MV4-11. Except for Jurkat, both hydroxamic acids and benzamides showed similar potency in the cancer models, with MOLT-4, MV4-11, and THP1 presenting the highest vulnerabilities to the compounds, and K562, HEL, and U937 being the least susceptible. Since the MV4-11 and THP1 are respectively macrophage and monocyte malignancies that suggests the clinical indication of such compounds should focus on tumors of the mononuclear phagocyte system.

**Table 2.** IC_50_ of the representative compounds in cancer cells compared to model predictions and CCR. Assessment of cancer cell QSAR models in biological testing.

| Cmp | IC<sub>50</sub> (μM) |  |  |  |  |  |  |  |  |  |  |  |  |
| --- | --- | --- | --- | --- | --- | --- | --- | --- | --- | --- | --- | --- | --- |
|  | HEL | Jurkat | K562 | HL-60 | MM1S | MOLT-4 | MV4-11 | NALM-6 | NB-4 | Raji | THP1 | U266 | U937 |
| 6a | 0.80 | 0.70 | 3.70 | 1.98 | 2.04 | 0.46 | 0.58 | 2.01 | 0.87 | 0.43 | 0.42 | 1.11 | 1.23 |
| 6b | 0.77 | 0.91 | 13.81 | 0.62 | 2.64 | 0.19 | 0.06 | 0.41 | 0.41 | 0.36 | 0.19 | 0.63 | 7.91 |
| 6c | 2.46 | 0.63 | 5.18 | 5.98 | 1.87 | 0.53 | 0.77 | 2.12 | 1.41 | 0.23 | 0.93 | 0.75 | 2.16 |
| 6d | 3.54 | 1.20 | 7.35 | 2.55 | 2.69 | 0.66 | 1.14 | 2.02 | 2.34 | 0.87 | 1.61 | 1.46 | 2.94 |
| 6k | 2.07 | 0.89 | 10.44 | 1.86 | 1.56 | 0.92 | 0.28 | 1.90 | 0.58 | 0.30 | 0.53 | 0.31 | 1.99 |
| 5a | 16.76 | 9.74 | >50 | 4.12 | 15.82 | 0.61 | 0.72 | 7.81 | 2.44 | 10.99 | 0.93 | 4.21 | 22.84 |
| 5b | 12.99 | 13.49 | 17.50 | 1.11 | 1.13 | 0.49 | 0.59 | 1.55 | 0.73 | 0.11 | 0.48 | 0.33 | 0.92 |
| 5c | 17.04 | 4.73 | 30.09 | 0.99 | 1.47 | 0.45 | 0.28 | 1.54 | 0.39 | 0.12 | 0.41 | 0.46 | 1.37 |
| Threshold ($\mu\text{M}$) | 2.0 | 0.5 | 1.0 | 1.0 | 0.5 | 0.5 | 0.05 | 2.0 | 2.0 | 1.0 | 2.0 | 1.0 | 1.0 |
| CCR (%) | 37.5 | 50.0 | 62.5 | 37.5 | 100.0 | 50.0 | 75.0 | 50.0 | 62.5 | 11.1 | 50.0 | 37.5 | 75.0 |

Lilac cells: IC_50_ is consistent with the QSAR model predictions, while white cells: IC_50_ values are inconsistent with the model predictions. CCR was displayed as a heat map where red represents higher predictability and purple has lower predictability rates of the models.

Models MM1S, MV4-11, and U937 presented CCR above 75%, K562, and NB-4 CCR above 62.5%, showing satisfactory correlation with the observed biological activity and rendering those as acceptable for further usage as a prediction tool in drug design programs. Additionally, the biological results dialogue with the target activity predictions, since the compounds were nominated active in targets commonly expressed in such hematological malignancies as HDACs, AKTs, JAK3, CK1α, and TRKα [48,49].

### 6b and 6k presented potent HDAC3/6 inhibition and a suboptimal effect on AKT subtypes

All hydroxamic acid hybrids presented higher inhibitory potency in HDAC6 in comparison to HDAC1 and HDAC8. **6a**, **6b**, and **6k** presented IC_50_ < 2 nM, the limit of detection of the method, and a selectivity index above 160-fold in comparison to HDAC1. On the other hand, benzamides **5b** and **5c** presented higher potency and selectivity for HDAC1, especially compared to HDAC8 inhibition (**Table 3**). **6b** and **6k** presented the highest selectivity indexes for HDAC6, 2-3 fold above the selective HDAC6 inhibitor Nexturastat A, so they were further tested in other HDAC isoforms. **6b** and **6k** also presented lower potency in most HDAC isoforms in comparison to HDAC6, except HDAC3, where the IC_50_ was respectively 15.1 nM and below 2.00 nM, suggesting a selective profile for HDAC3/6. There is only one report of similar HDAC3/6 dual activity in the literature [50], which attributes this unusual HDAC affinity to interactions within the L1-loop, which is more flexible in HDAC6, and in HDAC3, has a non-conserved asparagine (N197) in place of a glutamic acid (E203/E208) in HDACs 1 and 2, respectively. Nevertheless, the HDAC3 inhibition was an unexpected event, and its molecular aspects will be further studied in the future.

**Table 3.**
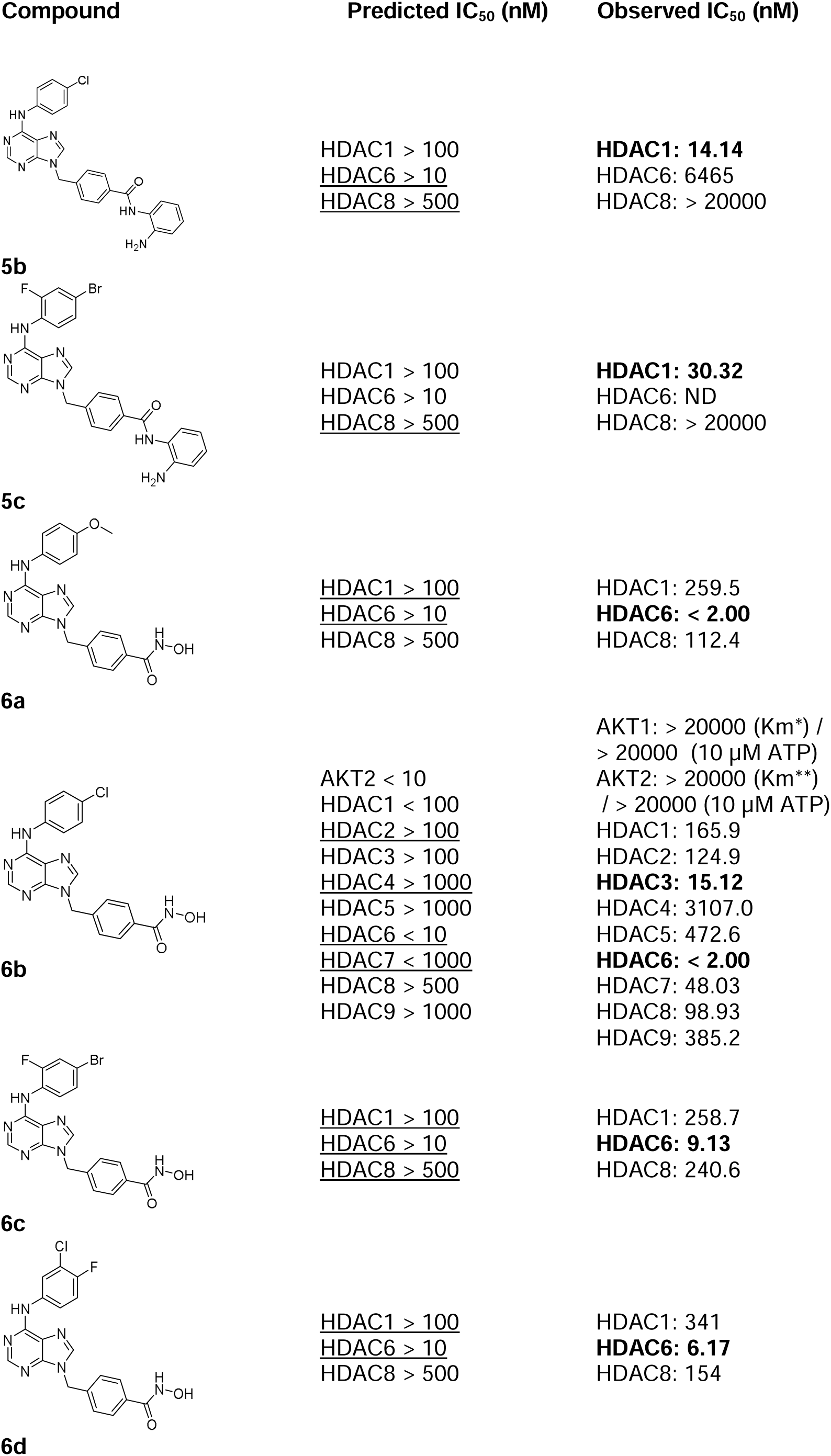
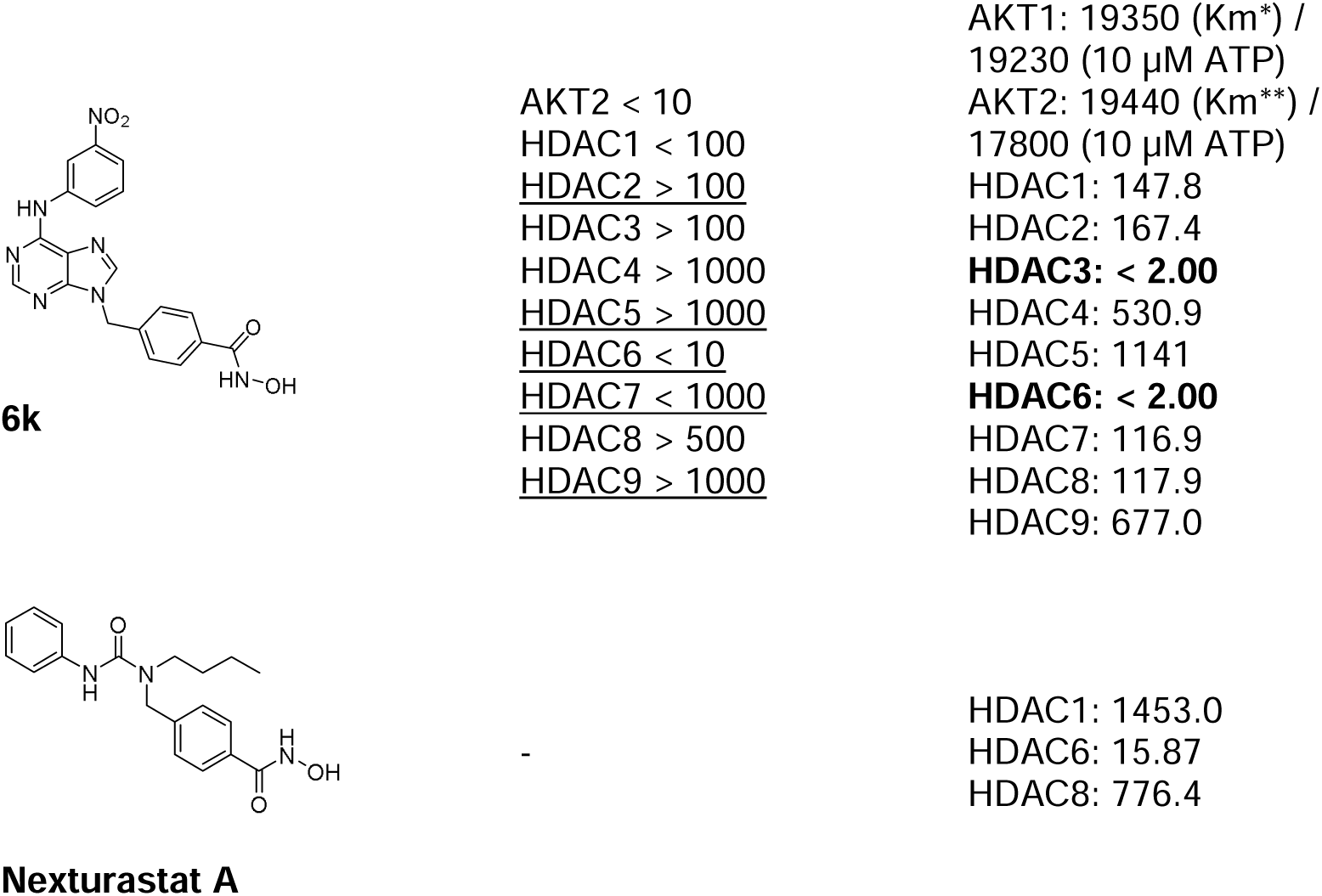
Predicted and observed IC_50_ of the representative compounds and Nexturastat A in HDAC and AKT subtypes.

### Nexturastat A

N.D.: Not determined. Underline: IC_50_ predictions which matched biological results; Bold: IC_50_ below 35 nM. *Km AKT1: 30 µM; **Km AKT2: 100 µM.

Among the target models, HDAC6 presented the best CCR displaying 86% agreement with the biological results, indicating it could be further used to classify the activity of novel HDAC6 inhibitors. HDAC7 inhibition for **6b** and **6k** also matched the model predictions, with **6b** presenting an IC_50_ of 48 nM. Indeed, in these two isoforms, all hydroxamic acid hybrids were nominated as active, which can be related to the sterically bulky phenyl linker found in these compounds, that can interact with the voluminous pocket of both HDAC6 and HDAC7 [51,52].

HDAC1 and HDAC8 models presented a CCR of 43%, indicating a low correlation with the biological activity. Notably HDAC1 models even failed to predict the activity of benzamides in this isoform, despite their selectivity for HDACs1-3 [53]. Model failure in those cases could be a reflex of this ZBG being less represented in databases, biasing the model predictions toward hydroxamic acids. The HDAC8 model, in turn, was able to properly classify benzamides **5b-c** as inactive but incorrectly labeled hydroxamic acids **6a-b**, **6d,** and **6k** as inactive below 500 nM in this isoform. This might indicate that a few compounds with similar structures to the hybrids were tested against HDAC8, biasing the model towards false negative values.

**6b** and **6k** were further tested in AKT1 and AKT2 since this was the kinase where most compounds were predicted to be active, and the model CCR was above 70% (**Table 3**). The hybrids were tested in two different ATP concentrations, namely 10 µM, and their Km (see **Table 3**), to represent the test conditions most frequent in the dataset used in the construction of the models. Unfortunately, **6b** did not inhibit AKT1 and AKT2, and **6k** presented IC_50_ around 17-19 µM in both isoforms, indicating low enzymatic inhibition, contrary to the model predictions. Interestingly, the reduction in ATP concentrations, didn’t increase the potency of the inhibition, suggesting an interaction profile of non-competitive inhibitors [54,55].

In our previous study, **6c** and **6d** had their HDAC1 and HDAC6 IC_50_ tested (IC_50_ HDAC6 184.3 – 341.0 nM, IC_50_ HDAC1: 83.5 – 188.5 nM) [29], however, these results greatly differed from the ones reported here, which could be due to differences in the employed assay substrates. **6c-d** were 3-fold more potent in HDAC1, and 20 to 100-fold more potent in HDAC6. The present study employed substrates developed in-house for monitoring HDAC activity, some even feasible for continuous assays, which could then lead to different results [56], as the ones shown here. These differences in reported IC_50_ values illustrate the challenges in combining biological data for building ML models. The noise found in the dataset due to subjective factors can lead to reduced accuracy in predictions [42]. That might be the reason **6k** AKT2 inhibition was found to be lower than the predicted threshold.

### 6b and 6k reduce the AKT’s expression and autophosphorylation in Jurkat cells

Since compounds **6b** and **6k** were nominated active in AKT, they were further analyzed to verify their ability to inhibit AKT phosphorylation. Western blotting pointed the ability of **6b** to modulate AKT signaling at 1.6 µM, while **6k** was able to modulate it at 3.2 µM, in a lower IC_50_ than the observed AKT inhibition, suggesting the decreased phosphorylation is not linked to blockade of AKT autophosphorylation, and thus its regulation [57].

The high potency of HDAC6 inhibition by both compounds helps to understand their blockade of pAKT expression below 2 µM, like the HDAC inhibitor vorinostat (**Figure 3**). Administration of HDAC inhibitors is linked to reduced pAKT expression, but not non-phosphorylated AKT expression in a dose-dependent manner in different cancer cell lines [58–60], with decreased expression when combined with a kinase inhibitor [61,62], possibly also regulating other kinases that also phosphorylate Ser473 in AKT [63]. A comparison of the potency of the two HDAC inhibitors vorinostat and trichostatin A (TSA) in gallbladder carcinoma cells indicated the latter presented a ten-fold higher potency in SGC-966 cells (10 µM vorinostat; 0.8 µM TSA) [64], which might be related to the ability of one of TSA conformers selectively bind to HDAC6 [65], suggesting increased interaction with this isoform might be important to suppress pAKT expression.

**Figure 3.**
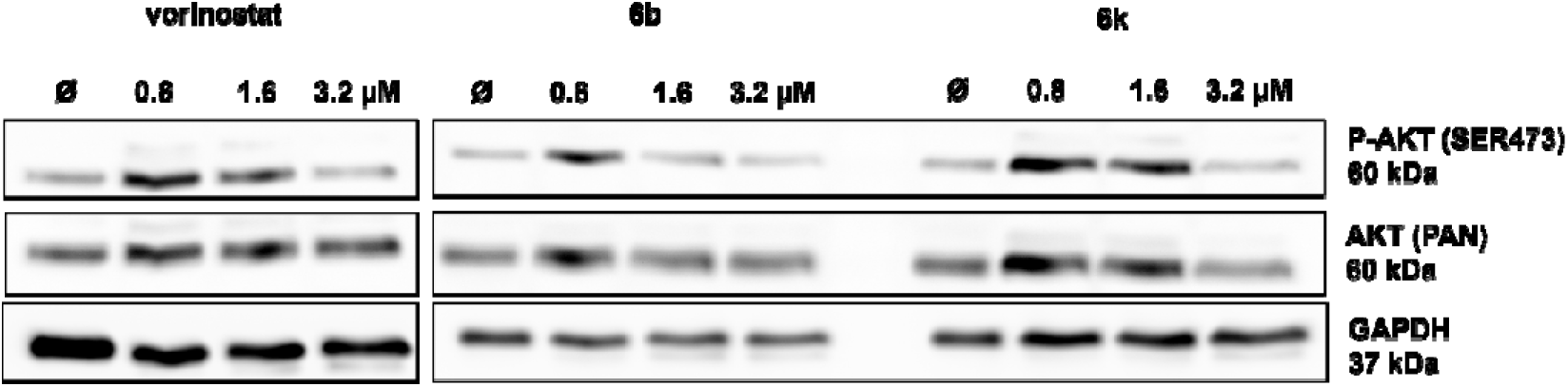
Compounds **6b** and **6k** decrease phosphorylation of AKT. Immunoblotting analysis for p-AKT, AKT, and GAPDH in Jurkat cells after incubation with DMSO (0.8% control), vorinostat, and compounds **6b**, and **6k** for 24h.

### 6b and 6k present favorable fragment-based contributions to the predicted affinity to HDAC6

To better understand the selectivity profile of the designed hybrids and their structure-activity relationships, fragment-based contribution maps were generated and compared to the activity predictions of the models. The region of the scaffold where the most significant differences could be noted among the targets was the aniline moiety. This fragment showed a negative contribution to the predicted activity in HDAC1 and HDAC7, especially in **6k**, and a neutral to positive contribution for HDAC6 and AKT2 in **6b** and **6k** (**Figure 4**). In HDAC7, the nitro group presented a negative contribution for the activity in comparison to the chlorine group from **6b**, which correlates to the higher potency observed in **6b** (IC_50_ = 48 nM), versus **6k** (IC_50_ = 116.9 nM). Additionally, it could be noted that nitro-substituted compounds such as **6k** were among the most selective hybrids designed (**Supplementary Table S6**). Due to its role as a strong electron-withdrawing group, the nitro substituent reduces the acidity of the aniline NH, which might be related to a reduction in polar bonding between the protein and the ligand [66], and that property might be related to the increased selectivity observed.

**Figure 4.**
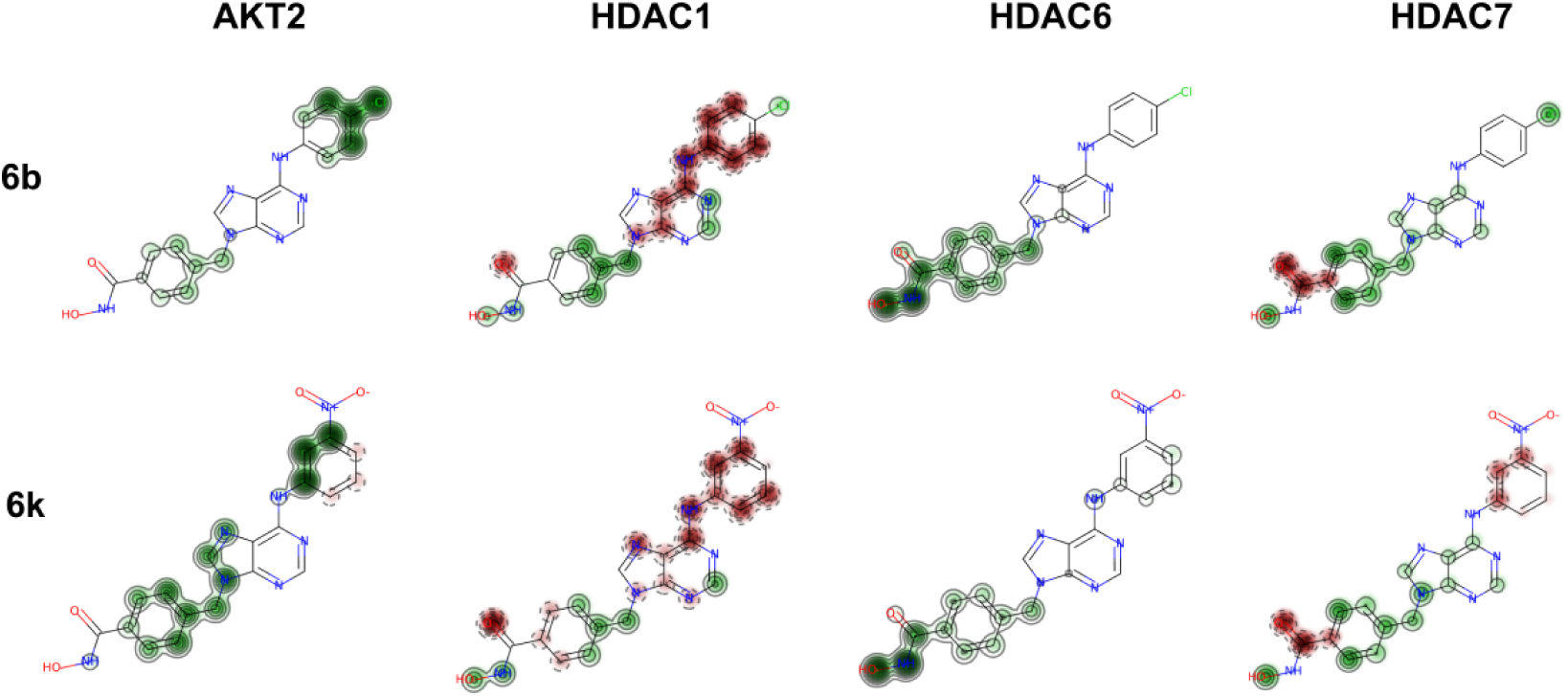
Fragment-based contribution maps of **6b** and **6k** in AKT2, HDAC1, HDAC6, and HDAC7. Green represents a positive contribution to the predicted activity in the target, while red shows the negative contributions. Regions without colors represent areas that don’t change the probability of the predicted outcome in the models.

The hydroxamate-benzyl group made a strong positive contribution to the activity of **6b** and **6k** in HDAC6 (**Figure 4**), while it had a lesser impact on HDACs 1 and 7, probably also contributing to the selectivity observed. This group also presented favorable contributions to the activity in AKT2, which might relate to the polar contacts interactions with residues from the hydrophobic region II (HR-II, **Figure 5A-C**), specifically with R274 and D275. Meanwhile, the **6k**’s nitro-benzyl ring can establish further charge-mediated interactions with W80 and R269 near the gatekeeper portion, which is absent in the **6b**. Those additional interactions are, however, not reflected in either the predicted binding energy, which is lower for **6b**, or the interaction stability as longer interactions with the hinge binding motif are observed (**Figure 5C**). Therefore agreeing with the lower inhibition observed.

**Figure 5.**
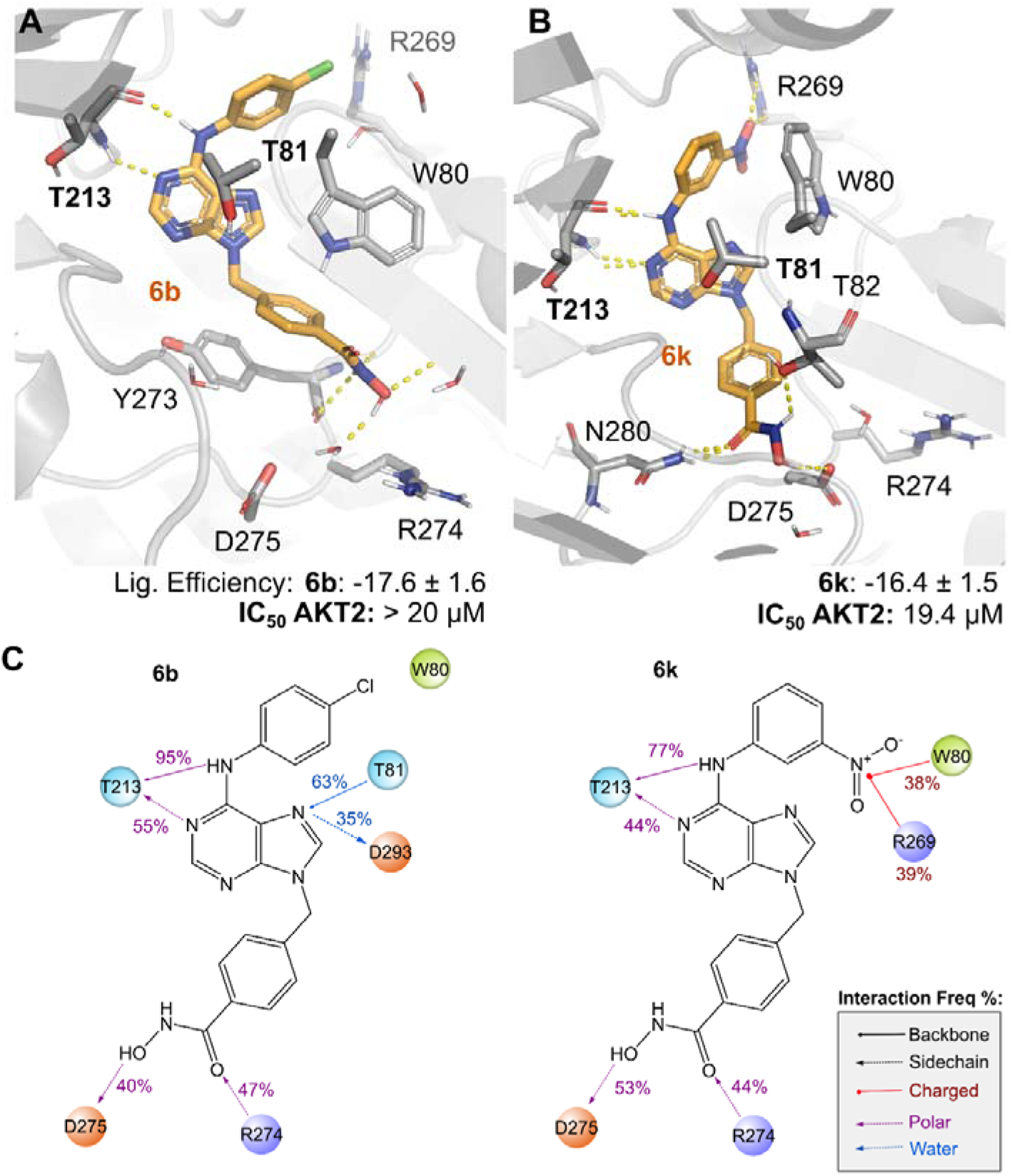
AKT2 binding site (PDB 9C1W) and the proposed binding mode of compounds **6b** (A) and **6k** (B) in AKT2’s catalytic site were generated from relevant frames from their MD simulations (5x500 ns). The predicted binding energies, generated from MM/GBSA (see methods) are depicted as numbers below, alongside the IC_50_ values for comparison). C) Protein-ligand interactions frequency represented over the calculated MD trajectory (5x500 ns), polar contacts with amino acid side chains are depicted in dashed purple lines, while solid lines show interactions with their backbones, similarly water-mediated interactions are blue and charged interactions are depicted in red.

In addition, the usage of aromatic linkers close to the hydroxamate moiety is a feature that favors HDAC6 inhibition [67]. Indeed HDAC6, selectivity can be increased by incorporating bulkier capping groups, such as our aniline-purine scaffold, that would fit in the wider rim of the enzyme [67]. The introduction of voluminous groups was also validated as a strategy to get selective inhibitors for AKT2, due to its enlarged subsite [68], which was favored by the dual HDAC/kinase design. So far, no hybrid AKT/HDAC inhibitor has been described, but only compounds that indirectly reduce its activity [10]. Considering the impact of its activation in different cell signaling pathways, the optimization of **6k** as a potent and selective AKT2/HDAC6 hybrid inhibitor could be promising for cancer treatment.

### Conclusions

Using machine learning, 13 cancer cell lines and 48 QSAR models were built for the putative targets of kinase/HDAC hybrid inhibitors. These hybrids were subjected to the LBVS, and a set of representative compounds with promising profiles were synthesized and subjected to biological evaluation. It can be noted that models K562, MV4-11, MM1S, NB-4, U937, and HDAC6 models presented correct classification rates of 62.5%, 75.0%, 62.5%, 77.8%, and 86.0% in biological testing, indicating these models are acceptable for further usage as prediction tools in drug design programs. Besides that, this study aided the identification of **6b** and **6k** as potent anticancer inhibitors with IC_50_ of 0.2-0.8 µM in three cancer cell lines, linked to HDAC6 inhibition below 2 nM and blockade of AKT phosphorylation at 2 μM, whose selectivity is favored by its bulky linker and nitro-substituent, as future candidates for optimization for the treatment of hematological malignancies.

## EXPERIMENTAL SECTION

### Cancer cell models and target selection

A set of 13 hematological cancer cell lines were selected to provide biological diversity of cancer models (**Supplementary Table S1**), which could be associated with different pathology mechanisms, and present super expression of different biological targets [30]. The hybrid kinase/HDAC inhibitors were drawn in MarvinSketch22.4 and saved as SDF and SMILES formats. The SDF files were uploaded into PharmMapper (http://www.lilab-ecust.cn/pharmmapper/), and the option Human Protein Targets Only (v2010, 2241) was selected while leaving the other parameters as default [35]. The compounds in SMILES format were uploaded into SwissTargetPrediction (http://www.swisstargetprediction.ch/), and the *Homo sapiens* species was selected for target fishing [36]. The targets retrieved from both tools were matched and filtered for kinase proteins. Two filtering approaches were conducted. In the first one, all protein targets were matched and then the data was filtered for kinases, and in the second one the filter for kinases was conducted first, and then only the kinase targets were matched. Only kinases that appeared as consensus in both methods were considered selected for the analysis. Twelve kinase targets linked to hematological malignancies and HDACs 1-10 were also chosen based on the biological test results of similar hybrid compounds in these cancer cell types [29].

### Dataset selection and preparation

Following target selection, a set of chemical structures with known activity against each enzyme was downloaded from the ChEMBL database (https://www.ebi.ac.uk/chembl/) [69] to construct the dataset. All molecular structures were subjected to chemical curation as previously described by Fourches *et al.* [70] and were standardized to canonical representations using Python and RDKit standardizer in KNIME 4.6 Software (Knime 4.6 the Konstanz Information Miner Copyright, 2003–2020, www.knime.org). To increase reproducibility, cancer cell models were subjected to a second round of curation, to select only IC_50_ data determined by MTT assays conducted between 0-72h. Assay descriptions with the following keywords were removed during the curation: *harboring, deficient, treated* (with an additional drug besides the one being tested)*, sensitive, atplite, glo-assay, blue, knockout, srb, hypoxia-resistant, expressing (*for genes that are not constitutively expressed in the cell line*), negative, absence, presence, dark, washout, mutant, mutated, photo*.

The molecules in each dataset were classified as active or inactive based on their IC_50_ values for the target. The activity cutoffs for each were chosen considering micromolar potency (set between 1 - 2000 nM for enzymatic models and between 0.05 – 2.0 µM for cancer cell models) and balancing between both active and inactive sets, to increase chemical diversity.

### QSAR modeling and validation

The QSAR models were developed and validated according to the best practices described in the literature [40]. All the analysis and *in silico* model generation were performed in KNIME 4.6 Software (Knime 4.6 the Konstanz Information Miner Copyright, 2003–2020, www.knime.org). Chemical descriptors were calculated employing the Morgan algorithm to generate ECFP4-like circular fingerprints with 2048 bits through “RDKit Fingerprint” and “Fingerprints Expander” nodes. Each dataset was divided using the “Partitioning” tool with the “Stratified Sample” option to create a training set with 80% of the compounds, and an external set with the remaining 20% molecules, while maintaining the same proportion of active and inactive samples in both. The fragment contribution maps were generated according to the described in the literature [71] using physicochemical and ECFP4-like circular fingerprints with 2048 bits. The models for AKT, CK1α, CSK, EPHA2, HDAC1, HDAC6, HDAC7, JAK3, PI3Kδ, and TRKα were built in Phyton 3 using the described dataset for each target (**Supplementary Table S3**).

The following 5-fold external cross-validation procedure was employed: the full set of compounds with known experimental activity was randomly divided into five subsets of equal size with the Random Forest algorithm (RF); then one of these subsets (20% of all compounds) is set aside as test set and the remaining four sets together form the modeling set (80% of the full set). This procedure was repeated five times allowing each of the five subsets to be used as a test set. The external performance of the selected models was assessed for accuracy through correct classification rate (CCR), sensitivity (true-positive rate), and specificity (true-negative rate). The positive (PPV) and negative predictive values (NPV) indicate the probability of predicted positives and negatives being true positives and negatives, respectively. The applicability domain (AD) was estimated as defined by Tropsha and Golbraikh [72].

### Ligand-Based Virtual screening

The hybrid kinase/HDAC inhibitors were written in SMILES format and virtual screening was performed using the classification models created with the RF algorithm.

### Chemistry

Solvents were purified according to standard procedures. Reagents and solvents were purchased from Synth®, Merck®, Sigma-Aldrich-Merck®, and Oakwood Chemicals®. Reactions were monitored by TLC on Merck silica gel (60 F 254) by using UV light (λ = 254 nm) and iodine as visualizing agents and ninhydrin and vanillin staining solutions. The compounds were purified by precipitation, recrystallization, or column chromatography with silica gel (pore size: 60Å). ^1^H and ^13^C NMR spectra were obtained on a 300/75 MHz Bruker spectrometer, using the solvent residual peak as the internal reference (chemical shifts: DMSO[Z]d_6_, 2.50/39.52). Analytical HPLC was carried out on a Shimadzu™ Proeminence® instrument under the following conditions: column, C-18 Gemini® (5 μm, 150 × 4.6 mm); mobile phase, 5–100 % H_2_O/CH_3_CN containing 0.1 % TFA at a flow rate of 1.0 mL/min for 25 min; UV detection at 254 nm. The purities of all tested compounds were >95 %, as determined by analytical HPLC.

### General Procedure A

In a 25 mL round-bottom flask, 5 mmol of 6-chloro-9*H*-purine (772.8 mg, 5 mmol, 1.0 eq.), and 5 mmol of the aniline (5 mmol, 1.0 eq.) were added, along with 10 mL of isopropanol (boiling point 82.3 °C). To that suspension, 4 mmol of concentrated HCl (0.4 mL, 0.8 eq.) was added, and the system was kept under agitation and reflux for 18 h. After the end of the reaction, the reaction mixture was filtered under a vacuum and washed with isopropanol (3 x 30 mL). The filtered solid was resuspended in 50 mL of NaHCO_3_ solution and stirred for 10 min. After that, the suspension was filtered under a vacuum, washed in H_2_O (3 x 30 mL), and dried under a vacuum to obtain product **2a-d**, and **2k** as solids.

### *N*-(4-methoxyphenyl)-9*H*-purin-6-amine (2a)

The product was obtained from 4-methoxyaniline (615.8 mg, 5 mmol) as a green solid with 73% yield (880.6 mg). ^1^H NMR (300 MHz, DMSO-*d*_6_), 13.06 (bs, 1H), 9.56 (s, 1H), 8.30 (s, 1H), 8.21 (s, 1H), 7.80 (d, *J* = 8.2 Hz, 2 H), 6.92 (d, *J* = 8.3 Hz, 2 H), 3.74 (s, 3H). ^13^C NMR (75 MHz, DMSO-*d*_6_), δ ^13^C RMN (75 MHz, DMSO-*d*_6_), δ ^13^C RMN (75 MHz, DMSO-*d*_6_), δ 154.9, 152.0, 151.8, 150.1, 139.4, 132.7, 122.4 (2C), 119.1, 113.5 (2C), 55.1.

### *N*-(4-chlorophenyl)-9*H*-purin-6-amine (2b)

The product was obtained from 4-chloroaniline (637.8 mg, 5 mmol) as a green solid with 71% yield (872.1mg). ^1^H NMR (300 MHz, DMSO-*d*_6_), δ 13.18 (bs, 1H), 9.89 (s, 1H), 8.39 (s, 1H), 8.29 (s, 1H), 8.02 (d, *J* = 8.2 Hz, 2H), 7.37 (d, *J* = 8.7 Hz, 2H). ^13^C NMR (75 MHz, DMSO-*d*_6_), δ 151.6 (2C), 150.5, 140.1, 138.9, 128.1 (2C), 125.8, 121.8 (2C), 119.5.

### *N*-(4-bromo-2-fluorophenyl)-9*H*-purin-6-amine (2c)

The product was obtained from 4-bromo-2-fluoroaniline (950.1 mg, 5 mmol), as a green solid with 84% yield (1294.1 mg). ^1^H NMR (300 MHz, DMSO-*d*_6_) 13.14 (bs, 1 H), 9.36 (s, 1 H), 8.28-8.27 (m, 2H), 7.70 (bs, 1H), 7.59 (d, *J* = 9.2 Hz, 1H), 7.41 (d, *J* = 8.7 Hz,1 H). ^13^C NMR (75 MHz, DMSO-*d*_6_): δ 155.5 (d, *J^1^* = 249.0 Hz, C-F), 151.7 (2C), 151.3, 140.8, 127.6, 127.2 (d, *J^4^* = 3.6 Hz, C-F), 126.5 (d, *J^3^* = 12.0 Hz, C-F), 118.3 (d, *J^2^*= 23.3 Hz, C-F), 116.1 (d, *J^3’^* = 9.0 Hz, C-F).

### *N*-(3-chloro-4-fluorophenyl)-9*H*-purin-6-amine (2d)

The product was obtained from 3-chloro-4-fluoroaniline (727.8 mg, 5 mmol), as a grey solid with 86% yield (1133.7 mg). ^1^H NMR (300 MHz, DMSO-*d*_6_) δ 9.87 (bs, 1H), 8.36 (s, 1H), 8.35-8.33 (m, 1H), 8.23 (s, 1H), 7.92-7.89 (m, 1 H), 7.38-7.32 (m, 1 H). ^13^C NMR (75 MHz, DMSO-*d*_6_), δ 152.36 (d, *J^1^* = 240.0 Hz, C-F), 152.5, 150.99, 150.91, 142.0, 137.5 (d, *J^4^* = 3.0 Hz, C-F), 121.2 (d, *J^3’^* = 6.8 Hz, C-F), 119.3, 118.6 (d, *J^2^* = 18.8 Hz, C-F), 116.4 (d, *J^2’^*= 21.0 Hz, C-F).

### *N*-(3-nitrophenyl)-9*H*-purin-6-amine (2k)

The product was obtained from 3-nitroaniline (690.65 mg, 5 mmol), as a green solid with 93% yield (1191.5 mg). ^1^H NMR (300 MHz, DMSO-*d*_6_) δ 13.28 (s, 1H), 10.32 (s, 1H), 9.10 (s, 1H), 8.48 (s, 1H), 8.37 (d, *J =* 8.4 Hz, 1H), 8.34 (s, 1H), 7.86-7.82 (m,1 H), 7.62-7.56 (t, *J* = 8.3 Hz, 1H).^13^C NMR (75 MHz, DMSO-*d*_6_), δ 151.5, 151.3, 150.8, 147.9, 141.2, 140.5, 129.6, 126.1, 119.7, 116.4, 114.1.

### General Procedure B

In a 50 mL round-bottom flask, 2 mmol of the intermediate **2a-d** or **2k** (2.0 mmol, 1.0 eq.), 2 mmol of methyl 4-(bromomethyl)benzoate (458.1 mg, 2.0 mmol, 1.0 eq.), 6 mmol of K_2_CO_3_ (829.2 mg, 6.0 mmol, 3 eq.), and 20 mL of acetone were added. The mixture was kept under agitation and reflux for 22h. The reaction was interrupted by adding 20 mL of H_2_O and the acetone was evaporated. Subsequently, the mixture was extracted with EtOAc (3 x 20 mL), and the combined organic phase was washed with H_2_O, and brine solution, and dried with Na_2_SO_4_. The solvent was evaporated, forming a solid that was purified by column chromatography with a gradient of elution of ethyl acetate-methanol (EtOAc-MeOH 99:1 to EtOAc-MeOH 95:5, 50 mL fractions), obtaining **3a-d** or **3k** as solids.

### Methyl 4-((6-((4-methoxyphenyl)amino)-*9H*-purin-9-yl)methyl)benzoate (3a)

The product was obtained from intermediate **2a** (482.5 mg, 2 mmol), as a white solid with 59% yield (459.5 mg). ^1^H NMR (300 MHz, DMSO-*d*_6_), δ 9.71 (s, 1H), 8.40 (s, 1H), 8.32 (s, 1H), 7.93 (d, *J* = 7.7 Hz, 2H), 7.77 (d, *J* = 8.3 Hz, 2H), 7.42 (d, *J* = 7.8 Hz, 2H), 6.90 (d, *J* = 8.4 Hz, 2H), 5.52 (s, 2H), 3.83 (s, 3H), 3.74 (s, 3H). ^13^C NMR (75 MHz, DMSO-*d*_6_), δ 165.8, 155.0, 152.2, 152.1, 149.4, 141.2, 141.4, 132.5 (2C), 129.5 (2C), 128.9, 127.6 (2C), 122.6 (2C), 119.4, 113.5 (2C), 55.1, 52.0, 45.9.

### Methyl 4-((6-((4-chlorophenyl)amino)-9*H*-purin-9-yl)methyl)benzoate (3b)

The product was obtained from intermediate **2b** (491.3 mg, 2 mmol), as a white solid with 60% yield (472.6 mg). ^1^H NMR (300 MHz, DMSO-*d*_6_), δ 10.05 (s, 1H), 8.47 (s, 1H), 8.41 (s, 1H), 8.01 (d, *J* = 7.9, 2H), 7.93 (d, *J =* 7.2 Hz, 2H), 7.43 (d, *J* = 7.4 Hz, 2H), 7.37 (d, *J =* 8.1 Hz, 2H) 5.54 (s, 2H), 3.82 (s, 3H). ^13^C NMR (75 MHz, DMSO-*d*_6_ δ 165.8, 152.0, 151.7, 149.7, 142.0, 141.9, 138.6, 129.5 (2C), 129.0, 128.1 (2C), 127.6 (2C), 126.1, 122.0 (2C), 119.7, 52.0, 46.0.

### Methyl 4-((6-((4-bromo-2-fluorophenyl)amino)-9*H*-purin-9-yl)methyl)benzoate (3c)

The product was obtained from intermediate **2c** (616.2 mg, 2 mmol), as a white solid with 60% yield (547.5 mg). ^1^H NMR (300 MHz, DMSO-*d*_6_), δ 9.58 (s, 1H), 8.44 (s, 1H), 8.29 (s, 1H), 7.93 (d, *J* = 7.5 Hz, 2H), 7.66-7.57 (m, 2H), 7.43-7.41 (m, 3H), 5.52 (s, 2H), 3.82 (s, 3H). ^13^C NMR (75 MHz, DMSO-*d*_6_), δ 165.8, 156.2 (d, *J^1^* = 249.8 Hz, C-F), 152.2, 152.1, 150.0, 142.2, 142.0, 129.5 (2C), 129.0, 128.5, 127.6 (2C), 127.3 (d, *J^4^* = 3.0 Hz, C-F), 126.3 (d, *J^3^* = 12.0 Hz, C-F), 119.6, 119.1 (d, *J2* = 23.3 Hz, C-F), 116.9 (d, *J* = 9.0 Hz, C-F), 52.1, 46.0.

### Methyl 4-((6-((3-chloro-4-fluorophenyl)amino)-9*H*-purin-9-yl)methyl)benzoate (3d)

The product was obtained from intermediate **2d** (527.3 mg, 2 mmol), as a gray solid with 37% yield (304.7 mg). ^1^H NMR (300 MHz, DMSO-*d*_6_), δ 10.12 (s, 1H), 8.48 (s, 1H), 8.44 (s, 1H), 8.320 (dd, *J* = 6.8 Hz, 2.5 Hz, 1H), 7.99 (d, J = 8.1 Hz, 2H), 7,93 (d, *J* = 8.2 Hz, 2H), 7.89-7.88 (m, 1H), 7.44 (d, *J* = 8.2 Hz, 2H), 7.40-7.33 (m, 1H), 5.54 (s, 2H), 3.82 (s, 3H). ^13^C NMR (75 MHz, DMSO-*d*_6_), δ 165.8, 152.7 (d, *J^1^* = 240.8 Hz, C-F), 152.0, 151.6, 149.7, 142.0, 136.9 (d, *J^4^* = 1.5 Hz, C-F), 129.5 (2C), 129.0, 127.6 (2C), 121.7, 120.7 (d, *J^3’^* = 6.8 Hz, C-F), 119.7, 118.6 (d, *J^2^* = 18.0 Hz, C-F), 116.4 (d,*’J^2’^* = 21.0 Hz, C-F), 52.0, 46.0.

### Methyl 4-((6-((3-nitrophenyl)amino)-9*H*-purin-9-yl)methyl)benzoate (3k)

The product was obtained from intermediate **2k** (512.5 mg, 2 mmol), as a yellow solid with 49% yield (396.3 mg). ^1^H NMR (300 MHz, DMSO-*d*_6_), δ 10.34 (s, 1H), 9.05 (s, 1H), 8.51 (s, 1H), 8.50 (s, 1H), 8.37 (d, *J* = 8.0 Hz, 1H), 7.93 (d, *J* = 7.8 Hz, 2H), 7.86 (d, *J* = 8.1 Hz, 1H), 7.642-7.57 (m, 1 H), 7.46 (d, *J* = 7.8 Hz, 2H), 5.57 (s, 2H), 3.83 (s, 3H). ^13^C NMR (75 MHz, DMSO-*d*_6_), δ 165.9, 152.0, 151.6, 150.0, 148.0, 142.5, 142.0, 141.1, 129.7, 129.6 (2C), 129.1, 127.7 (2C), 126.4, 120.0, 116.8, 114.4.

### General Procedure C

In a 25 mL round-bottom flask, 1 mmol of the esters **3b-c** (1 mmol, 1 eq.) were dissolved in 3 mL of dioxane. To that, 3.3 mL of a solution of KOH 2M was added (3.3 mL KOH 2M/mmol of ester, 6.6 eq.). The system was placed under agitation and reflux (100 °C). After the reaction was concluded, the system was cooled at room temperature, and placed in an ice bath. The pH was then adjusted to 4.0, with HCl 2 M. The suspension was then transferred to a beaker and stirred for 10 min with 5 mL of H_2_O. The solution was cooled in an ice bath, filtered under a vacuum, and washed with cold H_2_O (3 x 5 mL), and ethyl ether (3 x 5 mL). The solid was dried under vacuum to obtain products **4a-c** as solids.

### 4-((6-((4-methoxyphenyl)amino)-9H-purin-9-yl)methyl)benzoic acid (4a)

The product was obtained from intermediate **3a** (389.4 mg, 1 mmol), as a white solid with 98% yield (367.9 mg). ^1^H NMR (300 MHz, DMSO-*d*_6_), δ 9.71 (s, 1H), 8.40 (s, 1H), 8.32 (s, 1H), 7.92 (d, *J* = 7.7 Hz, 2H), 7.78 (d, *J* = 8.3 Hz, 2H), 7.40 (d, *J =* 7.4 Hz, 2H), 6.91 (d, *J =* 8.3 Hz, 2H), 5.51 (s, 2H), 3.73 (s, 3H). ^13^C NMR (75 MHz, DMSO-*d*_6_), δ 166.9, 155.1, 152.2, 152.1, 149.4, 141.6, 141.4, 132.5, 130.3, 129.6 (2C), 127.4 (2C), 122.7 (2C), 119.4, 113.5 (2C), 55.1, 45.9.

### 4-((6-((4-chlorophenyl)amino)-9H-purin-9-yl)methyl)benzoic acid (4b)

The product was obtained from intermediate **3b** (393.8 mg, 1 mmol), as an orange solid with 88% yield (334.2 mg). ^1^H NMR (300 MHz, DMSO-*d*_6_), δ 12.91 (bs, 1H), 10.04 (s, 1H), 8.47 (s, 1H), 8.42 (s, 1H), 8.01 (d, J = 8.8 Hz, 2H), 7.92 (d, J = 8.0 Hz, 2H), 7.41 (d, J = 8.3 Hz, 2H), 7.37 (d, J = 8.8 Hz, 2H), 5.53 (s, 2H). ^13^C NMR (75 MHz, DMSO-*d*_6_), δ 166.9, 152.0, 151.7, 149.7, 142.0, 141.6, 138.7, 130.2, 129.7 (2C), 128.2 (2C), 127.5 (2C), 126.1, 122.1 (2C), 119.8, 46.0.

### 4-((6-((4-bromo-2-fluorophenyl)amino)-9H-purin-9-yl)methyl)benzoic acid (4c)

The product was obtained from intermediate **3c** (456.3 mg, 1 mmol), as a yellow solid with 82% yield (362.6 mg). ^1^H NMR (300 MHz, DMSO-*d*_6_), δ 9.58 (bs, 1H), 8.44 (s, 1H), 8.29 (s, 1H), 7.91 (d, *J* = 8.0 Hz, 2H), 7.66-7.57 (m, 2H), 7.41-7,39 (m, 3 H), 5.52 (s, 2H). ^13^C NMR (75 MHz, DMSO-*d*_6_), δ 166.9, 156.2 (d, *J^1^* = 249.8 Hz, C-F), 152.2, 152.1, 150.0, 142.2, 141.5, 130.4, 129.7 (2C), 128.5, 127.4 (2C), 127.2 (d, *J^4^* = 3.8 Hz, C-F), 126.3 (d, *J^3^* = 12.0 Hz, C-F), 119.5, 119.1 (d, J^2^ = 23.3 Hz, C-F), 116.9 (d, *J^3’^* = 9.0 Hz, C-F), 46.0.

### General Procedure D

In a 10 mL round-bottom flask, 0.5 mmol of the carboxylic acid intermediate **4b-c** (0.5 mmol, 1 eq.), 1 mmol of EDC (191.7 mg, 1 mmol, 2 eq.), and 0.7 mmol of HOBt (114.8 mg, 0.7 mmol, 1.5 eq.) were added. Then, the reagents were dissolved in 5 mL of DMF, cooled to 0°C, and 1.4 mmol of triethylamine (0.2 mL, 1.4 mmol, 2.5 eq.) was added. The solution was maintained at this temperature for 10 min and at room temperature for 1 hour. After this period, 0.5 mmol of 1,2-diaminebenzene (54.1 mg, 0.5 mmol, 1 eq.) was added, and the reaction mixture was stirred for 18 hours or until the starting materials were consumed. The reaction was quenched with water and extracted with EtOAc (50 mL x 3). The combined organic extracts were washed with brine, dried with Na_2_SO_4_, and concentrated. The desired product was isolated by column chromatography in a gradient of elution of ethyl acetate-methanol (EtOAc-MeOH 99:1 to EtOAc-MeOH 95:5, with 1% acetic acid in 50 mL fractions), yielding **5b-c** as solids.

### N-(2-aminophenyl)-4-((6-((4-methoxyphenyl)amino)-9H-purin-9-yl)methyl)benzamide (5a)

The final product was obtained from intermediate **4a** (187.7 mg, 0.5 mmol), as an orange solid with 14% yield (32.6 mg). ^1^H NMR (300 MHz, DMSO-*d*_6_), δ 9.70 (s, 1H), 8.62 (s, 1H), 8.43 (s, 1H), 8.33 (s, 1H), 7.95 (d, *J* = 8.0 Hz, 2H), 7.77 (d, *J* = 8.9 Hz, 2H), 7.44 (d, *J =* 8.1 Hz, 2H), 7.15 (d, *J =* 7.8 Hz, 1H), 6.95-6-89 (m, 3H),, 6.76 (d, *J* = 7.1 Hz, 1H), 6.58 (t, *J* = 7.1 Hz, 1H), 5.52 (s, 2H) 3.74 (s, 3H). ^13^C NMR (75 MHz, DMSO-*d*_6_), δ 165.1, 155.2, 152.1 (2C), 149.4, 141.4, 140.3, 133.8, 132.4, 128.2 (2C), 127.4 (2C), 126.7, 126.5, 125.1, 122.8 (2C), 119.4, 119.0, 117.9, 113.6 (2C), 55.2, 46.1. HRMS (ESI) *m/z*: [M + H]^+^ Calcd for C_26_H_23_N_7_O_2_ 466.1991; Found 466.1989. Purity: 97.5% (254 nm).

### *N*-(2-aminophenyl)-4-((6-((4-chlorophenyl)amino)-9*H*-purin-9-yl)methyl)benzamide (5b)

The final product was obtained from intermediate **4b** (189.9 mg, 0.5 mmol), as an orange solid with 37% yield (86.9 mg). ^1^H NMR (300 MHz, DMSO-*d*_6_), δ 10.04 (s, 1H), 9.59 (s, 1H), 8.49 (s, 1H), 8.43 (s, 1H), 8.01 (d, *J* = 8.1 Hz, 2H), 7.94 (d, *J =* 7.6 Hz, 2H), 7.45 (d, *J* = 7.7 Hz, 2H), 7.37 (d, *J =* 8.3 Hz, 2H), 7.14 (d, *J* = 7.4 Hz, 1H), 6.95 (t, *J* = 7.3 Hz, 1H), 6.76 (d, *J* = 7.9 Hz, 1H), 6.58 (t, *J =* 7.4 Hz, 1H), 5.53 (s, 2H), 4.85 (s, 2H). ^13^C NMR (75 MHz, DMSO-*d*_6_) δ 164.9, 152.0, 151.7, 149.7, 143.0, 142.0, 140.0, 138.7, 134.1,128.2 (2C), 128.1 (2C), 127.4 (2C), 126.6, 126.5, 126.1, 123.1, 122.1 (2C), 119.8, 116.2, 116.0, 46.1. HRMS (ESI) *m/z*: [M + H]^+^ Calcd for C_25_H_20_ClN_7_O 470.1496; Found 470.1492. Purity: 95.9% (254 nm).

### *N*-(2-aminophenyl)-4-((6-((4-bromo-2-fluorophenyl)amino)-9*H*-purin-9-yl)methyl)benzamide (5c)

The final product was obtained from intermediate **4c** (221.1 mg, 0.5 mmol), as a white solid with 33% yield (34.6 mg). ^1^H NMR (300 MHz, DMSO-*d*_6_), δ 9.58-9.57 (m, 2H), 8.46 (s, 1H), 8.31 (s, 1H), 7.94 (d, *J* = 7.6, 2H), 7.66-7.58 (m, 2H), 7.45-7.40 (m, 3H), 7.15 (d, *J* = 7.5 Hz, 1H), 6.95 (t, *J* = 7.4 Hz, 1H), 5.52 (s, 2H), 4.85 (s, 2H). ^13^C NMR (75 MHz, DMSO-*d*_6_), δ 164.9, 156.2 (d, *J^1^* = 250.5 Hz, C-F), 152.2, 152.1, 150.0, 143.0, 142.1, 139.9, 134.1, 128.4, 128.1 (2C), 127.3 (2C), 127.2 (d, *J^4^* = 3.8 Hz, C-F), 126.5, 126.4, 126.2 (d, *J^3^* = 11.5 Hz, C-F), 123.1, 119.6, 119.0 (d, *J^2’^* = 23.3 Hz, C-F), 116.8 (d, *J^3’^* = 9 Hz, C-F), 116.1, 116.0, 46.1. HRMS (ESI) *m/z*: [M + H]^+^ Calcd for C_25_H_19_BrFN7O 532.0896; Found 532.0890. Purity: 95.4% (254 nm).

### General Procedure E

A 10 mL round-bottom flask containing 4.0 mmol of NaOH (159.9 mg, 8.0 eq.) was placed in an ice bath. When the temperature achieved 0° C, 25 mmol of hydroxylamine in an aqueous solution (NH_2_OH 50% p/v, 1.7 mL, 25.0 mmol) was added to dissolve the NaOH. To that, a solution containing 0.5 mmol intermediate **3** (0.5 mmol, 1 eq.) dissolved in 3 mL tetrahydrofuran, methanol, and dioxane (THF:MeOH:dioxane 1:1:1) was added dropwise. The mixture was kept under agitation at room temperature for 2 h. After the reaction was completed, the solution was moved to a 50 mL flask, placed in an ice bath, and neutralized to pH 7.0. To that, 20 ml of water was added, forming a precipitate. The resulting solution was filtered under a vacuum, washed with cold H_2_O (3 x 30 mL), and dried under a vacuum to obtain product **6a-d**, and **6k** as solids.

### *N*-hydroxy-4-((6-((4-methoxyphenyl)amino)-*9H*-purin-9-yl)methyl)benzamide (6a)

The final product was obtained from intermediate **3a** (194.7 mg, 0.5 mmol), as a white solid with 72% yield (140.5 mg). ^1^H NMR (300 MHz, DMSO-*d*_6_), δ 11.14 (bs, 1H), 9.70 (s, 1H), 9.04 (bs, 1H), 8.41 (s, 1H), 8.32 (s, 1H), 7.78 (d, *J* = 8.2 Hz, 2H), 7.72 (d, *J* = 7.6 Hz, 2H), 7.38 (d, *J =* 7.5 Hz, 2H), 6.91 (d, *J =* 8.2 Hz, 2H), 5.47 (s, 2H), 3.74 (s, 3H). ^13^C NMR (75 MHz, DMSO-*d*_6_), δ 163.7, 155.1, 152.2, 152.1, 149.4, 141.4, 139.9, 132.5, 132.2, 127.4 (2C), 127.2 (2C), 122.7 (2C), 119.4, 113.5 (2C), 55.1, 45.9. Purity: 99.9% (254 nm).

### 4-((6-((4-chlorophenyl)amino)-9H-purin-9-yl)methyl)-N-hydroxybenzamide (6b)

The final product was obtained from intermediate **3b** (196.9 mg, 0.5 mmol), as a white solid with 83% yield (163.9 mg). ^1^H NMR (300 MHz, DMSO-*d*_6_), δ 11.16 (bs, 1 H), 10.03 (s, 1H), 9.02 (bs, 1H), 8.47 (s, 1H), 8.42 (s, 1H), 8.01 (d, *J* = 7.1, 2H), 7.72 (d, *J* = 6.9 Hz, 2H), 7.38-7.37 (m, 4H), 5.50 (s, 2H). ^13^C NMR (75 MHz, DMSO-*d*_6_), δ 163.8, 152.0, 151.7, 149.7, 141.9, 139.7, 138.7, 132.3, 128.2 (2C), 127.4 (2C), 127.2 (2C), 126.1, 122.1 (2C), 119.7, 46.0. HRMS (ESI) *m/z*: [M + H]^+^ Calcd for C_19_H_15_ClN_6_O_2_ 395.1023; Found 395.1019. Purity: 99.1% (254 nm).

### 4-((6-((4-bromo-2-fluorophenyl)amino)-9H-purin-9-yl)methyl)-N-hydroxybenzamide (6c)

The final product was obtained from intermediate **3c** (228.14 mg, 0.5 mmol), as a white solid with 94% yield (214.9 mg). ^1^H NMR (300 MHz, DMSO-*d*_6_), δ 11.15 (s, 1H), 9.56 (s, 1H), 8.99 (s, 1H), 8.44 (s, 1H), 8.29 (s, 1H), 7.71 (d, *J* = 7.3, 2H), 7.65 (m, 2H), 7.43-7.36 (m, 3H) 5.48 (s, 2H). ^13^C NMR (75 MHz, DMSO-*d*_6_), δ 163.8, 156.2 (d, *J^1^* = 250.5 Hz, C-F), 152.2, 152.1, 150.0, 142.1, 139.7, 132.3, 128.4, 127.4 (2C), 127.2 (3C), 126.8 (d, *J^4^* = 3.75 Hz, C-F), 126.2 (d, *J^3^* = 11.25 Hz, C-F), 119.6, 119.0 (d, *J^2^* = 24 Hz, C-F), 116.8 (d, *J^3’^* = 9.0 Hz, C-F), 46.0. Purity: 98.7% (254 nm).

### 4-((6-((3-chloro-4-fluorophenyl)amino)-9H-purin-9-yl)methyl)-N-hydroxybenzamide (6d)

The final product was obtained from intermediate **3d** (205.9 mg, 0.5 mmol), as a white solid with 70% yield (144.5 mg). ^1^H NMR (300 MHz, DMSO-*d*_6_), δ 11.62 (s, 1H), 10.12 (s, 1H), 8.99 (s, 1H), 8.48 (s, 1H), 8.44 (s, 1H), 8.32-8.30 (m, 1H), 7.92-7.89 (m,1H), 7.72 (d, *J* = 7.7 Hz, 2H), 7.39 (d, *J =* 8.0 Hz, 2H), 7.34 (m, 1H), 5.50 (s, 2H). ^13^C NMR (75 MHz, DMSO-*d*_6_), δ 163.8, 152.7 (d, *J^1^* = 240 Hz, C-F), 151.9 (), 151.6, 149.7, 142.1, 139.7, 137.0 (d, *J^4^* = 3.0 Hz, C-F), 132.3, 127.4 (3C), 127.2 (2C), 121.7, 120.8 (d, *J^3^* = 6.0 Hz, C-F), 119.7, 118.7 (d, *J^2^* = 18.8 Hz, C-F), 116.5 (d, *J^2’^* = 21.0 Hz, C-F), 46.0. Purity: 95.1% (254 nm).

### *N*-hydroxy-4-((6-((3-nitrophenyl)amino)-9*H*-purin-9-yl)methyl)benzamide (6k)

The final product was obtained from intermediate **3k** (202.2 mg, 0.5 mmol), as a yellow solid with 51% yield (103.4 mg). ^1^H NMR (300 MHz, DMSO-*d*_6_), δ 11.16 (bs, 1H), 10.44 (s, 1H), 9.08 (s, 1H), 9.01 (bs, 1 H), 8.52 (s, 2H), 8.36 (d, *J* = 7.5 Hz, 1H), 7.87 (d, *J* = 7.6 Hz, 1H), 7.72 (d, *J* = 6.5 Hz, 2H), 7.62-7.57 (m, 1H), 7.40 (d, *J* = 6.8 Hz, 2H), 5.52 (s, 2H). ^13^C NMR (75 MHz, DMSO-*d*_6_), δ 163.7, 151.9, 151.5, 149.9, 147.9, 142.4, 141.0, 139.7, 132.3, 129.6, 127.4 (2C), 127.3 (2C), 126.3, 120.0, 116.7, 114.3. HRMS (ESI) *m/z*: [M + H]^+^ Calcd for C_19_H_15_N_7_O_4_ 406.1263; Found 4061256. Purity: 100% (254 nm).

### Cell culture

HEL (AML), Jurkat (ALL), K562 (CML) U266 (MM), U937 (AML) and MM1S (MM), MOLT-4 (ALL) were kindly provided by Prof. Dr. Sara Teresinha Olalla Saad (Hemocentro, State University of Campinas). HL-60 (AML), MV4-11 (AML), NALM-6 (ALL), NB-4 (APL), Raji (ALL), THP1 (AML) were kindly provided by Dr. Gilberto Carlos Franchi Junior (Universidade Estadual de Campinas, Campinas, Brazil). The cell lines were cultivated in a culture medium indicated by the American Type Culture Collection (ATCC) or Deutsche Sammlung von Mikroorganismen und Zellkulturen (DSMZ), supplemented with fetal bovine serum and penicillin/streptomycin. The cells will be maintained at 37°C, 5% CO2. RUXOLITINIB were obtained from Target Mol (Target Mol, Boston, MA, USA). Vorinostat (SAHA) were obtained from Sigma-Aldrich (Sigma-Aldrich, St. Louis, MO, USA). Dimethyl sulfoxide (DMSO) (Synth, Diadema, SP, Brazil) was used as a vehicle for the dilution of the tested compounds.

### Cell viability evaluation by MTT assay

The cytotoxicity of the compounds was measured by methylthiazol tetrazolium (MTT) assay [73] (Thermo Fisher Scientific, San Jose, CA, USA). HEL, HL-60, Jurkat, K562, MM1S, MOLT-4, MV4-11, NALM-6, NB-4, Raji, THP1, U266, U937 (2 × 10^4^ cells/well), were cultured in a 96-well plate in appropriate medium in the presence of vehicle or increased concentrations (0.0032, 0.016, 0.08, 0.4, 2, 10 and 50 μM) of vorinostat, venetoclax and tested compounds for 72 h. At least three independent experiments were to be performed for each condition. After treatment, 10 μL/well of MTT solution was added (0.5 mg/mL final concentration) and incubated for an additional 3 h. After MTT incubation, 100 μL of 0.1 N HCl in isopropanol solution was added per well-stopping reaction. IC_50_ values and their 95% confidence intervals were calculated using sigmoidal nonlinear regression analysis performed with GraphPad Prism 5 (GraphPad Software, Inc., San Diego, CA, USA).

### IC_50_ determination in HDACs

HDAC inhibition assays were performed using the recombinant proteins purchased from ENZO Life Sciences AG (Lausen, Switzerland) for HDAC1, 2, 3, and 6, whereas HDAC4, 5, 7, and 9 were produced as described in previous work [56]. All inhibitors were tested in an enzymatic *in vitro* assay, as described previously, using 384-well plates (GreinerONe, catalogue no. 784900) [56,74]. After five minutes of incubation of inhibitors with the respective enzyme (HDAC1: 10 nM, HDAC2 and 3: 3 nM, HDAC4: 5 nM, HDAC5: 10 nM, HDAC6: 1 nM, HDAC7: 5 nM, HDAC8: 2 nM, HDAC 9: 20 nM, HDAC10: 5 nM), the reactions were started by the addition of the substrate.

For HDAC1, 2, 3, and 6, an acetylated peptide substrate derived from p53 (Ac-RHKK(Acetyl)-AMC) was used in a discontinuous fluorescence assay, as described previously [74]. All reactions were performed in assay buffer (20 mM HEPES, 140 mM NaCl, 10 mM MgCl_2_, 1 mM TCEP, and 0.2 mg/mL BSA, pH 7.4 adjusted with NaOH) at room temperature for HDACs 2, 3 and 6, and at 37 °C for HDAC1. After 45 min for HDAC1, 2 and 3 or after 90 min for HDAC1, the reaction was quenched by adding trypsin and SAHA. The fluorescence intensity was measured after 1 h of incubation using an Envision 2104 Multilabel Plate Reader (PerkinElmer, Waltham, MA, USA) with an excitation wavelength of 380 ± 8 nm and an emission wavelength of 430 ± 8 nm.

HDAC4–7, 8, and 9 were measured in a continuous manner using the thioacetylated peptide substrate (Abz-SRGGK(thio-TFA)FFRR-NH2) which was described previously [56].The fluorescence increase was followed for 1 h with two reads per min with an excitation wavelength of 320 ± 8 nm and an emission wavelength of 430 ± 8 nm. For all measurements, positive (enzyme, substrate, DMSO, and buffer) and negative (substrate, DMSO, and buffer) controls were included in every measurement and were set as 100 and 0%, respectively. The measured values were normalized accordingly. For HDAC1, 6 and 8 Nexturastat A was used as positive control for a HDAC6-selective inhibitor. Dose−response curves were generated starting at 20 μM compound to generate 8-dose plots. IC_50_ values were then generated from the resulting plots.

### IC_50_ determination in AKT1 and AKT2

The inhibitory activity of the hybrids in the target enzymes was assayed at Reaction Biology company according to their RCB HotSpot Kinase Assay protocol [67]. The substrates were prepared in Reaction Buffer (20 mM HEPES pH 7.5, 10 mM MgCl_2_, 1 mM EGTA, 0.01 % Brij35, 0.02 mg/mL BSA, 0.1 mM Na_3_VO_4_ 2 mM DTT 1 % DMSO) and any required cofactors were added individually to each kinase reaction solution. The kinase was then added to the solution and the contents were gently mixed. The tested compounds were dissolved in 100 % DMSO and added into the kinase reaction mixture by Acoustic technology (Echo550, nanoliter range), and incubated for 20 min at room temperature. After that, ^33^P-ATP was added into the reaction mixture to initiate the reaction (10 µM and Km concentrations), and the solution was incubated for 2 h at room temperature. The kinase activity was detected by P81-filter-binding method.

### Western Blotting

Total protein extraction was performed using a buffer containing 100 mM Tris (pH 7.6), 1% Triton X-100, 2 mM PMSF, 10 mM Na_3_VO_4_, 100 mM NaF, 10 mM Na_4_P_2_O_7_, and 4 mM EDTA. Equal amounts of protein (15 μg) from Jurkat cell line sample were subjected to SDS-PAGE in an electrophoresis device, followed by electrotransfer of the proteins to nitrocellulose membranes. The membranes were blocked with 5% non-fat dry milk and incubated with specific primary antibodies diluted in blocking buffer, followed by secondary antibodies conjugated to horseradish peroxidase (HRP). Western blot analysis was performed using a SuperSignalTM West Dura Extended Duration substrate system (Thermo Fisher Scientific) and a G:BOX Chemi XX6 gel document system (Syngene). Antibodies against P-AKT (SER473) (#4060), AKT (PAN) (#4685) and GAPD (#5174) were obtained from Cell Signaling Technology (Danvers, MA, USA).

### Molecular modelling

The proposed binding mode of compounds **6b** and **6k** within the AKT2 catalytic binding site was generated to rationalize their interactions and support SAR discussion. We generated AKT2’s kinase domain using crystal (PDB ID: 9C1W, UniProt: P31751, residues from K136 until end) as template, due to their high resolution and similarity between co-crystallized ligands and our series. Structures were prepared using Protein preparation wizard. To obtain the starting configuration for the systems without crystals, Glide docking was conducted (Glide v. 7.7) [75,76] For docking, we used default settings and defined residues with 13 Å around the co-crystallized ligand for the binding site. Docking was conducted using standard precision (SP) with restrictions to at least one interaction with the hinge region. Of note, docking and simulations using PDB ID 3E87 were also attempted, however, the ligand was not stable within the binding pocket.

### Molecular dynamics simulations

We simulated the AKT2 with our ligands using the Desmond MD simulation engine [77] and the OPLS4 force-field [78]. he prepared systems were solvated in a cubic box with the size of the box set as 10 Å minimum distance from the box edges to any atom of the protein with periodic bound conditions. TIP3P water model [79] as used to describe the solvent and the net charge was neutralized using Na^+^ ions. RESPA integrator timesteps of 2 fs for bonded and near and 6 fs for far were applied. The short-range coulombic interactions were treated using a cut-off value of 9.0 Å, whereas long-range coulombic interactions were estimated using the Smooth Particle Mesh Ewald method [80]. Before the production simulations, the systems were relaxed using the default Desmond relaxation protocol. Simulations were run in an NPT ensemble, with a temperature of 310 K (using the Nosé-Hoover thermostat [81,82]) and pressure of 1.01325 bar (Martyna-Tobias-Klein barostat [83]). For each LXR-ligand combination, five independent simulations of 500 ns were carried out, resulting in at least 2.5 µs simulation data for each system. Maestro simulation interaction analysis tool (Schrödinger, LLC) was used for the analysis of RMSD and interaction analysis. All molecular dynamics trajectories and raw data related to the protein-ligand interactions within the simulations are available in the repository: 10.5281/zenodo.13881230.

### MM/GBSA binding energy calculations

To gain deeper insight into the interactions between AKT2-ligands, we explored their predicted binding energies, employing the molecular mechanics-generalized Born surface area (MM/GBSA) approach as outlined [84]. MM/GBSA predicts the binding free energy of protein-ligand complexes [84,85]. The ligands’ ranking based on the free energy could be correlated to the experimental binding affinities, especially in a congeneric series. Every 20^th^ frame from the simulations was considered for the calculations. These were used as input files for the MM/GBSA calculations with thermal_mmgbsa.py script from Schrödinger package. Calculated free-binding energies (kcal/mol) are represented by the MM/GBSA and normalized by the number of heavy atoms (HAC), according to the following formula: Ligand Efficiency = Ln(binding energy)/(1 + Ln(HAC)).

## ASSOCIATED CONTENT

### Software and Data Availability Statement

The data used in this work are available in a compressed folder named “Supplementary data”. The KNIME workflows, containing the input file, and models are available in the repository at 10.5281/zenodo.14106193. All molecular dynamics trajectories and, respective scripts and raw files/data related to the protein-ligand interactions within the simulations are available in the repository: 10.5281/zenodo.13881230.

## Support Information

List of targets retrieved, dataset size for each target, activity thresholds, and *in vitro* assays results are available in the Support information file.

## Author Contributions

K.B.W., and V.M.A. conceptualized the project. K.B.W. designed the library of hybrids, collected and curated data used in the study. K.B.W. and H.J.B., and built and validated machine learning models used in the study. R.C.B. built the fragment-based contribution maps. K.B.W, V.A.M.S., and M.T.T. synthesized and characterized the compounds. J.A.E.G.C. conducted cytotoxicity and western blotting assays. K.B.W and S.H. conducted enzymatic testing in HDAC isoforms. T.K. executed the MD experiments and respective analyses. M.S., W.S, J.A.M.N., and L.V.C.L. supervised biological assays. M.F.Z.J.T. supervised the synthesis. K.B.W., H.J.B., C.C.M.F., E.N.M., R.P.F., and T.K. wrote and edited the manuscript. V.M.A., R.P.F., and E.N.M. supervised the project.

## CONFLICT OF INTEREST

E.N.M. is co-founder of Predictive, LLC., which develops novel alternative methodologies and software for toxicity prediction. R.C.B. is CTO of InsilicAll Inc. All the other authors declare no conflicts.

## ACKNOWLEDGEMENTS

The authors are grateful to the Faculty of Pharmacy of University of São Paulo, the University of North Carolina, Chapel Hill, the Institut für Pharmazie of Martin-Luther Universität Halle-Wittenberg, and the Institute of Pharmaceutical Sciences of Eberhard-Karls-Universität which allowed the development of this work. This study was financed, in part, by the São Paulo Research Foundation (FAPESP), Brasil. Process Numbers #2021/08260-8, #2202/07275-4, #2023/07455-5, and 2024/07723-2. Coordenação de Aperfeiçoamento de Pessoal de Nível Superior (CAPES), also provided finantial support. This work was supported in large part by a grant from the National Institutes of Health, NIH R01GM140154 (to E.N.M.). T.K. is funded by the TüCAD2 and DZIF. TüCAD2 is funded by the Federal Ministry of Education and Research (BMBF) and the Baden-Württemberg Ministry of Science as part of the Excellence Strategy of the German Federal and State Governments. T.K. is funded by the German Center for Infection Research (DZIF, TTU06.716). S.H. and W.S. acknowledge funding by the German Research Foundation DFG (project number 469954457). The authors would like to thank the CSC-Finland for the generous computational resources.

## ABBREVIATIONS

AD: applicability domain
AKT2: Serine/threonine-protein kinase AKT2
BSA: bovine serum albumin
BTK: Bruton’s tyrosine kinase
CADD: computer-aided drug discovery
CCR: correct classification rate
CDK: Cyclin-dependent kinase
CHEK1: Serine/threonine-protein kinase Chk1
CK1α: Casein kinase 1α
CSK: Tyrosine-protein kinase CSK
CYP450: cytochrome P450
DCM: dichloromethane
DMPK: drug metabolism and pharmacokinetics
DMSO: dimethyl sulfoxide
DT: decision tree
EGFR: Epidermal growth factor receptor erbB1
EPHA2: Ephrin type-A receptor 2
EWG: electron-withdrawing group
EDG: electron-donating group
FDA: Food and Drug Administration
FGFR: Fibroblast growth factor receptor
FN: false negatives
FP: false positives
HAC: Heavy atom count
HAT: histone acetyltransferases
HBG: hinge-binding group
HCK: Tyrosine-protein kinase HCK
HDAC: histone deacetylase
HDACi: histone deacetylase inhibitor
HPLC: high performance liquid chromatography
HRMS: high resolution mass spectrometry
IC_50_: half-maximal inhibitory concentration
IGF1R: Insulin-like growth factor I receptor
INSR: Insulin receptor
JAK: Janus kinase
KDR: Vascular endothelial growth factor receptor 2
KIT: Stem cell growth factor receptor
MAP2K1: Dual specificity mitogen-activated protein kinase kinase 1
MAPK8: c-Jun N-terminal kinase 1
MAPK1: MAP kinase ERK2
MAPK12: MAP kinase p38 γ
MAPKAPK2: MAP kinase-activated protein kinase 2
MET: Hepatocyte growth factor receptor
NMR: nuclear magnetic resonance
ML: machine learning
MTT: 3-(4,5-dimethylthiazol-2-yl)-2,5-diphenyltetrazolium bromide
NPV: negative predictive values
PDK2: Pyruvate dehydrogenase kinase isoform 2
PDPK1: 3-phosphoinositide-dependent protein kinase-1
PI3K: Phosphoinositide 3-kinase
PIM1: Serine/threonine-protein kinase PIM1
PLK1: Serine/threonine-protein kinase PLK1
PPV: positive predictive values
QSAR: Quantitative structure–activity relationship
RF: random forest
SAHA: suberoylanilide hydroxamic acid or vorinostat
SAR: structure-activity relationship
SI: selectivity index
SRC: Tyrosine-protein kinase SRC
SYK: Tyrosine-protein kinase SYK
TEK: Tyrosine-protein kinase TIE-2
TFA: trifluoroacetic acid
TGFβR1: TGF-β receptor type I
THF: tetrahydrofuran
TLC: thin-layer chromatography
TN: true negatives
TNR: true negative rate
TP: true positives
TPR: true positive rate
TRKα: Tropomyosin receptor kinase
WHO: World Health Organization
ZBG: zinc-binding group

## Notes

https://doi.org/10.5281/zenodo.14106193

https://doi.org/10.5281/zenodo.13881230

## References

1. Amado J. Dona Flor e seus dois maridos: história moral e de amor. 1st ed. Companhia de Bolso, São Paulo.

2. World Health Organization. Cancer - Fact Sheet [Internet]. (2022). Available from: https://www.who.int/news-room/fact-sheets/detail/cancer.

3. Paliouras S, Pearson A, Barkalow F. The most successful oncology drug portfolios of the past decade. Nat Rev Drug Discov. 20(11), 811–812 (2021).

4. Roser M, Ritchie H. Cancer [Internet]. Our World in Data (2015). Available from: https://ourworldindata.org/cancer.

5. Roskoski R. Properties of FDA-approved small molecule protein kinase inhibitors: A 2023 update. Pharmacol Res [Internet]. 187(November 2022), 106552 (2023). Available from: 10.1016/j.phrs.2022.106552.

6. Attwood MM, Fabbro D, Sokolov A V., Knapp S, Schiöth HB. Trends in kinase drug discovery: targets, indications and inhibitor design. Nat Rev Drug Discov. 20(11), 839–861 (2021).

7. West AC, Johnstone RW. New and emerging HDAC inhibitors for cancer treatment. J Clin Invest. 124(1), 30–39 (2014).

8. Hanahan D. Hallmarks of Cancer: New Dimensions. Cancer Discov. 12(1), 31– 46 (2022).

9. Anighoro A, Bajorath J, Rastelli G. PolypharmacologyL: Challenges and Opportunities in Drug Discovery. J Med Chem. 57(19), 7874–87 (2014).

10. Waitman K, Parise-Filho R. New kinase and HDAC hybrid inhibitors: recent advances and perspectives. Future Med Chem. 14(10), 745–766 (2022).

11. Oki Y, Kelly KR, Flinn I, et al. CUDC-907 in relapsed/refractory diffuse large B-cell lymphoma, including patients with MYC-alterations: Results from an expanded phase I trial. Haematologica. 102(11), 1923–1930 (2017).

12. Lai CJ, Bao R, Tao X, et al. CUDC-101, a multitargeted inhibitor of histone deacetylase, epidermal growth factor receptor, and human epidermal growth factor receptor 2, exerts potent anticancer activity. Cancer Res. 70(9), 3647– 3656 (2010).

13. USA NL of M. NCT03874234 Clinical Trial [Internet]. ClinicalTrials.gov (2021). Available from: https://clinicaltrials.gov/ct2/show/NCT03874234.

14. America) P (Pharmaceutical R and M of. PHRMA Industry Profile 2021 [Internet]. 2021 Profile Biopharmaceutical Research Industry (2021). Available from: https://www.phrma.org/resource-center/Topics/Research-and-Development/Industry-Profile-2021.

15. Vamathevan J, Clark D, Czodrowski P, et al. Applications of machine learning in drug discovery and development. Nat Rev Drug Discov. 18(6), 463–477 (2019).

16. Sun D, Macedonia C, Chen Z, et al. Can Machine Learning Overcome the 95% Failure Rate and Reality that Only 30% of Approved Cancer Drugs Meaningfully Extend Patient Survival? J Med Chem [Internet]. (2024). Available from: https://pubs.acs.org/doi/10.1021/acs.jmedchem.4c01684.

17. Mendez D, Gaulton A, Bento AP, et al. ChEMBL: Towards direct deposition of bioassay data. Nucleic Acids Res. 47(D1), D930–D940 (2019).

18. Sessions Z, Sánchez-Cruz N, Prieto-Martínez FD, et al. Recent progress on cheminformatics approaches to epigenetic drug discovery. Drug Discov Today. 25(12), 2268–2276 (2020).

19. Alves VM, Korn D, Pervitsky V, et al. Knowledge-based approaches to drug discovery for rare diseases. Drug Discov Today. 27(2), 490–502 (2022).

20. Borba JVB, Braga RC, Alves VM, et al. Pred-Skin: A Web Portal for Accurate Prediction of Human Skin Sensitizers. Chem Res Toxicol. 34(2), 258–267 (2021).

21. Bennani FE, Doudach L, Karrouchi K, et al. Design and prediction of novel pyrazole derivatives as potential anti-cancer compounds based on 2D-2D-QSAR study against PC-3, B16F10, K562, MDA-MB-231, A2780, ACHN and NUGC cancer cell lines. Heliyon [Internet]. 8(8), e10003 (2022). Available from: 10.1016/j.heliyon.2022.e10003.

22. Umar AB, Uzairu A, Shallangwa GA, Uba S. QSAR modelling and molecular docking studies for anti-cancer compounds against melanoma cell line SK-MEL-2. Heliyon [Internet]. 6(3), e03640 (2020). Available from: https://linkinghub.elsevier.com/retrieve/pii/S2405844020304850.

23. Nagai J, Imamura M, Sakagami H, Uesawa Y. QSAR Prediction Model to Search for Compounds with Selective Cytotoxicity Against Oral Cell Cancer. Medicines. 6(2), 45 (2019).

24. Speck-Planche A. Multicellular Target QSAR Model for Simultaneous Prediction and Design of Anti-Pancreatic Cancer Agents. ACS Omega. 4(2), 3122–3132 (2019).

25. Ferreira GM, Magalhães JG de, Maltarollo VG, et al. QSAR studies on the human sirtuin 2 inhibition by non-covalent 7,5,2-anilinobenzamide derivatives. J Biomol Struct Dyn [Internet]. 38(2), 354–363 (2020). Available from: 10.1080/07391102.2019.1574603.

26. Melge AR, Parate S, Pavithran K, Koyakutty M, Mohan CG. Discovery of Anticancer Hybrid Molecules by Supervised Machine Learning Models and in Vitro Validation in Drug Resistant Chronic Myeloid Leukemia Cells. J Chem Inf Model. 62(4), 1126–1146 (2022).

27. Bohari MH, Srivastava HK, Sastry GN. Analogue-based approaches in anti-cancer compound modelling: the relevance of QSAR models. Org Med Chem Lett. 1(1), 1–12 (2011).

28. Arthur DE, Uzairu A, Mamza P, Abechi S. Quantitative structure–activity relationship study on potent anticancer compounds against MOLT-4 and P388 leukemia cell lines. J Adv Res [Internet]. 7(5), 823–837 (2016). Available from: 10.1016/j.jare.2016.03.010.

29. Waitman KB, de Almeida LC, Primi MC, et al. HDAC specificity and kinase off-targeting by purine-benzohydroxamate anti-hematological tumor agents. Eur J Med Chem [Internet]. 263, 115935 (2024). Available from: https://linkinghub.elsevier.com/retrieve/pii/S0223523423009029.

30. Kandoth C, McLellan MD, Vandin F, et al. Mutational landscape and significance across 12 major cancer types. Nature. 502(7471), 333–339 (2013).

31. Love C, Sun Z, Jima D, et al. The genetic landscape of mutations in Burkitt lymphoma. Nat Genet. 44(12), 1321–1325 (2012).

32. Kataoka K, Nagata Y, Kitanaka A, et al. Integrated molecular analysis of adult T cell leukemia/lymphoma. Nat Genet. 47(11), 1304–1315 (2015).

33. Losson H, Schnekenburger M, Dicato M, Diederich M. HDAC6—an emerging target against chronic myeloid leukemia? Cancers (Basel). 12(2) (2020).

34. Yuan S, Wang X, Hou S, et al. PHF6 and JAK3 mutations cooperate to drive T-cell acute lymphoblastic leukemia progression. Leukemia. 36(2), 370–382 (2022).

35. Wang X, Shen Y, Wang S, et al. PharmMapper 2017 update: A web server for potential drug target identification with a comprehensive target pharmacophore database. Nucleic Acids Res. 45(W1), W356–W360 (2017).

36. Daina A, Michielin O, Zoete V. SwissTargetPrediction: updated data and new features for efficient prediction of protein targets of small molecules. Nucleic Acids Res. 47(W1), W357–W3664 (2019).

37. Ji KY, Liu C, Liu ZQ, Deng YF, Hou TJ, Cao DS. Comprehensive assessment of nine target prediction web services: which should we choose for target fishing? Brief Bioinform. 24(2) (2023).

38. Fourches D, Muratov E, Tropsha A. Trust, but verify: On the importance of chemical structure curation in cheminformatics and QSAR modeling research. J Chem Inf Model. 50(7), 1189–1204 (2010).

39. ORGANISATION FOR ECONOMIC CO-OPERATION AND DEVELOPMENT (OECD). GUIDANCE DOCUMENT ON THE VALIDATION OF (QUANTITATIVE) STRUCTURE-ACTIVITY RELATIONSHIP [(Q)SAR] MODELS. Environment Health and Safety Publications, Series on Testing and Assessment, No. 69 [Internet]. France. Available from: https://read.oecd-ilibrary.org/environment/guidance-document-on-the-validation-of-quantitative-structure-activity-relationship-q-sar-models_9789264085442-en.

40. Tropsha A. Best practices for QSAR model development, validation, and exploitation. Mol Inform. 29(6–7), 476–488 (2010).

41. Alves VM, Auerbach SS, Kleinstreuer N, et al. Curated Data In - Trustworthy In Silico Models Out: The Impact of Data Quality on the Reliability of Artificial Intelligence Models as Alternatives to Animal Testing. Altern Lab Anim. 49(3), 73–82 (2021).

42. Landrum GA, Riniker S. Combining IC50 or Ki Values from Different Sources Is a Source of Significant Noise. J Chem Inf Model. 64(5), 1560–1567 (2024).

43. Zhang L, Zhang J, Jiang Q, Zhang L, Song W. Zinc binding groups for histone deacetylase inhibitors. J Enzyme Inhib Med Chem. 33(1), 714–721 (2018).

44. Roskoski R. Properties of FDA-approved small molecule protein kinase inhibitors. Pharmacol Res. 144(March), 19–50 (2019).

45. Raghavendra NM, Pingili D, Kadasi S, Mettu A, Prasad SVUM. Dual or multi-targeting inhibitors: The next generation anticancer agents. Eur J Med Chem [Internet]. 143, 1277–1300 (2018). Available from: 10.1016/j.ejmech.2017.10.021.

46. Manos-Turvey A, Watson EE, Sykes ML, et al. Synthesis and evaluation of phenoxymethylbenzamide analogues as anti-trypanosomal agents. Medchemcomm [Internet]. 6(3), 403–406 (2015). Available from: 10.1039/C4MD00406J.

47. Nepali K, Chang TY, Lai MJ, et al. Purine/purine isoster based scaffolds as new derivatives of benzamide class of HDAC inhibitors. Eur J Med Chem [Internet]. 196, 112291 (2020). Available from: 10.1016/j.ejmech.2020.112291.

48. Hull EE, Montgomery MR, Leyva KJ. HDAC Inhibitors as Epigenetic Regulators of the Immune System: Impacts on Cancer Therapy and Inflammatory Diseases. Biomed Res Int. 2016 (2016).

49. Lucet IS, Fantino E, Styles M, et al. The structural basis of Janus kinase 2 inhibition by a potent and specific pan-Janus kinase inhibitor. Blood. 107(1), 176–183 (2006).

50. Soumyanarayanan U, Ramanujulu PM, Mustafa N, et al. Discovery of a potent histone deacetylase (HDAC) 3/6 selective dual inhibitor. Eur J Med Chem. 184 (2019).

51. Porter NJ, Mahendran A, Breslow R, Christianson DW. Unusual zinc-binding mode of HDAC6-selective hydroxamate inhibitors. Proc Natl Acad Sci U S A. 114(51), 13459–13464 (2017).

52. Mak JYW, Wu KC, Gupta PK, et al. HDAC7 Inhibition by Phenacetyl and Phenylbenzoyl Hydroxamates. J Med Chem. 64(4), 2186–2204 (2021).

53. Melesina J, Simoben C V., Praetorius L, Bülbül EF, Robaa D, Sippl W. Strategies To Design Selective Histone Deacetylase Inhibitors. ChemMedChem. 16(9), 1336–1359 (2021).

54. Zhang J, Yang PL, Gray NS. Targeting cancer with small molecule kinase inhibitors. Nat Rev Cancer. 9(1), 28–39 (2009).

55. Walter NM, Wentsch HK, Bührmann M, et al. Design, Synthesis, and Biological Evaluation of Novel Type I1/2 p38α MAP Kinase Inhibitors with Excellent Selectivity, High Potency, and Prolonged Target Residence Time by Interfering with the R-Spine. J Med Chem. 60(19), 8027–8054 (2017).

56. Zessin M, Kutil Z, Meleshin M, et al. One-Atom substitution enables direct and continuous monitoring of histone deacylase activity. Biochemistry 58(48), 4777–4789 (2019).

57. Chan TO, Tsichlis PN. PDK2: A Complex Tail in One Akt. Sci Signal [Internet]. 2001(66) (2001). Available from: www.stke.org/cgi/content/full/OC_sigtrans;2001/66/pe1.

58. Yan S, Chen L, Zhuang H, et al. HDAC Inhibition Sensitize Hepatocellular Carcinoma to Lenvatinib via Suppressing AKT Activation. Int J Biol Sci. 20(8), 3046–3060 (2024).

59. An X, Wei Z, Ran B, et al. Histone Deacetylase Inhibitor Trichostatin A Suppresses Cell Proliferation and Induces Apoptosis by Regulating the PI3K/AKT Signalling Pathway in Gastric Cancer Cells. Anticancer Agents Med Chem. 20(17), 2114–2124 (2020).

60. Morgan SS, Cranmer LD. Vorinostat synergizes with ridaforolimus and abrogates the ridaforolimus-induced activation of AKT in synovial sarcoma cells. BMC Res Notes. 7(1) (2014).

61. Hao X, Xing W, Yuan J, et al. Cotargeting the JAK/STAT signaling pathway and histone deacetylase by ruxolitinib and vorinostat elicits synergistic effects against myeloproliferative neoplasms. Invest New Drugs. 38(3), 610–620 (2020).

62. Xia C, He Z, Cai Y, Liang S. Vorinostat upregulates MICA via the PI3K/Akt pathway to enhance the ability of natural killer cells to kill tumor cells. Eur J Pharmacol. 875 (2020).

63. Li X, Lu Y, Jin W, Liang K, Mills GB, Fan Z. Autophosphorylation of Akt at threonine 72 and serine 246: A potential mechanism of regulation of Akt kinase activity. Journal of Biological Chemistry. 281(19), 13837–13843 (2006).

64. Zhang P, Guo Z, Wu Y, et al. Histone deacetylase inhibitors inhibit the proliferation of gallbladder carcinoma cells by suppressing AKT/mTOR signaling. PLoS One. 10(8) (2015).

65. Porter NJ, Mahendran A, Breslow R, Christianson DW. Unusual zinc-binding mode of HDAC6-selective hydroxamate inhibitors. Proceedings of the National Academy of Sciences [Internet]. 114(51), 13459–13464 (2017). Available from: https://pnas.org/doi/full/10.1073/pnas.1718823114.

66. Nepali K, Lee HY, Liou JP. Nitro-Group-Containing Drugs. J Med Chem. 62(6), 2851–2893 (2019).

67. Melesina J, Simoben C V., Praetorius L, Bülbül EF, Robaa D, Sippl W. Strategies To Design Selective Histone Deacetylase Inhibitors. ChemMedChem. 16(9), 1336–1359 (2021).

68. Kumar CC, Madison V. AKT crystal structure and AKT-specific inhibitors. Oncogene. 24(50), 7493–7501 (2005).

69. Mendez D, Gaulton A, Bento AP, et al. ChEMBL: Towards direct deposition of bioassay data. Nucleic Acids Res. 47(D1), D930–D940 (2019).

70. Fourches D, Muratov E, Tropsha A. Trust, but Verify II: A Practical Guide to Chemogenomics Data Curation. J Chem Inf Model. 56(7), 1243–1252 (2016).

71. Riniker S, Landrum GA. Similarity maps - A visualization strategy for molecular fingerprints and machine-learning methods. J Cheminform. 5(9), 1–7 (2013).

72. Tropsha A, Golbraikh A. Predictive QSAR Modeling Workflow, Model Applicability Domains, and Virtual Screening. Curr Pharm Des. 13(34), 3494– 3504 (2007).

73. Johan van Meerloo, Gertjan J.L. Kaspers and JC. Cell Sensitivity Assays: The MTT Assay [Internet]. In: Methods Mol Biol, 79–91 (2011). Available from: http://www.springerlink.com/index/10.1007/978-1-61779-0805%5Cn, http://link.springer.com/10.1007/978-1-61779-080-5.

74. Heimburg T, Kolbinger FR, Zeyen P, et al. Structure-Based Design and Biological Characterization of Selective Histone Deacetylase 8 (HDAC8) Inhibitors with Anti-Neuroblastoma Activity. J Med Chem. 60(24), 10188–10204 (2017).

75. Friesner RA, Banks JL, Murphy RB, et al. Glide: A New Approach for Rapid, Accurate Docking and Scoring. 1. Method and Assessment of Docking Accuracy. J Med Chem. 47(7), 1750–1759 (2004).

76. Friesner RA, Murphy RB, Repasky MP, et al. Extra precision glide: Docking and scoring incorporating a model of hydrophobic enclosure for protein-ligand complexes. J Med Chem. 49(21), 6177–6196 (2006).

77. Bowers KJ, Chow E, Xu H, et al. Scalable Algorithms for Molecular Dynamics Simulations on Commodity Clusters. Proceedings of the 2006 ACM/IEEE Conference on Supercomputing, 1–13 (2006).

78. Lu C, Wu C, Ghoreishi D, et al. OPLS4: Improving force field accuracy on challenging regimes of chemical space. J Chem Theory Comput. 17(7), 4291– 4300 (2021).

79. Jorgensen WL, Chandrasekhar J, Madura JD, Impey RW, Klein ML. Comparison of simple potential functions for simulating liquid water. J Chem Phys. 79(2), 926–935 (1983).

80. Darden T, York D, Pedersen L. Particle mesh Ewald: An N·log(N) method for Ewald sums in large systems. J Chem Phys. 98(12), 10089–10092 (1993).

81. Hoover WG. Canonical dynamics: Equilibrium phase-space distributions. Phisical Review A. 31(3), 1695–1697 (1985).

82. Nosé S. A unified formulation of the constant temperature molecular dynamics methods. J Chem Phys. 81(1), 511–519 (1984).

83. Martyna GJ, Tuckerman ME, Tobias DJ, Klein ML. Explicit reversible integrators for extended systems dynamics. Mol Phys. 87(5), 1117–1157 (1996).

84. Li J, Abel R, Zhu K, Cao Y, Zhao S, Friesner RA. The VSGB 2.0 model: A next generation energy model for high resolution protein structure modeling. *Proteins: Structure*, Function and Bioinformatics. 79(10), 2794–2812 (2011).

85. Genheden S, Ryde U. The MM/PBSA and MM/GBSA methods to estimate ligand-binding affinities. Expert Opin Drug Discov 10(5), 449–461 (2015).

