## Supplementary Material for "Dona Flor and her two husbands: Discovery of novel HDAC6/AKT2 inhibitors for myeloid cancer treatment"

**TABLE OF CONTENTS**

**Supplementary Figures and Tables………………………………………………..……2**

**NMR Spectra………………………………………………………………………………....15**

**Chromatograms…………………………………………………………………………..…37**

**Supplementary Table S1:** List of hematological cancer cell lines selected for modeling and its associated malignancies.

| **Cell name** | **Type of malignancy** |
| --- | --- |
| HEL | Acute myeloid leukemia (erythroleukemia) |
| HL-60 | Acute promyelocytic leukemia |
| Jurkat | Acute T-cell leukemia/lymphoma |
| K562 | Chronic myeloid leucemia, BCR-ABL+ (erythroleukemia) |
| MM1S | Mature B-cell myeloma / Immunoglobulin A Labda myeloma |
| MOLT-4 | T-cell lymphobastic leukemia/lymphoma |
| MV4-11 | Acute myeloid leukemia / Biphenotypic B myelomonocytic leukemia |
| NALM-6 | B-cell lymphobastic leukemia/lymphoma / ALL |
| NB-4 | Acute myeloid leukemia / Acute promyelocytic leukemia |
| Raji | Burkitt lymphoma |
| THP1 | Acute monocytic leukemia / Acute myeloid leukemia |
| U266 | Mature B-cell neoplasm |
| U937 | Multiple myeloma |

**Supplementary Table S2:** Number of kinase targets retrieved for each hybrid compound in PharmMapper, SwissTargetPrediction, and the consensus between both methods.

| **Compound** | **Kinases PharmMapper** | **Kinases Swiss TargetPrediction** | **Consensus** |
| --- | --- | --- | --- |
| **5a** | 39 | 12 | 3 |
| **5b** | 37 | 22 | 5 |
| **5c** | 35 | 28 | 5 |
| **5d** | 33 | 31 | 7 |
| **5e** | 36 | 23 | 3 |
| **5f** | 32 | 24 | 2 |
| **5g** | 39 | 19 | 2 |
| **5h** | 39 | 12 | 1 |
| **5i** | 35 | 10 | 1 |
| **5j** | 35 | 25 | 5 |
| **5k** | 37 | 22 | 4 |
| **5l** | 36 | 22 | 3 |
| **5m** | 36 | 64 | 3 |
| **5n** | 34 | 42 | 3 |
| **5o** | 38 | 19 | 4 |
| **6a** | 39 | 8 | 0 |
| **6b** | 38 | 6 | 1 |
| **6c** | 39 | 7 | 2 |
| **6d** | 36 | 9 | 1 |
| **6e** | 36 | 20 | 3 |
| **6f** | 36 | 9 | 1 |
| **6g** | 40 | 12 | 1 |
| **6h** | 40 | 10 | 0 |
| **6i** | 36 | 13 | 3 |
| **6j** | 36 | 8 | 0 |
| **6k** | 38 | 9 | 1 |
| **6l** | 37 | 5 | 0 |
| **6m** | 36 | 21 | 5 |
| **6n** | 34 | 20 | 3 |
| **6o** | 37 | 13 | 1 |

**Supplementary Table S3:** Number of targets retrieved for each hybrid compound in PharmMapper, SwissTargetPrediction, and the consensus between both methods for all targets and for kinases. AH: hydroxamic acid; BZ: benzamide.

| **Compound** | **PharmMapper** | **Swiss TargetPrediction** | **Consensus** | **Kinases** |
| --- | --- | --- | --- | --- |
| **5a** | 297 | 100 | 9 | 1 |
| **5b** | 296 | 100 | 11 | 5 |
| **5c** | 299 | 100 | 9 | 3 |
| **5d** | 295 | 100 | 10 | 4 |
| **5e** | 295 | 100 | 5 | 2 |
| **5f** | 299 | 100 | 10 | 4 |
| **5g** | 299 | 100 | 12 | 1 |
| **5h** | 299 | 100 | 8 | 1 |
| **5i** | 299 | 100 | 5 | 0 |
| **5j** | 299 | 100 | 8 | 2 |
| **5k** | 299 | 100 | 8 | 1 |
| **5l** | 299 | 100 | 8 | 3 |
| **5m** | 299 | 100 | 3 | 3 |
| **5n** | 299 | 100 | 11 | 4 |
| **5o** | 299 | 100 | 10 | 2 |
| **6a** | 297 | 100 | 6 | 0 |
| **6b** | 296 | 100 | 8 | 1 |
| **6c** | 299 | 100 | 6 | 2 |
| **6d** | 295 | 100 | 6 | 1 |
| **6e** | 294 | 100 | 11 | 2 |
| **6f** | 299 | 100 | 7 | 3 |
| **6g** | 299 | 100 | 8 | 0 |
| **6h** | 299 | 100 | 7 | 1 |
| **6i** | 299 | 100 | 7 | 1 |
| **6j** | 299 | 100 | 10 | 1 |
| **6k** | 299 | 100 | 3 | 0 |
| **6l** | 299 | 100 | 11 | 2 |
| **6m** | 299 | 100 | 13 | 4 |
| **6n** | 299 | 100 | 9 | 6 |
| **6o** | 299 | 100 | 9 | 1 |

**Supplementary Table S4:** Initial number of compounds for each dataset, final dataset size, lost in curation rate (LIC), and activity thresholds employed in the construction of each cancer cell and enzyme model.

| **Dataset** | **Initial number  of compounds** | **Final dataset size** | **Lost in curation  (LIC) rate, %** | **Activity Threshold (nM)** |
| --- | --- | --- | --- | --- |
| HEL | 278 | 249 | 10.43 | 2.0 |
| HL-60 | 6062 | 5063 | 16.48 | 1 |
| Jurkat | 1135 | 947 | 16.56 | 0.5 |
| K562 | 5309 | 4583 | 13.67 | 1.0 |
| MM1S | 116 | 92 | 20.69 | 0.5 |
| MOLT-4 | 354 | 328 | 7.34 | 0.5 |
| MV4-11 | 431 | 417 | 3.25 | 0.05 |
| NALM-6 | 133 | 125 | 6.02 | 2.0 |
| NB-4 | 109 | 102 | 6.42 | 2.0 |
| Raji | 579 | 488 | 15.72 | 1.0 |
| THP1 | 488 | 456 | 6.56 | 2.0 |
| U266 | 106 | 101 | 4.72 | 1.0 |
| U937 | 859 | 769 | 10.48 | 1.0 |
| AKT2 | 992 | 621 | 37.40 | 10 |
| BTK | 2044 | 1606 | 21.43 | 10 |
| CDK4 | 1093 | 905 | 17.20 | 50 |
| CDK6 | 362 | 314 | 13.26 | 10 |
| CHEK1 | 1776 | 1644 | 7.43 | 10 |
| CK1α | 81 | 76 | 6.17 | 2000 |
| CSK | 119 | 102 | 14.29 | 500 |
| EGFR | 13786 | 8173 | 40.72 | 50 |
| EPHA2 | 167 | 132 | 20.96 | 1000 |
| FGFR1 | 4339 | 2647 | 39.00 | 10 |
| FGFR2 | 904 | 822 | 9.07 | 3 |
| HCK | 463 | 407 | 12.10 | 10 |
| HDAC1 | 4194 | 3404 | 18.84 | 100 |
| HDAC2 | 2136 | 1638 | 23.31 | 100 |
| HDAC3 | 1556 | 1294 | 16.84 | 100 |
| HDAC4 | 559 | 433 | 22.54 | 1000 |
| HDAC5 | 289 | 227 | 21.45 | 1000 |
| HDAC6 | 2886 | 2119 | 26.58 | 10 |
| HDAC7 | 317 | 259 | 18.30 | 1000 |
| HDAC8 | 1640 | 1343 | 18.11 | 500 |
| HDAC9 | 285 | 233 | 18.25 | 1000 |
| HDAC10 | 394 | 302 | 23.35 | 100 |
| IGFR1 | 4348 | 2797 | 35.67 | 10 |
| INSR | 944 | 875 | 7.31 | 100 |
| JAK1 | 3762 | 2970 | 21.05 | 10 |
| JAK2 | 5492 | 4514 | 17.81 | 10 |
| JAK3 | 3349 | 2951 | 11.88 | 10 |
| KIT | 1413 | 1128 | 20.17 | 10 |
| MAP2K1 | 943 | 790 | 16.22 | 10 |
| MAPK1 | 3484 | 2749 | 21.10 | 1 |
| MAPK12 | 117 | 104 | 11.11 | 100 |
| MAPK8 | 1730 | 1486 | 14.10 | 50 |
| MAPKAPK2 | 1031 | 985 | 4.46 | 100 |
| MET | 3522 | 2981 | 15.36 | 10 |
| PDK2 | 770 | 717 | 6.88 | 50 |
| PDPK1 | 1061 | 821 | 22.62 | 10 |
| PI3Kα | 4977 | 4496 | 9.66 | 10 |
| PI3Kβ | 1931 | 1673 | 13.36 | 100 |
| PI3Kδ | 2991 | 2520 | 15.75 | 10 |
| PI3Kγ | 2625 | 2303 | 12.27 | 50 |
| PIM1 | 3011 | 2477 | 17.73 | 5 |
| PLK1 | 770 | 685 | 11.04 | 1000 |
| SRC | 4839 | 3470 | 28.29 | 50 |
| SYK | 3562 | 3066 | 13.92 | 10 |
| TEK | 1170 | 949 | 18.89 | 50 |
| TGFβ1 | 1567 | 1004 | 35.93 | 10 |
| TRKα | 1735 | 1592 | 8.24 | 10 |
| VEGFR2/KDR | 8573 | 7364 | 14.10 | 50 |

**Supplementary Table S5:** Predicted activity outcomes for the hybrid compounds in the cancer cell models. Cells colored green: compound predicted active; cells colored gray: compound predicted inactive.

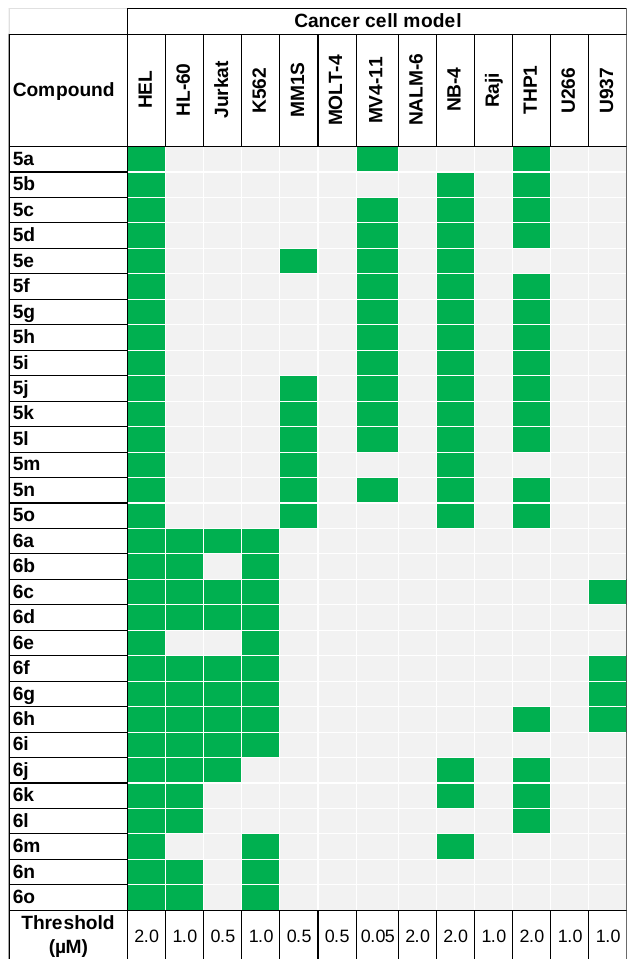

**Supplementary Table S6:** Predicted activity outcomes for the hybrid compounds in the kinase and HDAC models. Cells colored green: compound predicted active; cells colored gray: compound predicted inactive.

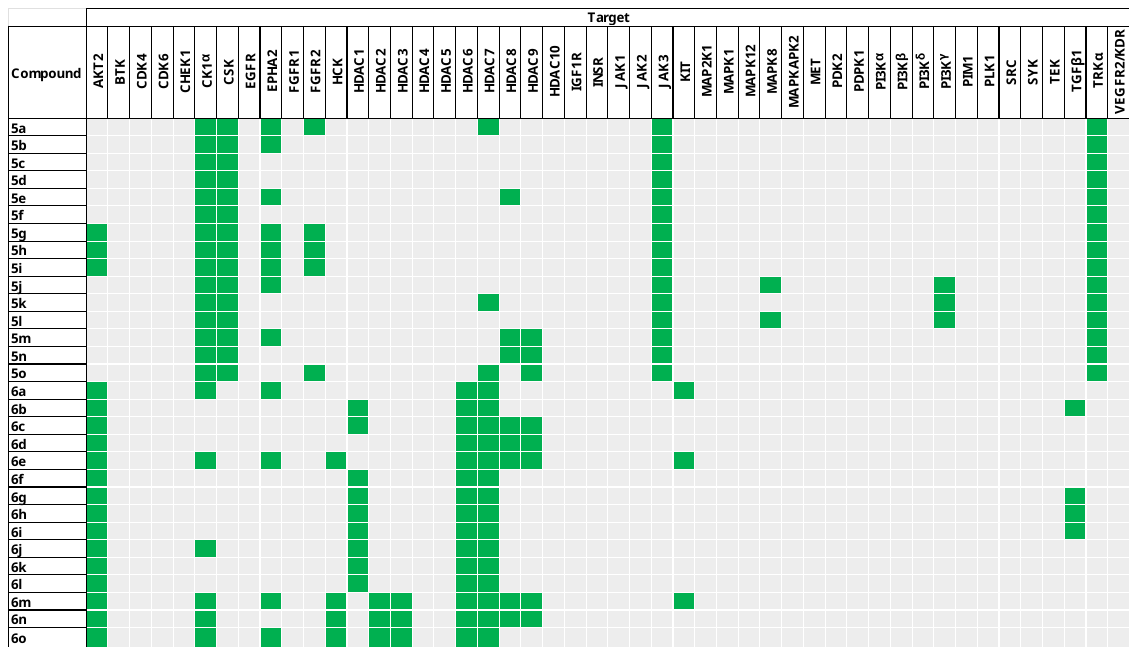

**Supplementary Table S7:** Predicted vulnerable cell lines and targets of the hybrids selected for synthesis and their IC_50_ threshold.

| **Cmp** | **Predicted vulnerable cancer cell lines and IC_50_ threshold (µM)** | **Predicted targets and**  **IC_50_ threshold (nM)** |
| --- | --- | --- |
| 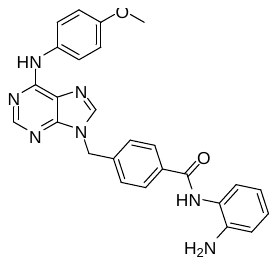  **5a** | HEL < 2  MV4-11 < 0.05  THP1 < 2 | CK1α < 2000 CSK < 500 EPHA2 < 1000 FGFR2 < 3 HDAC7 < 1000 JAK3 < 10 TRKα < 10 |
| 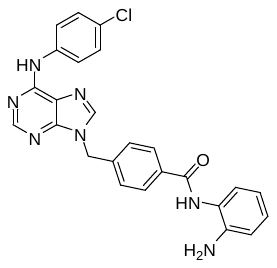  **5b** | HEL < 2  NB-4 < 2  THP1 < 2 | CK1α < 2000 CSK < 500 EPHA2 < 1000 JAK3 < 10 TRKα < 10 |
| 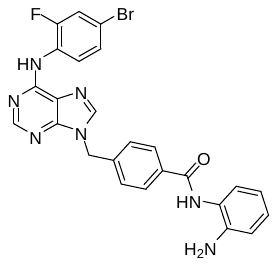  **5c** | HEL < 2  MV4-11 < 0.05  NB-4 < 2  THP1 < 2 | CK1α < 2000 CSK < 500 JAK3 < 10 TRKα < 10 |
| 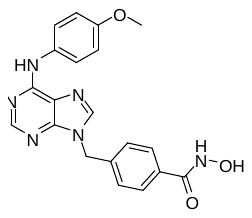  **6a** | HEL < 2  HL-60 < 1  Jurkat < 0.5  K562 < 1  MM1S < 0.5 | AKT2 < 10 CK1α < 2000 EPHA2 < 1000 HDAC6 < 10 HDAC7 < 1000 KIT < 10 |
| 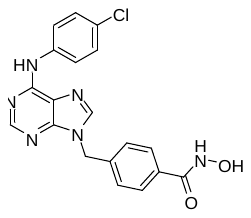  **6b** | HEL < 2  HL-60 < 1  K562 < 1  MM1S < 0.5 | AKT2 < 10 HDAC1 < 100 HDAC6 < 10 HDAC7 < 1000 TGFβ1 < 10 |
| 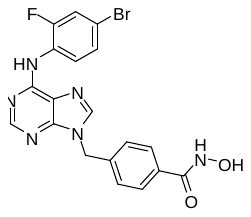  **6c** | HEL < 2  HL-60 < 1  Jurkat < 0.5  K562 < 1  MM1S < 0.5  U937 < 1 | AKT2 < 10 HDAC1 < 100 HDAC6 < 10 HDAC7 < 100 HDAC8 < 500 HDAC9 < 1000 |
| 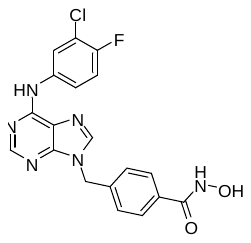  **6d** | HEL < 2  HL-60 < 1  Jurkat < 0.5  K562 < 1  MM1S < 0.5 | AKT2 < 10 HDAC6 < 10 HDAC7 < 1000 HDAC8 < 500 HDAC9 < 1000 |
| 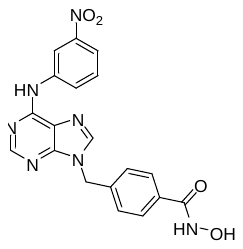  **6k** | HEL < 2  HL-60 < 1  NB-4 < 2  THP1 < 2 | AKT2 < 10 HDAC1 < 100 HDAC6 < 10 HDAC7 < 1000 |

**Figure S1**: HDAC1 dose-response curves of tested inhibitors. IC_50_ values are expressed in terms of the mean and standard deviation (mean ± SD), with *n* = 3.
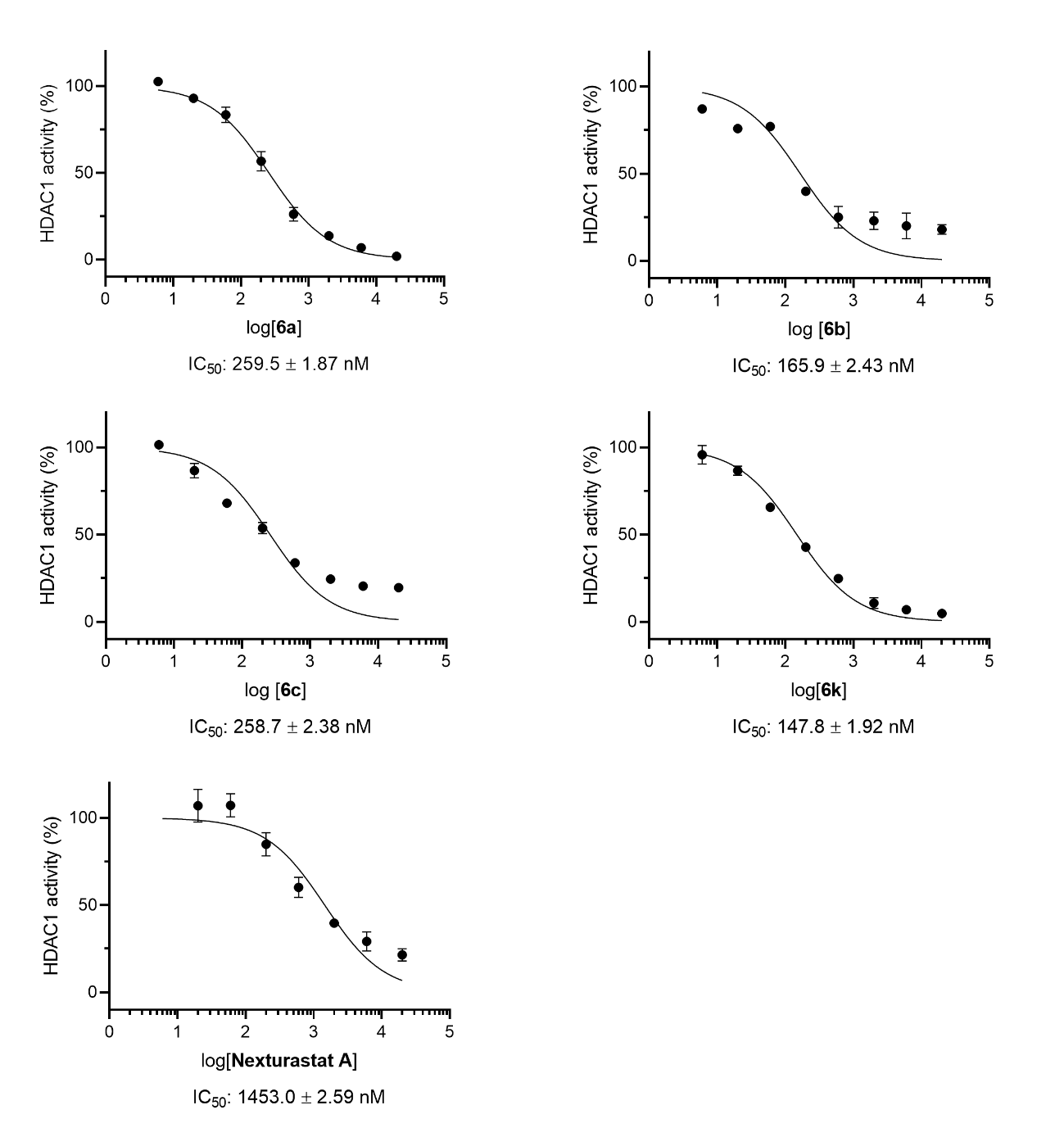

**Figure S2**: HDAC2, HDAC3, HDAC4, and HDAC5 dose-response curves of tested inhibitors. IC_50_ values are expressed in terms of the mean and standard deviation (mean ± SD), with *n* = 3.

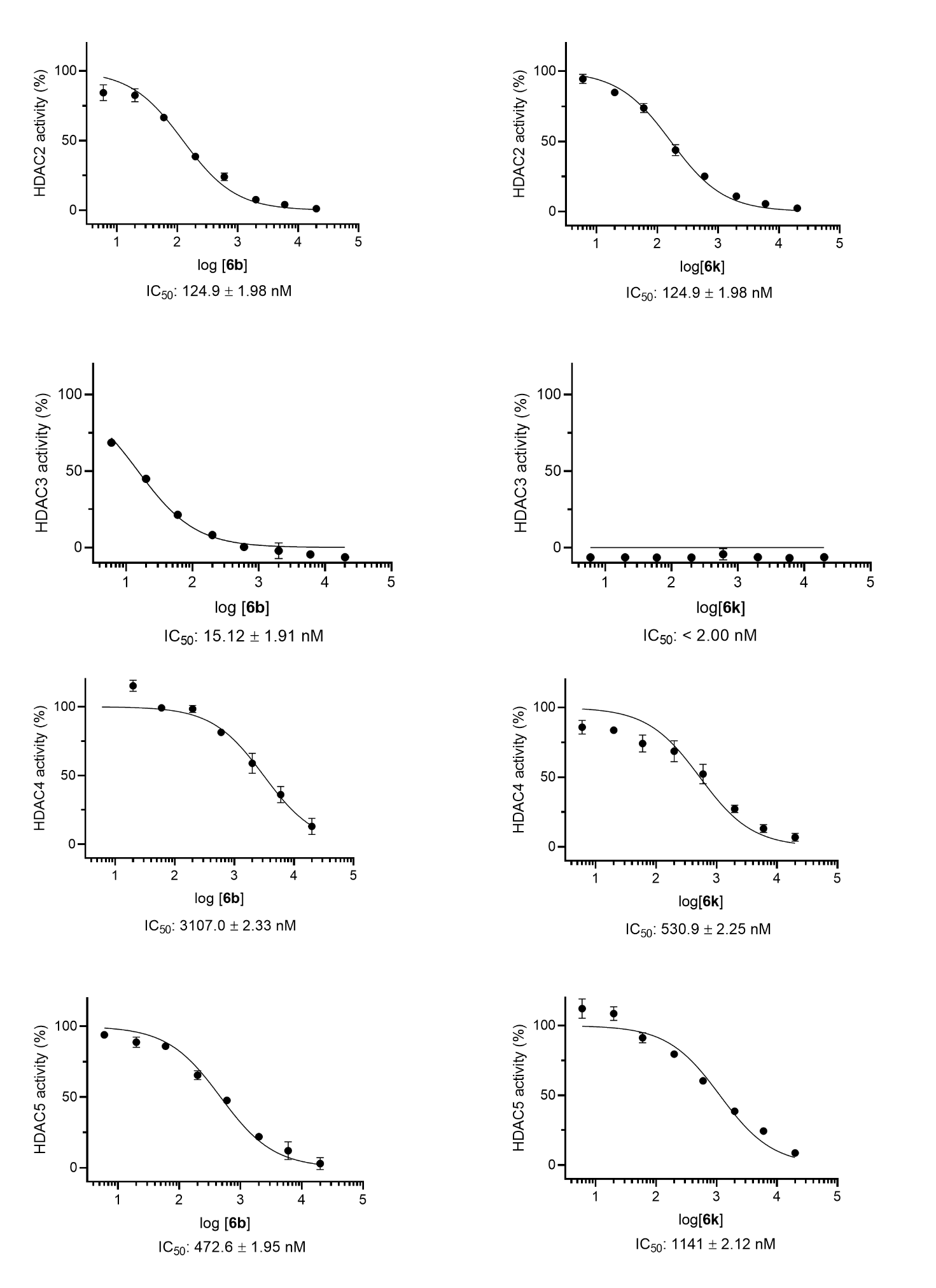

**Figure S3**: HDAC6 dose-response curves of tested inhibitors. IC_50_ values are expressed in terms of the mean and standard deviation (mean ± SD), with *n* = 3.

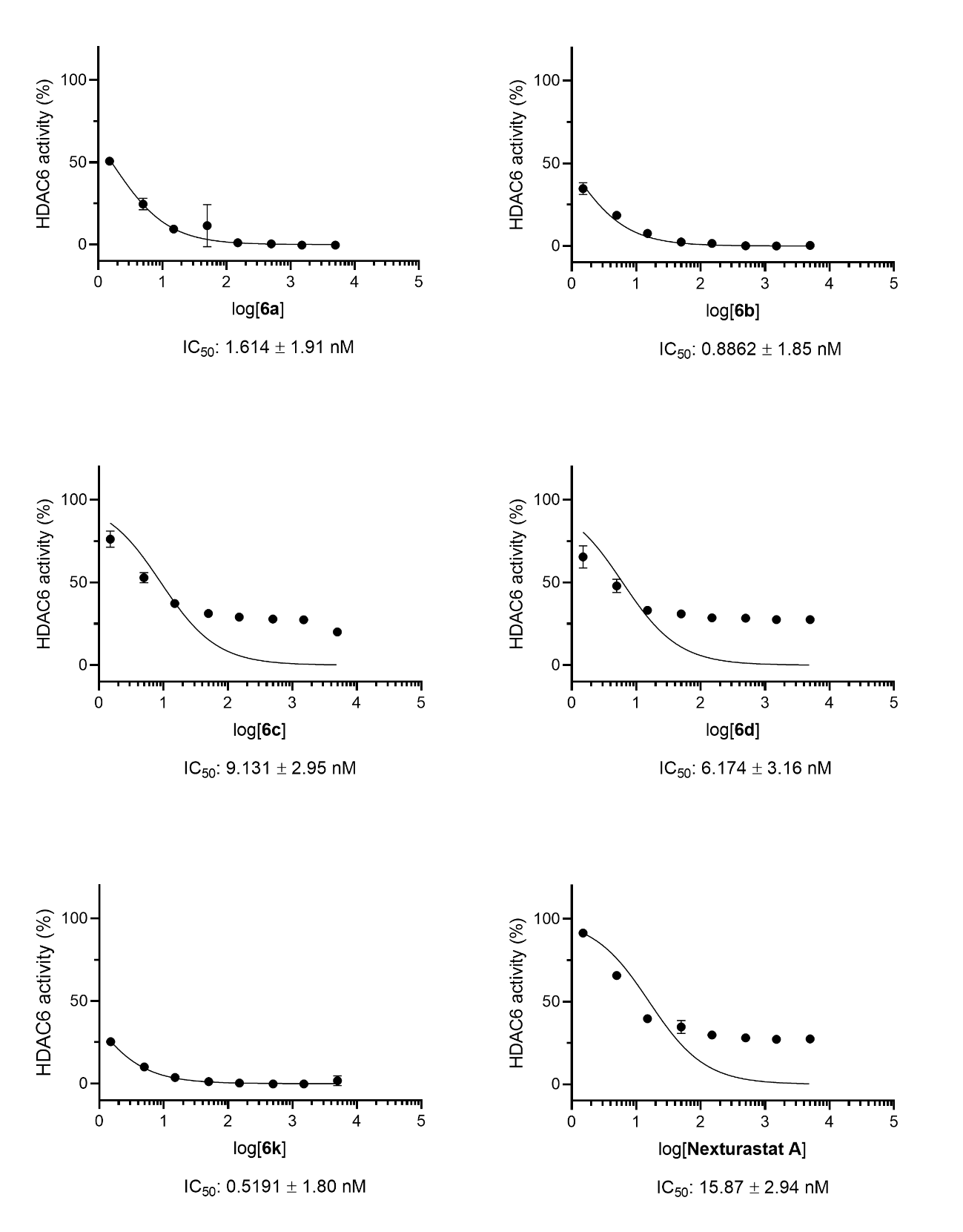

**Figure S4:** HDAC8 dose-response curves of tested inhibitors. IC_50_ values are expressed in terms of the mean and standard deviation (mean ± SD), with *n* = 3.

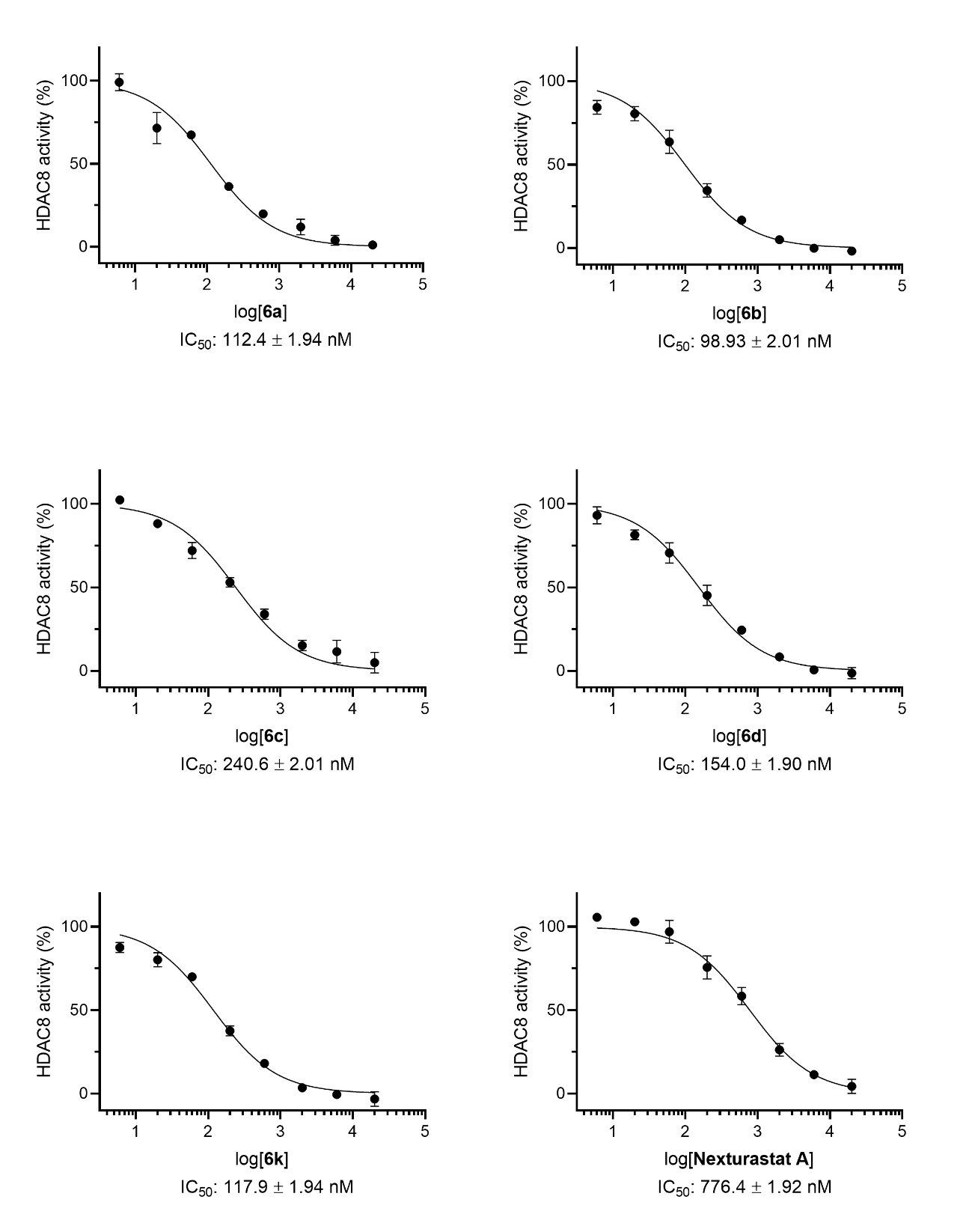

**Figure S5:** HDAC7 and HDAC9 dose-response curves of tested inhibitors. IC_50_ values are expressed in terms of the mean and standard deviation (mean ± SD), with *n* = 3.
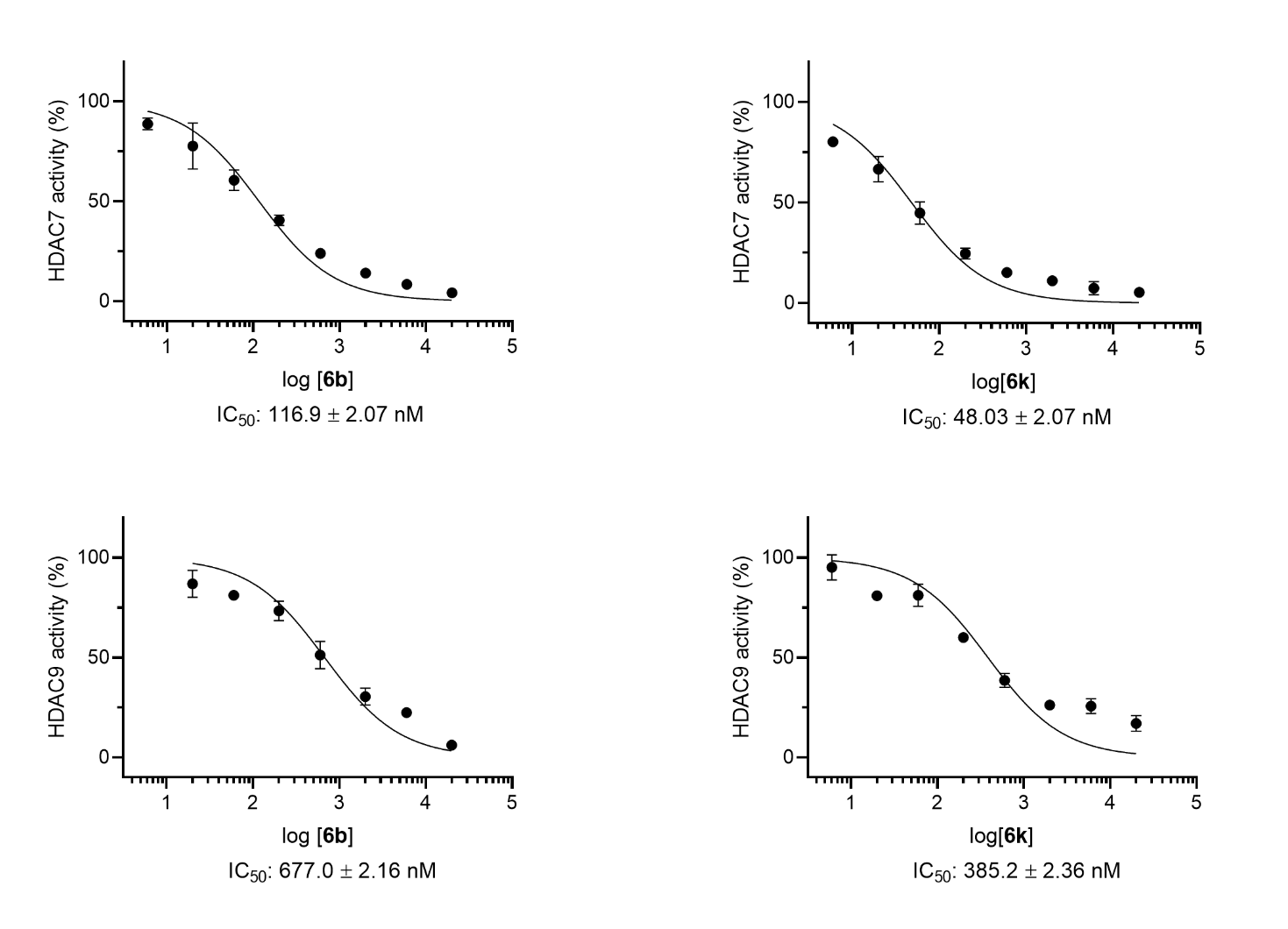

**Supplementary Figure S6:** ^1^H NMR spectra of the intermediate 2a (300 MHz, DMSO-d_6_, δ = ppm)

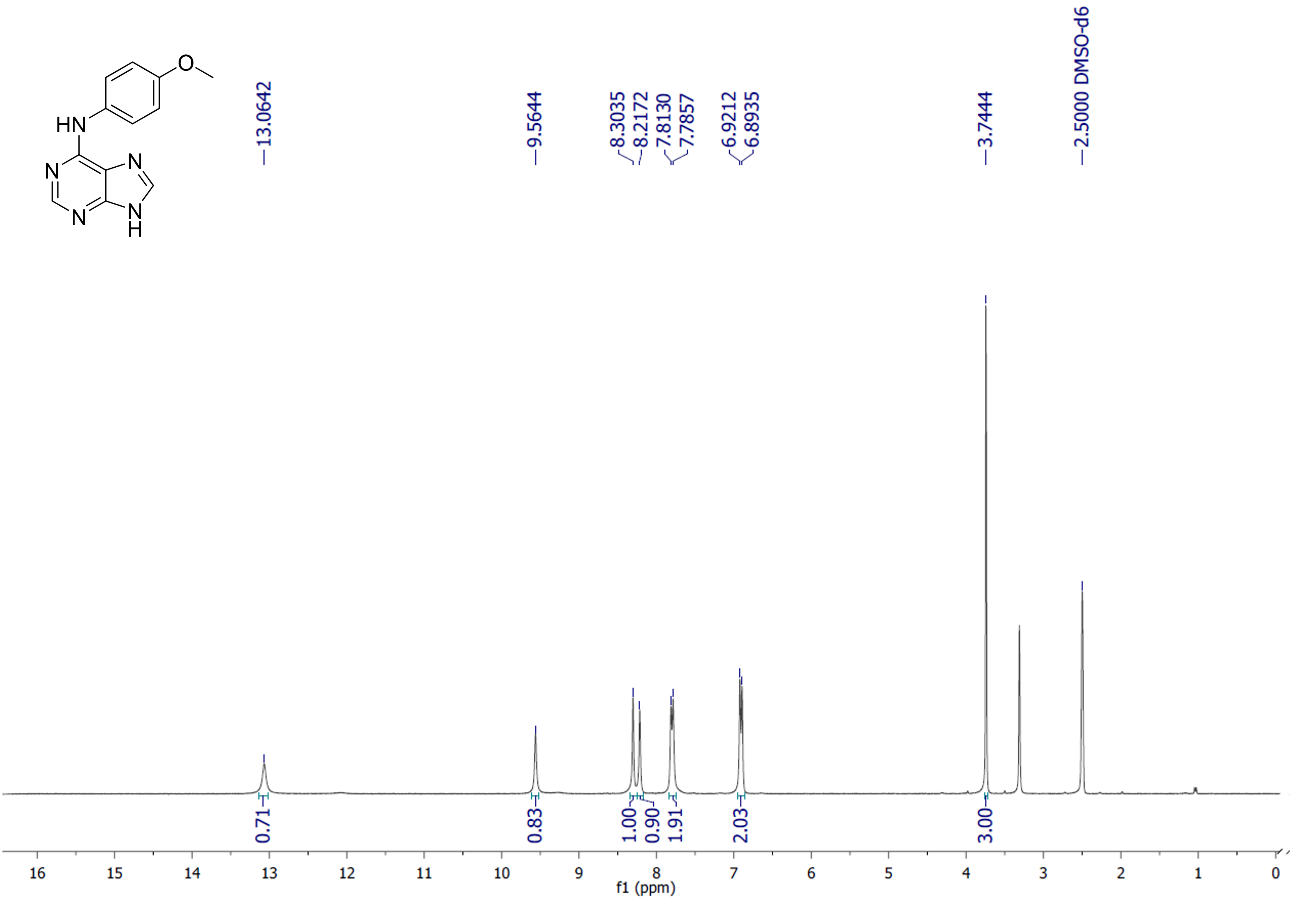

**Supplementary Figure S7:** ^13^C NMR spectra of the intermediate 2a (75 MHz, DMSO-d_6_, δ = ppm).

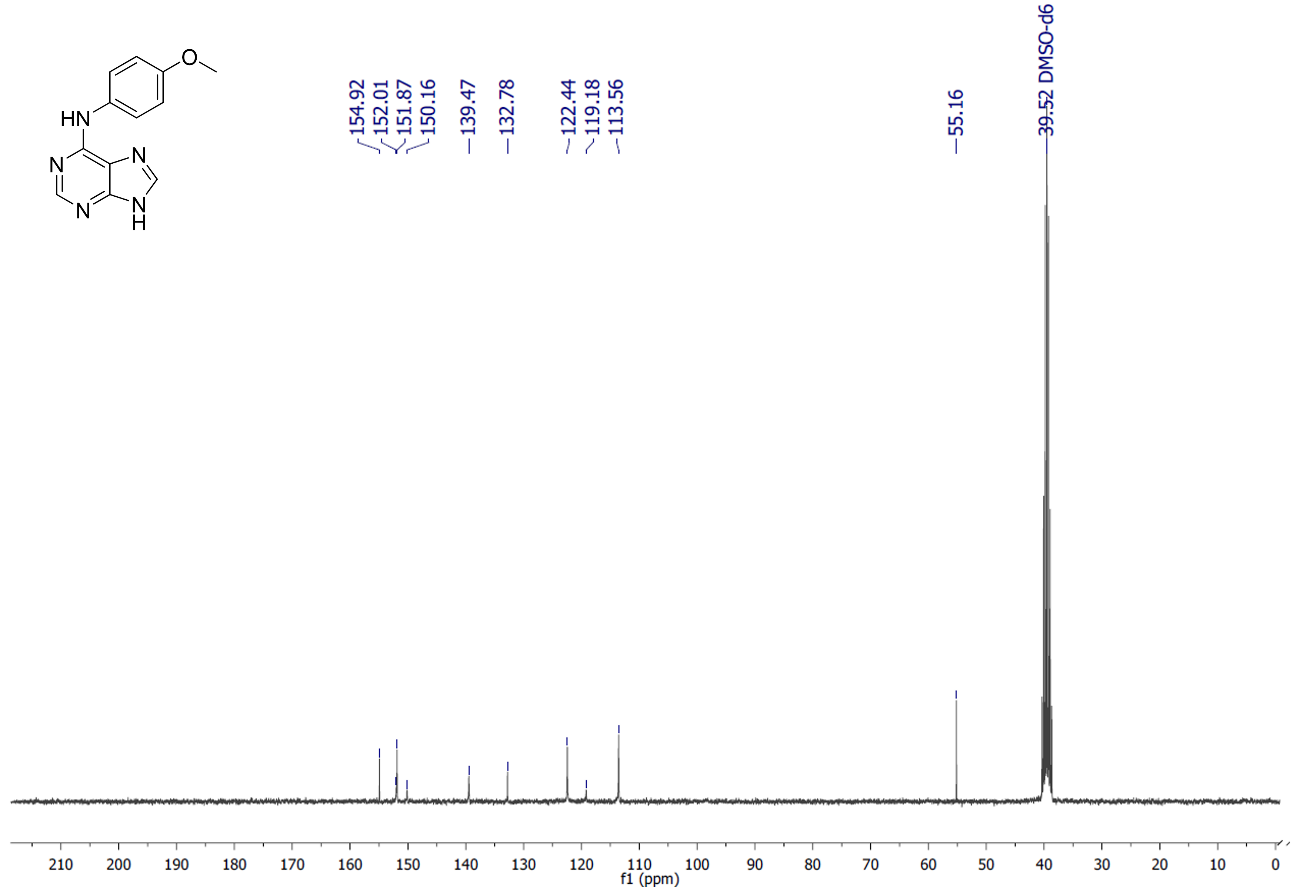

**Supplementary Figure S8:** ^1^H NMR spectra of the intermediate 2b (300 MHz, DMSO-d_6_, δ = ppm).

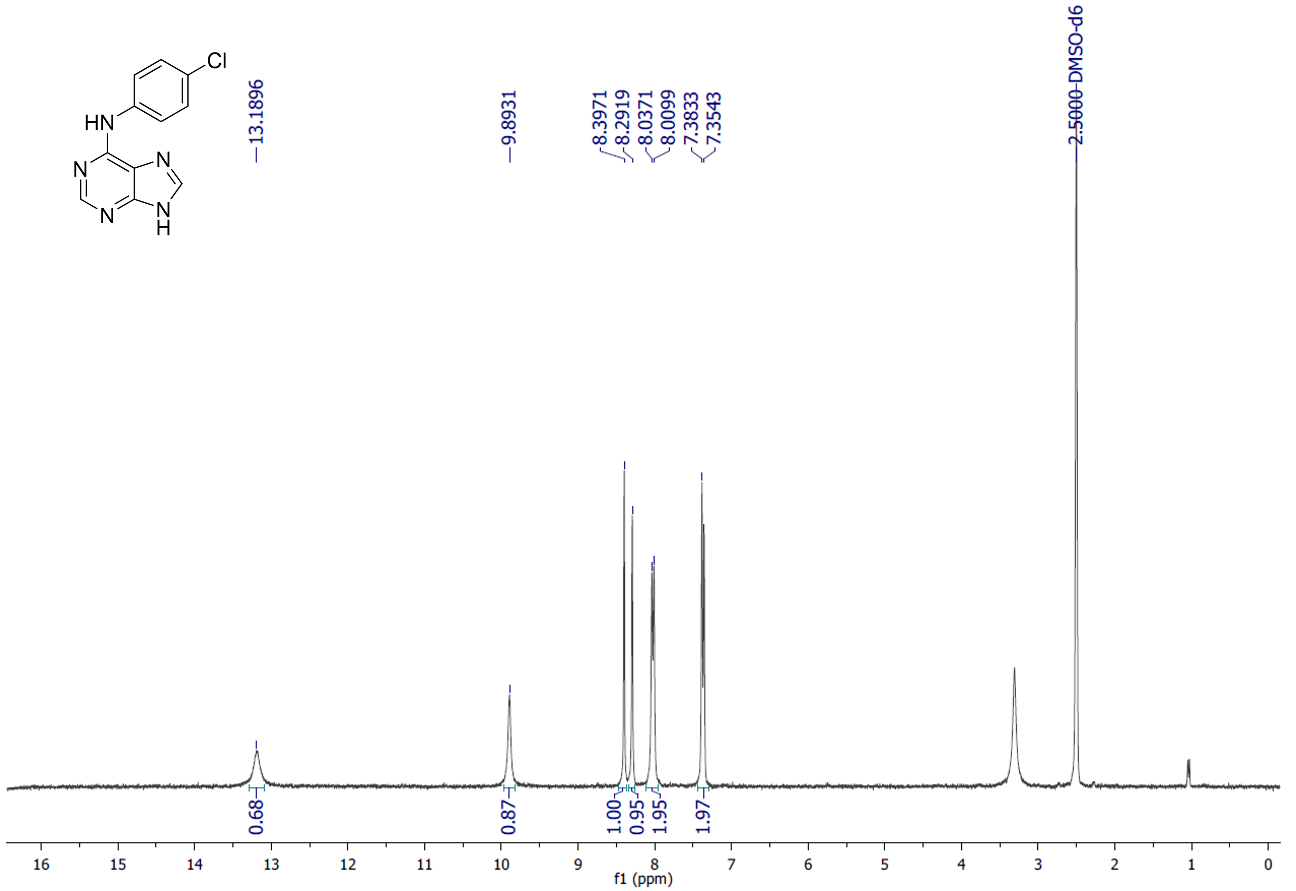

**Supplementary Figure S9:** ^13^C NMR spectra of the intermediate 2b (75 MHz, DMSO-d_6_, δ = ppm).

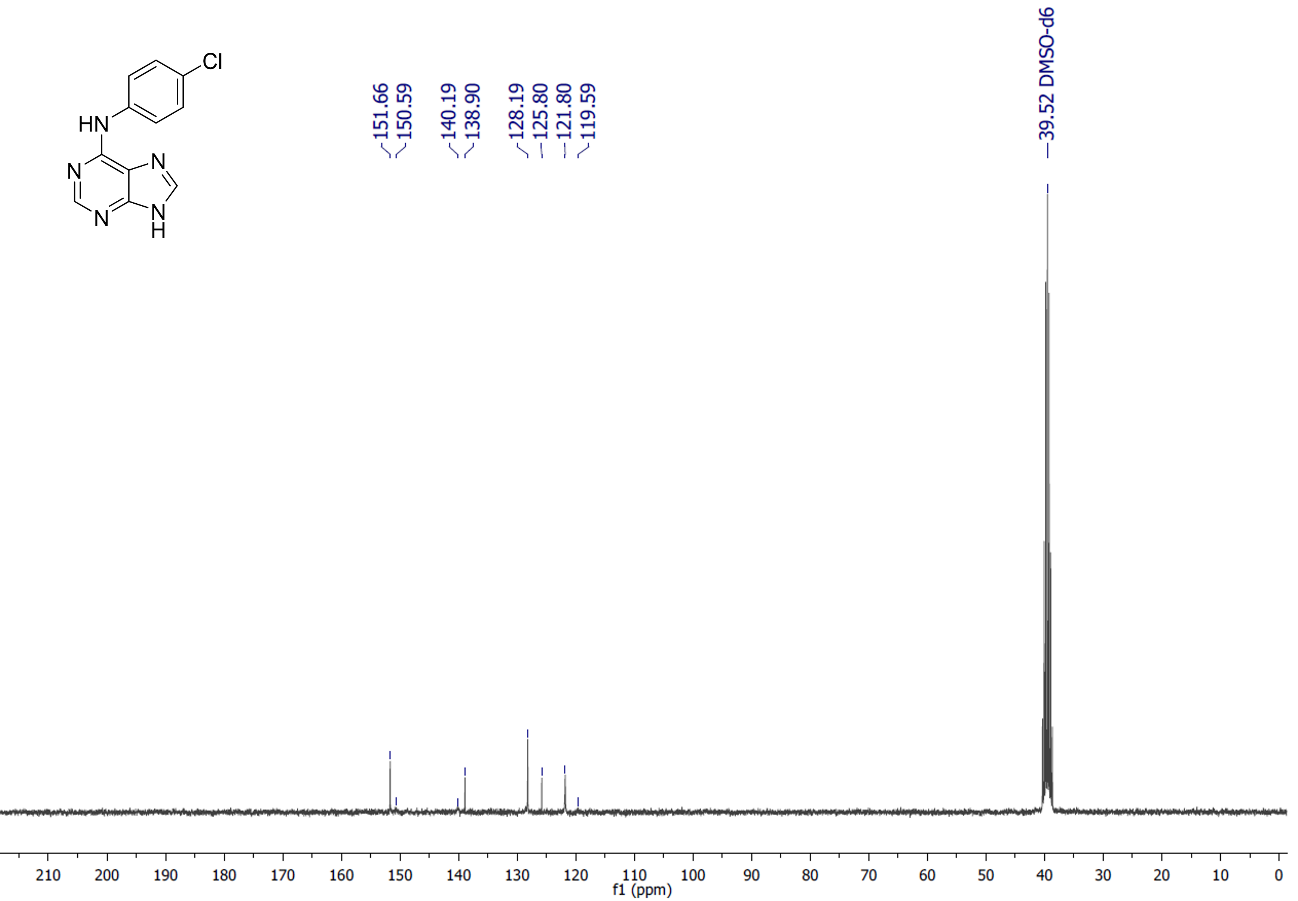

**Supplementary Figure S10:** ^1^H NMR spectra of the intermediate 2c (300 MHz, DMSO-d_6_, δ = ppm).

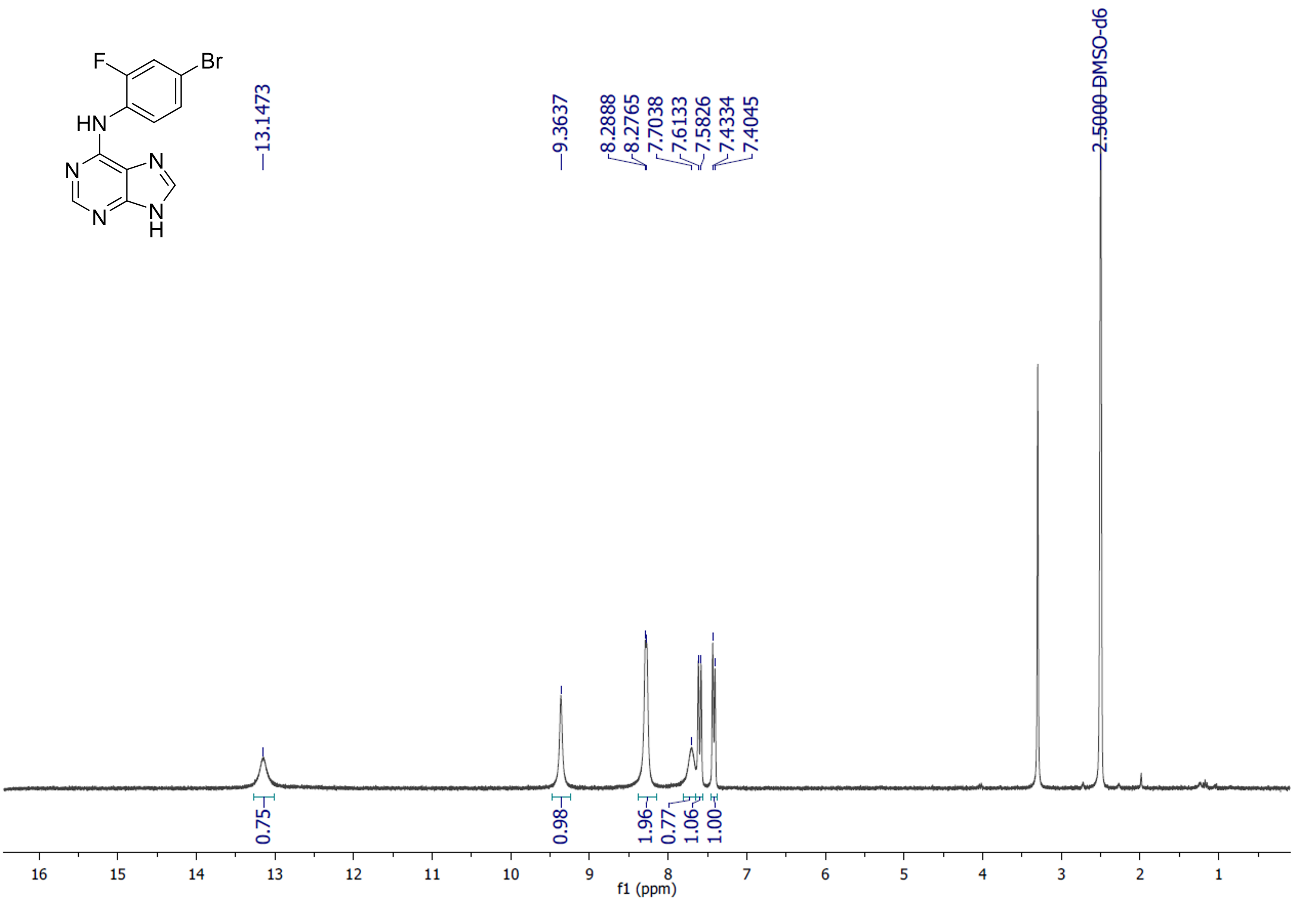
**Supplementary Figure S11*:*** ^13^C NMR spectra of the intermediate 2c (300 MHz, DMSO-d_6_, δ = ppm).
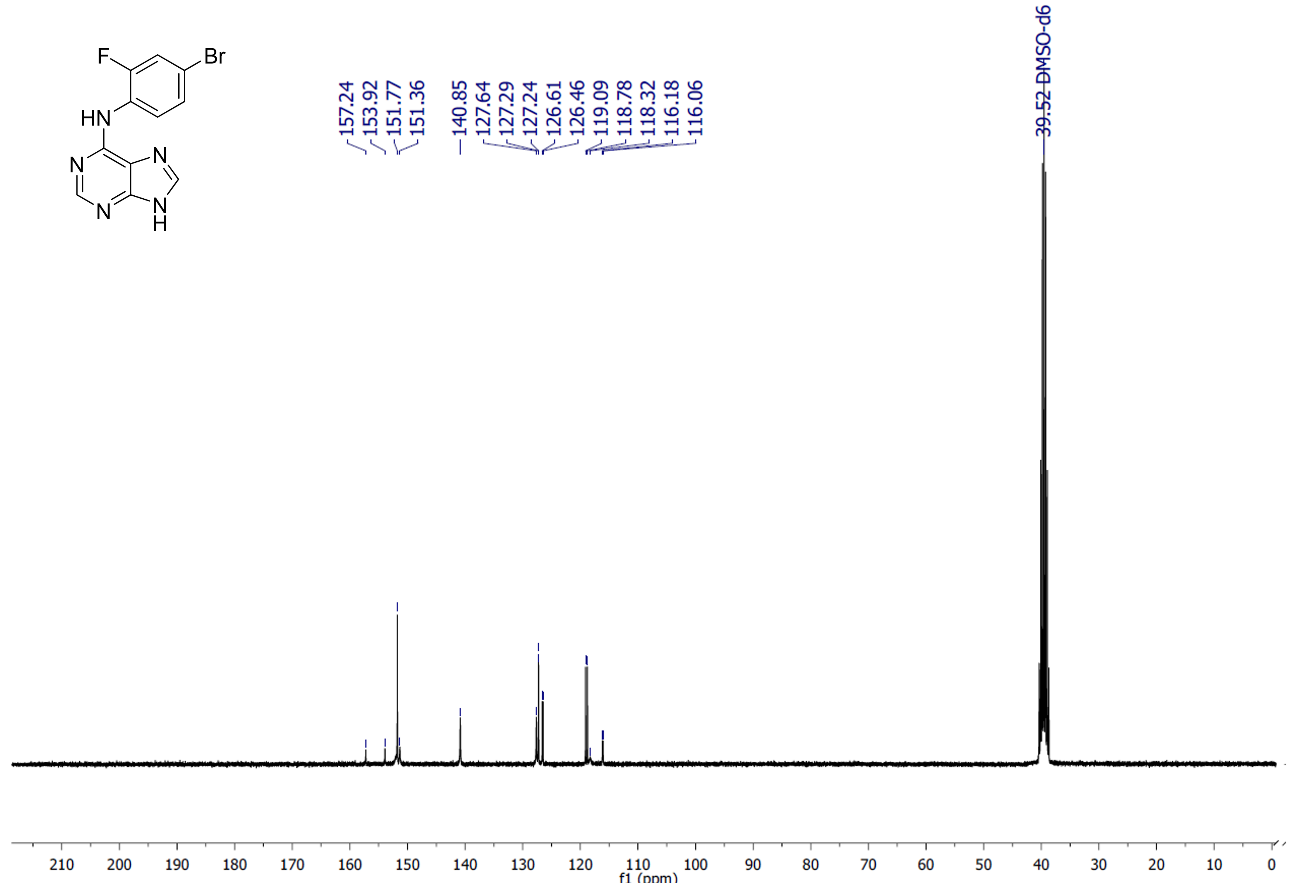

**Supplementary Figure S12:** ^1^H NMR spectra of the intermediate 2d (300 MHz, DMSO-d_6_, δ = ppm).

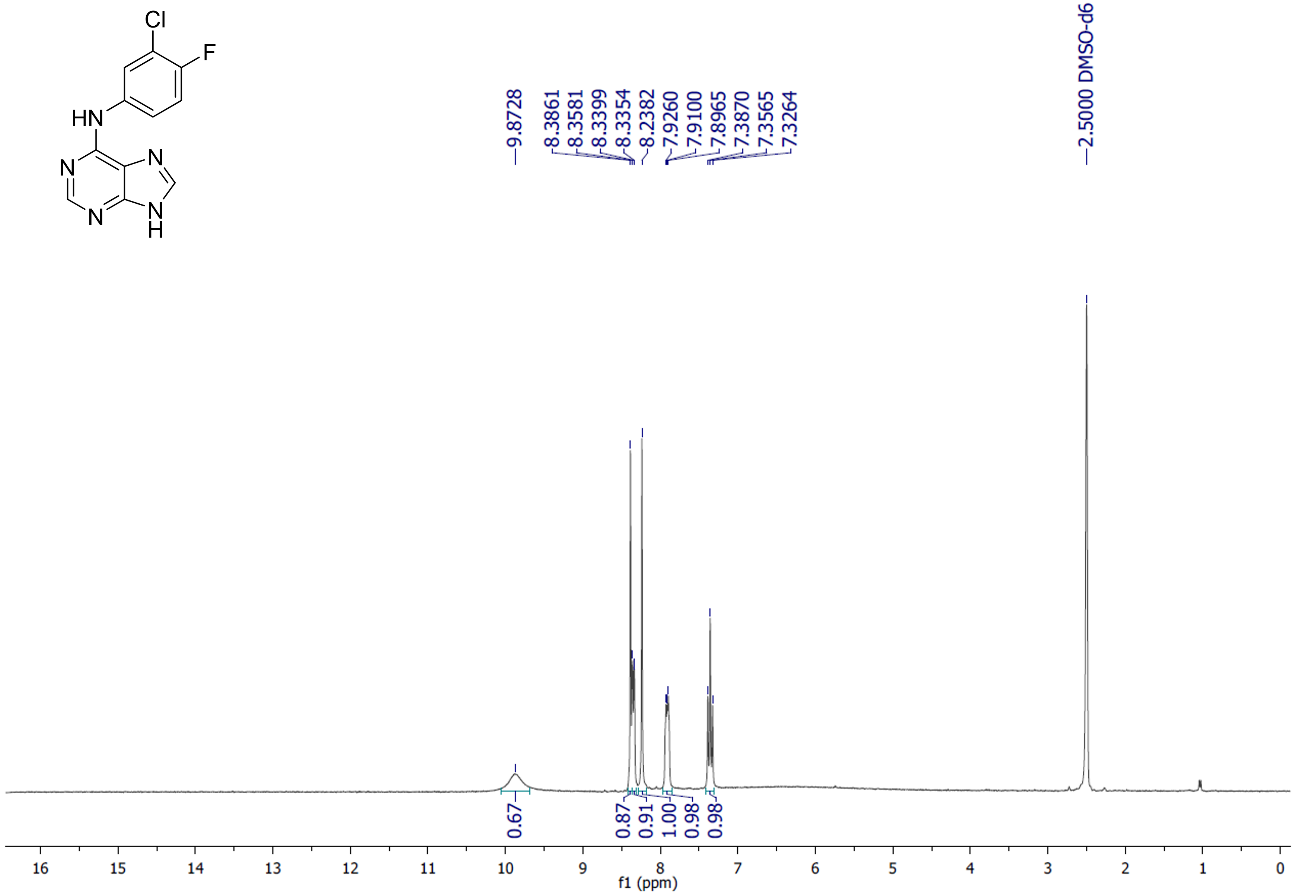

**Supplementary Figure S13:** ^13^C NMR spectra of the intermediate 2d (75 MHz, DMSO-d_6_, δ = ppm).

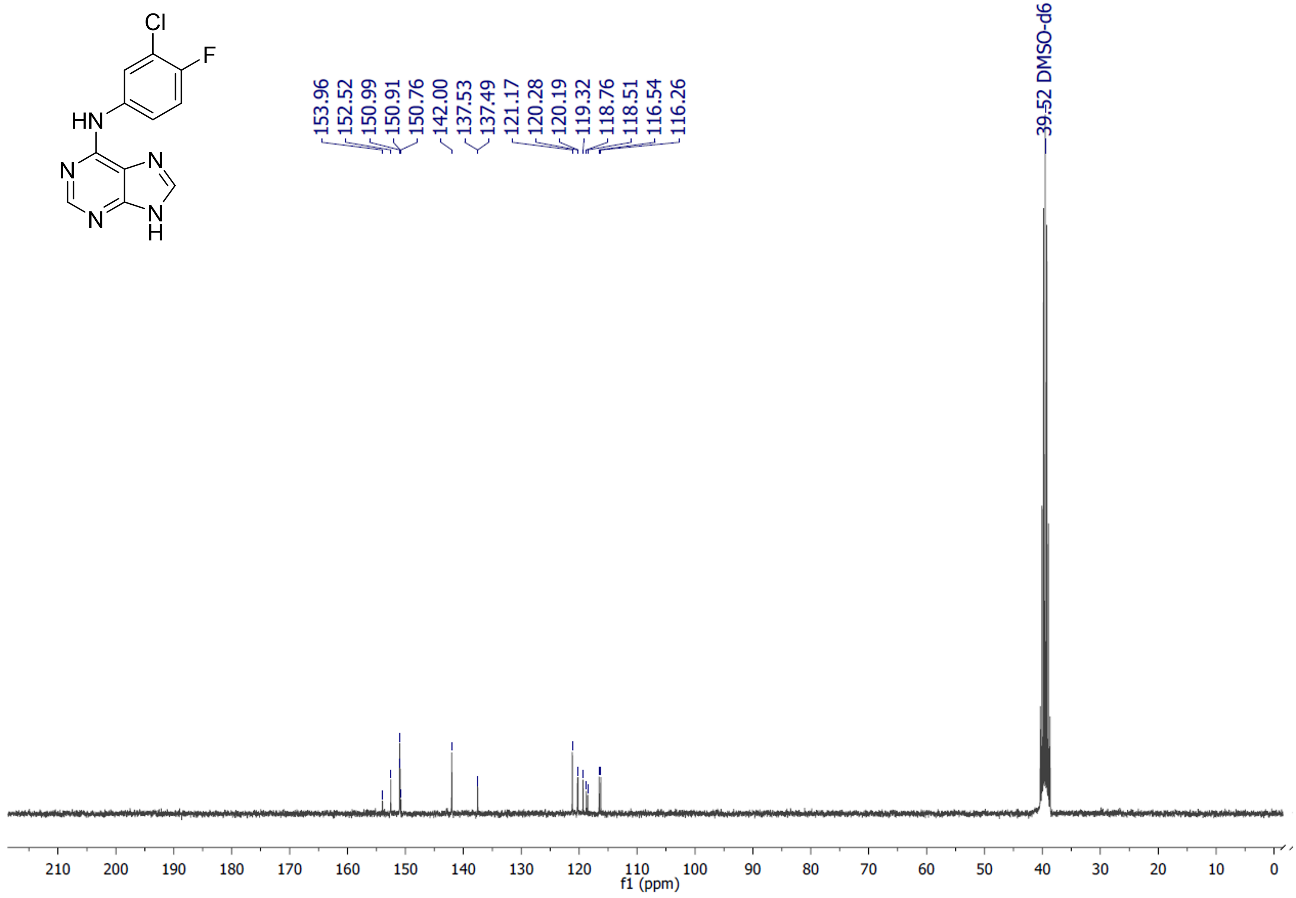

**Supplementary Figure S14:** ^1^H NMR spectra of the intermediate 2k (300 MHz, DMSO-d_6_, δ = ppm).

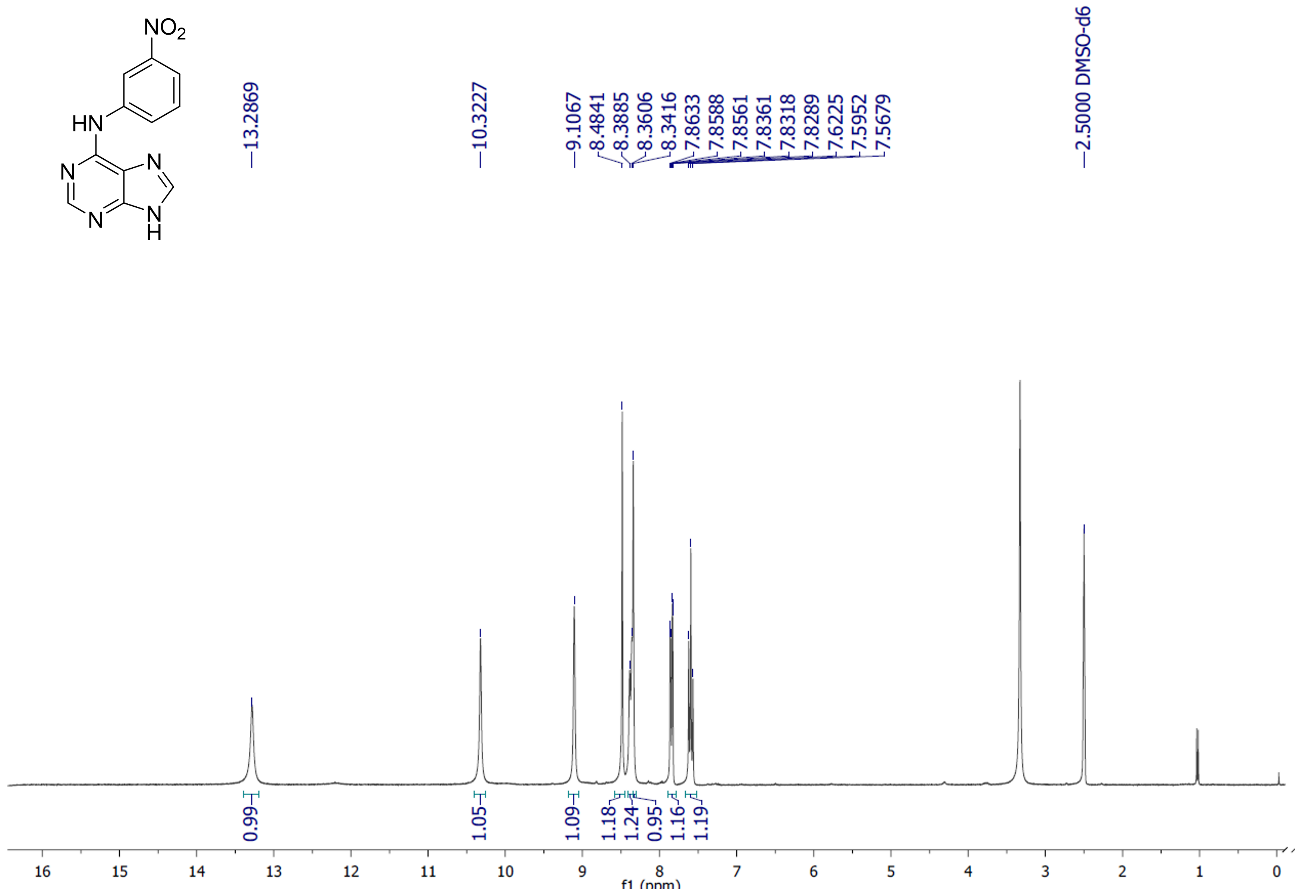

**Supplementary Figure S15:** ^13^C NMR spectra of the intermediate 2k (75 MHz, DMSO-d_6_, δ = ppm).

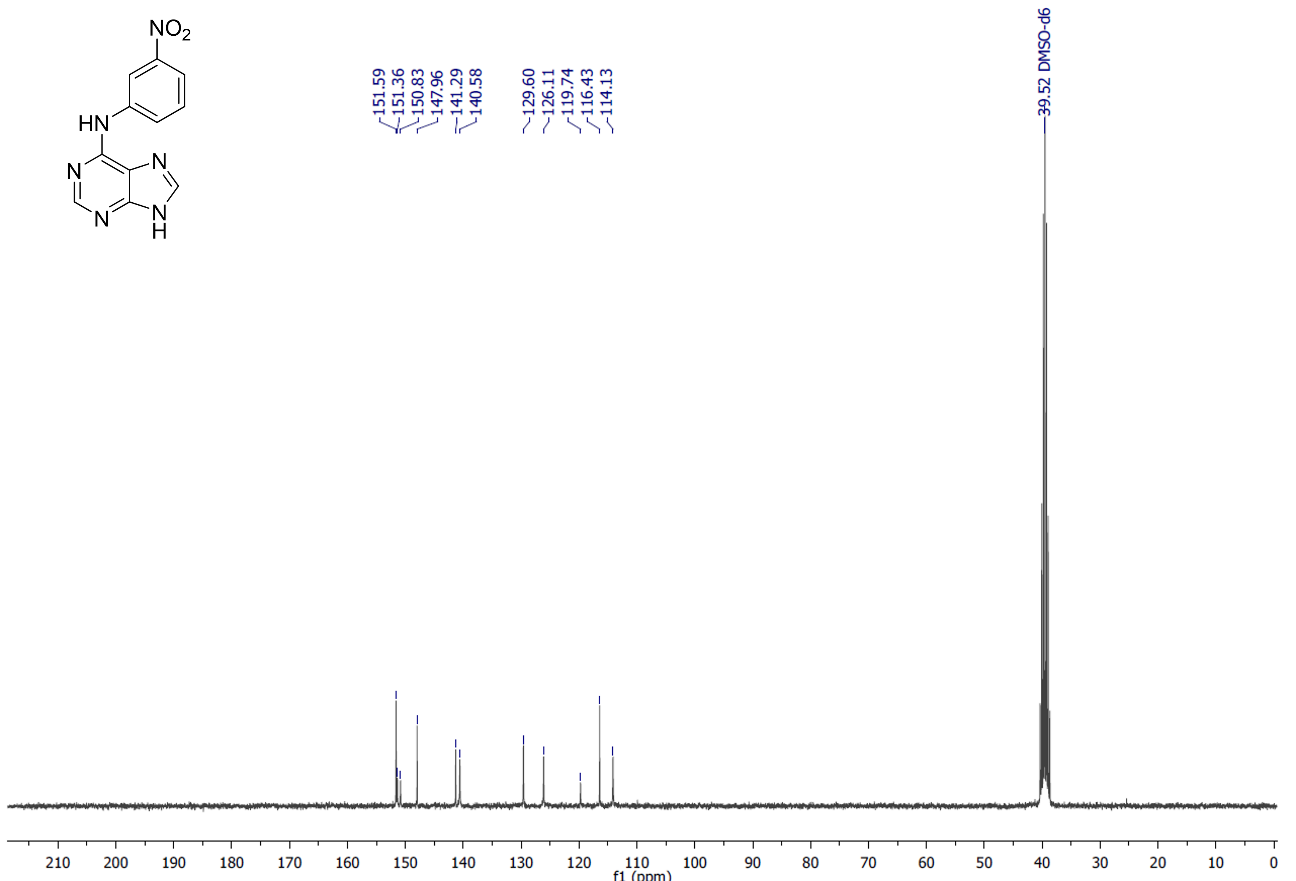

**Supplementary Figure S16:** ^1^H NMR spectra of the intermediate 3a (300 MHz, DMSO-d_6_, δ = ppm).

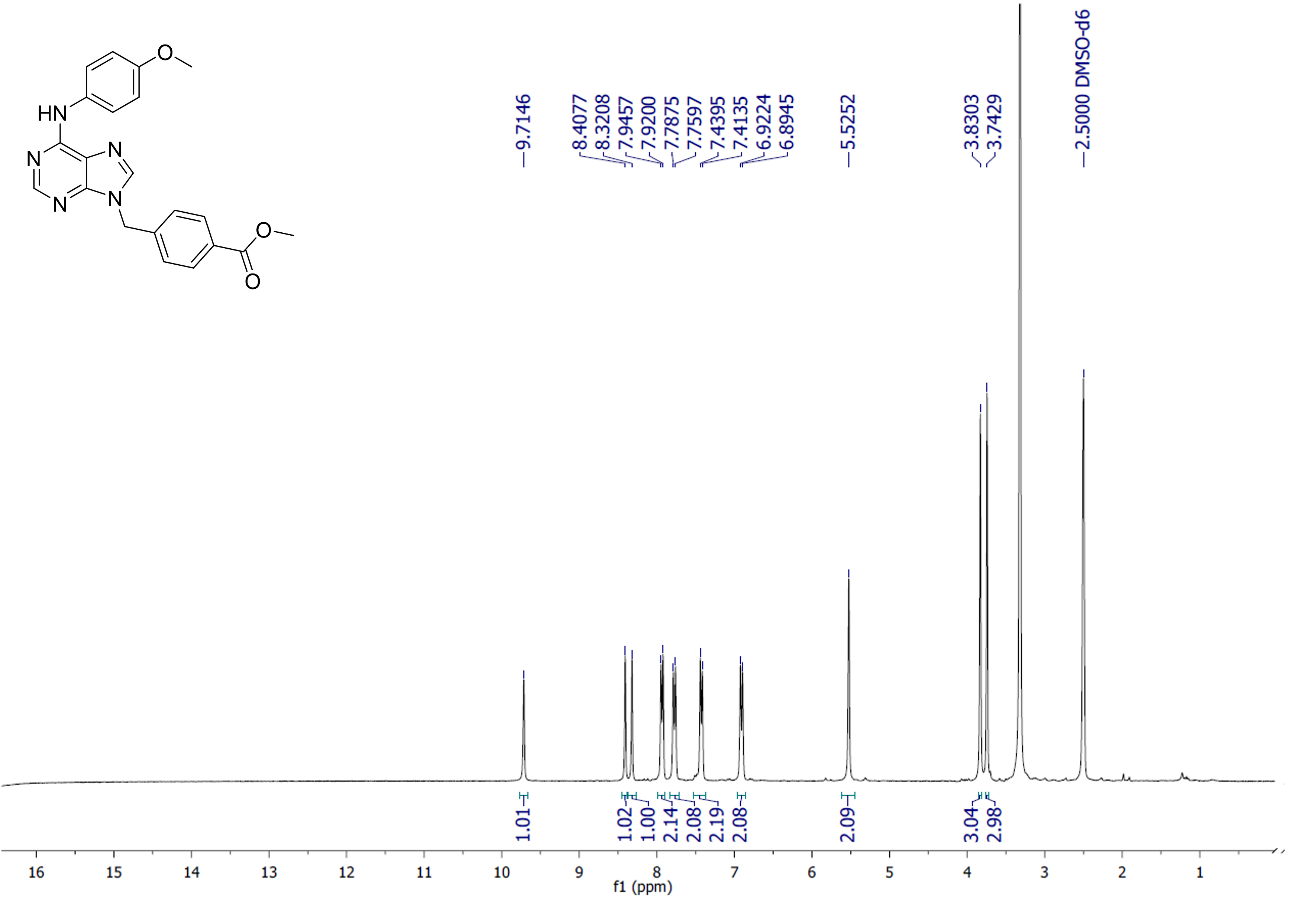

**Supplementary Figure S17:** ^13^C NMR spectra of the intermediate 3a (75 MHz, DMSO-d_6_, δ = ppm).

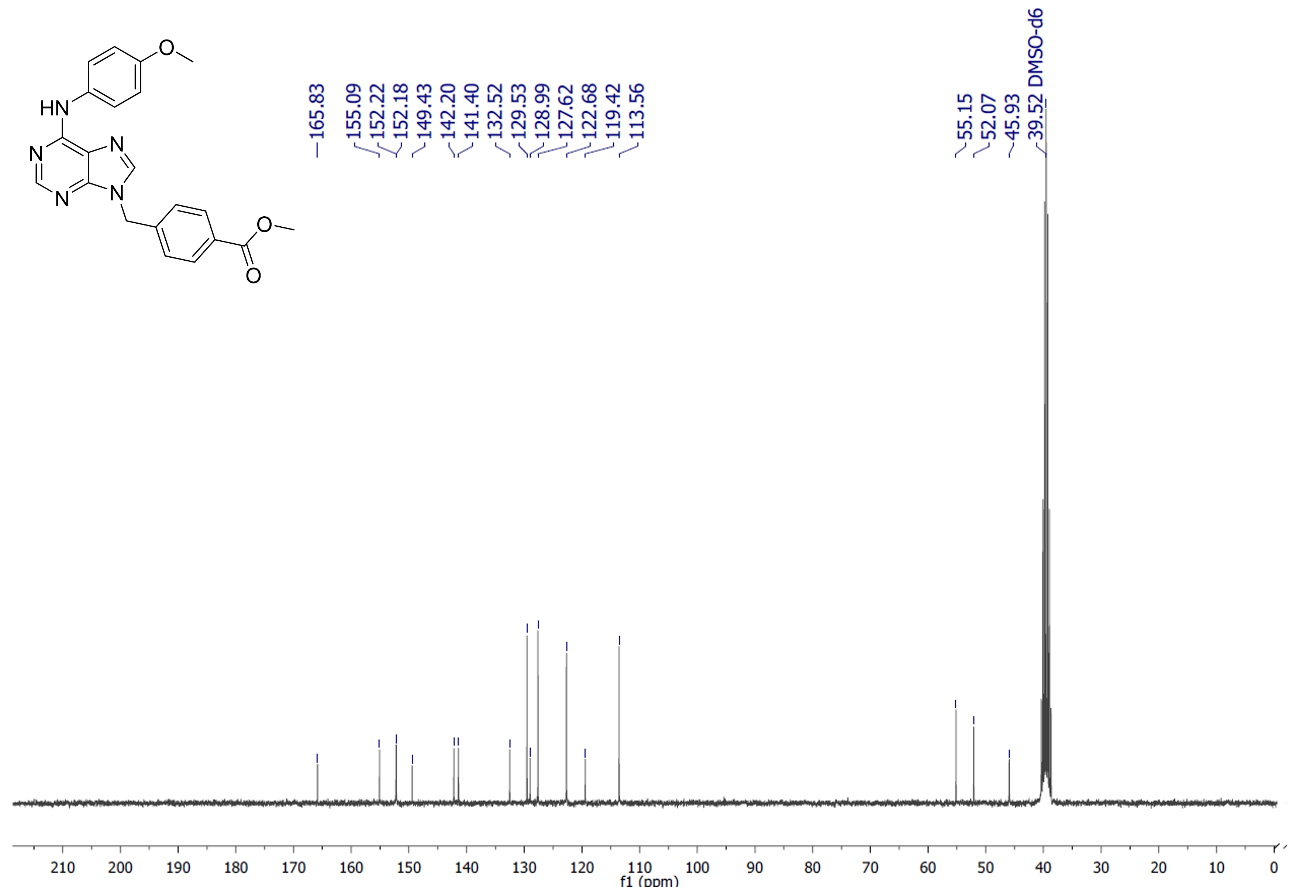

**Supplementary Figure S18:** ^1^H NMR spectra of the intermediate 3b (300 MHz, DMSO-d_6_, δ = ppm)

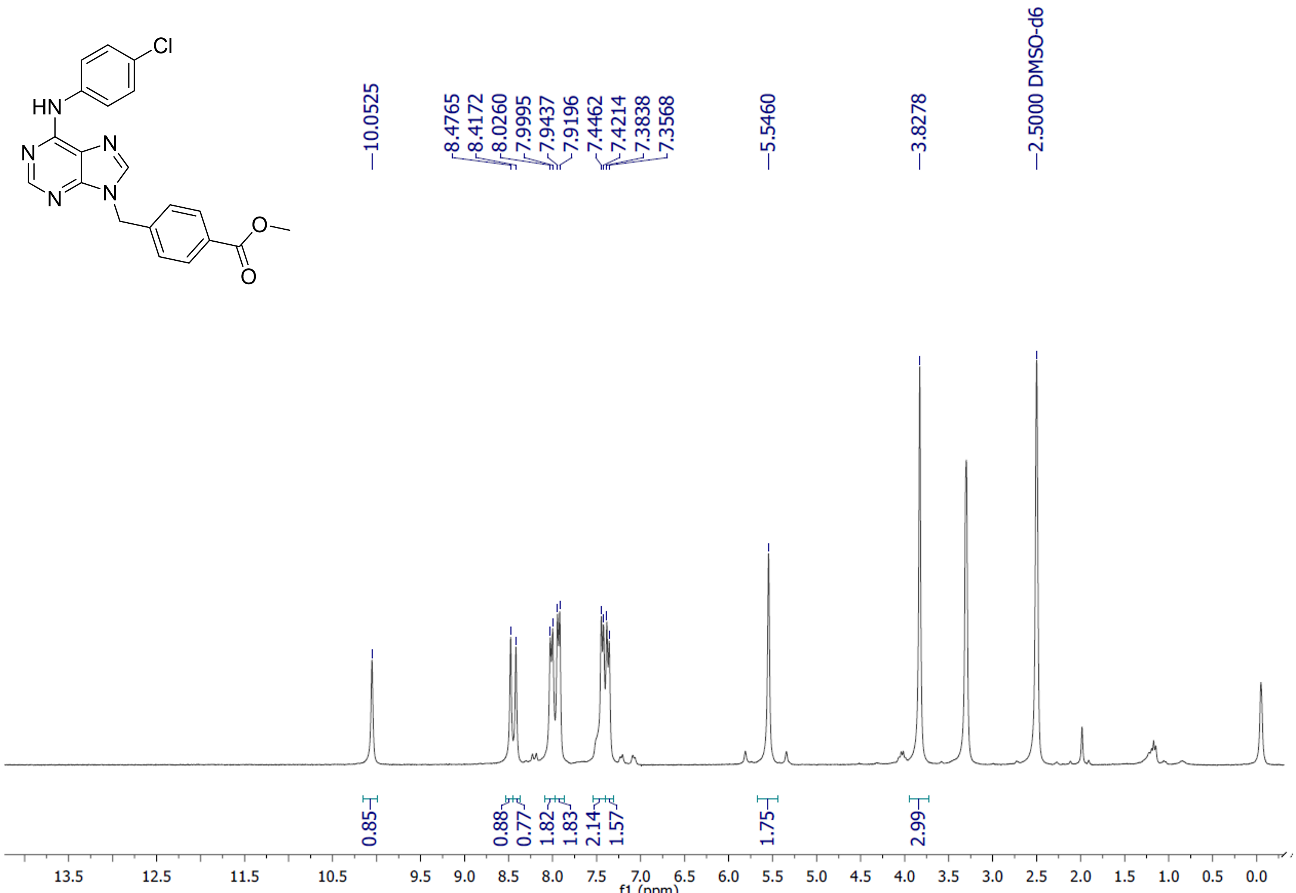

**Supplementary Figure S19:** ^13^C NMR spectra of the intermediate 3b (75 MHz, DMSO-d_6_, δ = ppm)

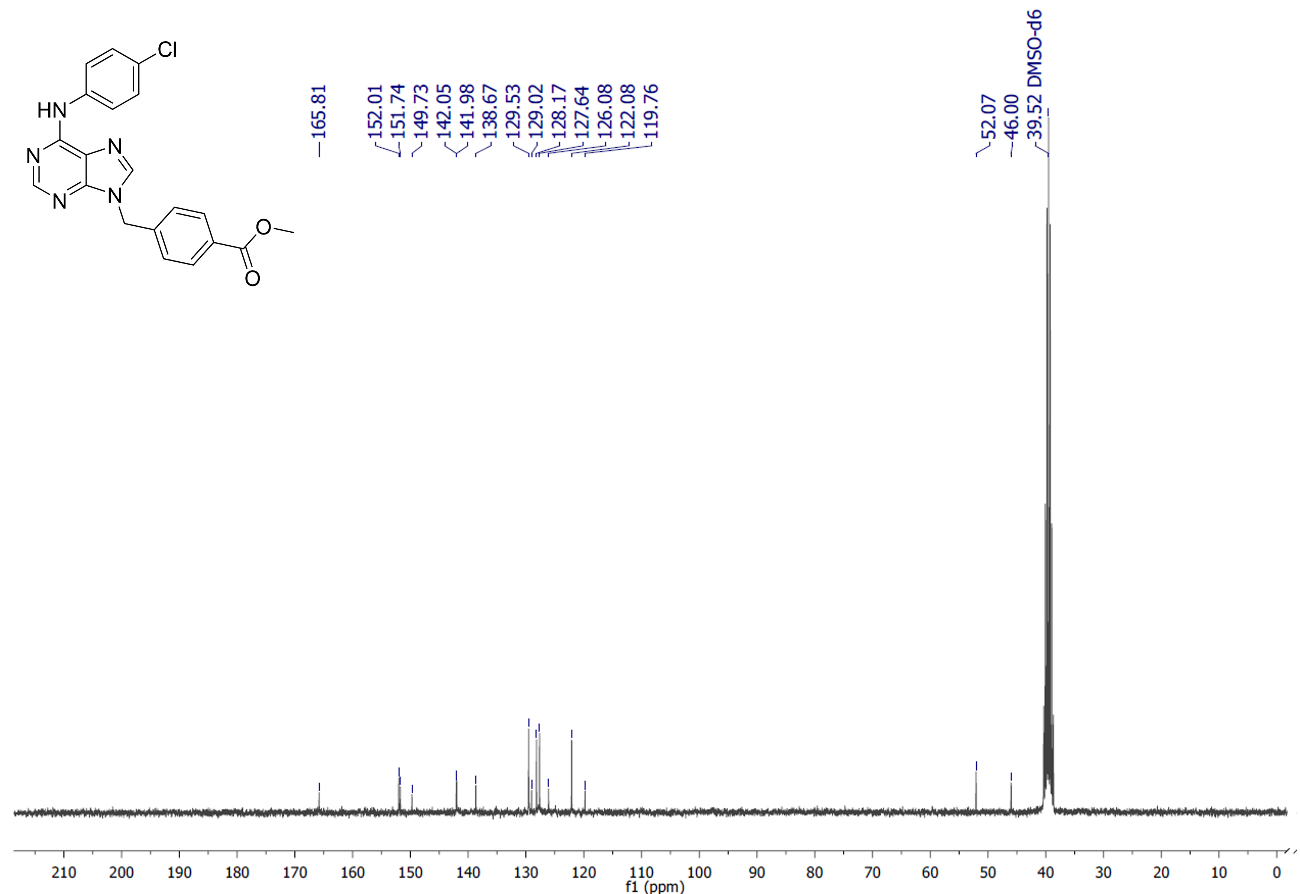

**Supplementary Figure S20:** ^1^H NMR spectra of the intermediate 3c (300 MHz, DMSO-d_6_, δ = ppm)

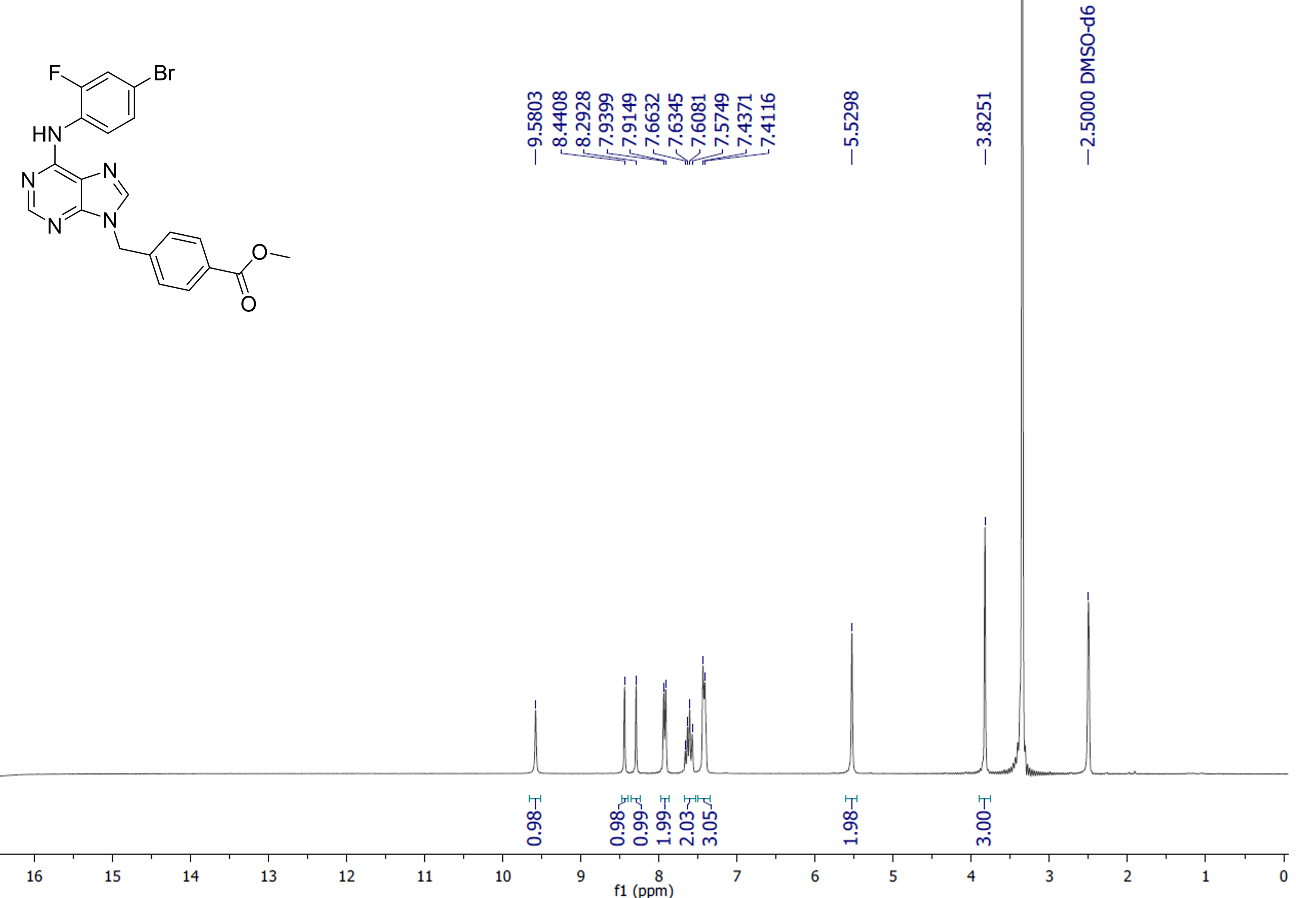

**Supplementary Figure S21:** ^13^C NMR spectra of the intermediate 3c (75 MHz, DMSO-d_6_, δ = ppm)

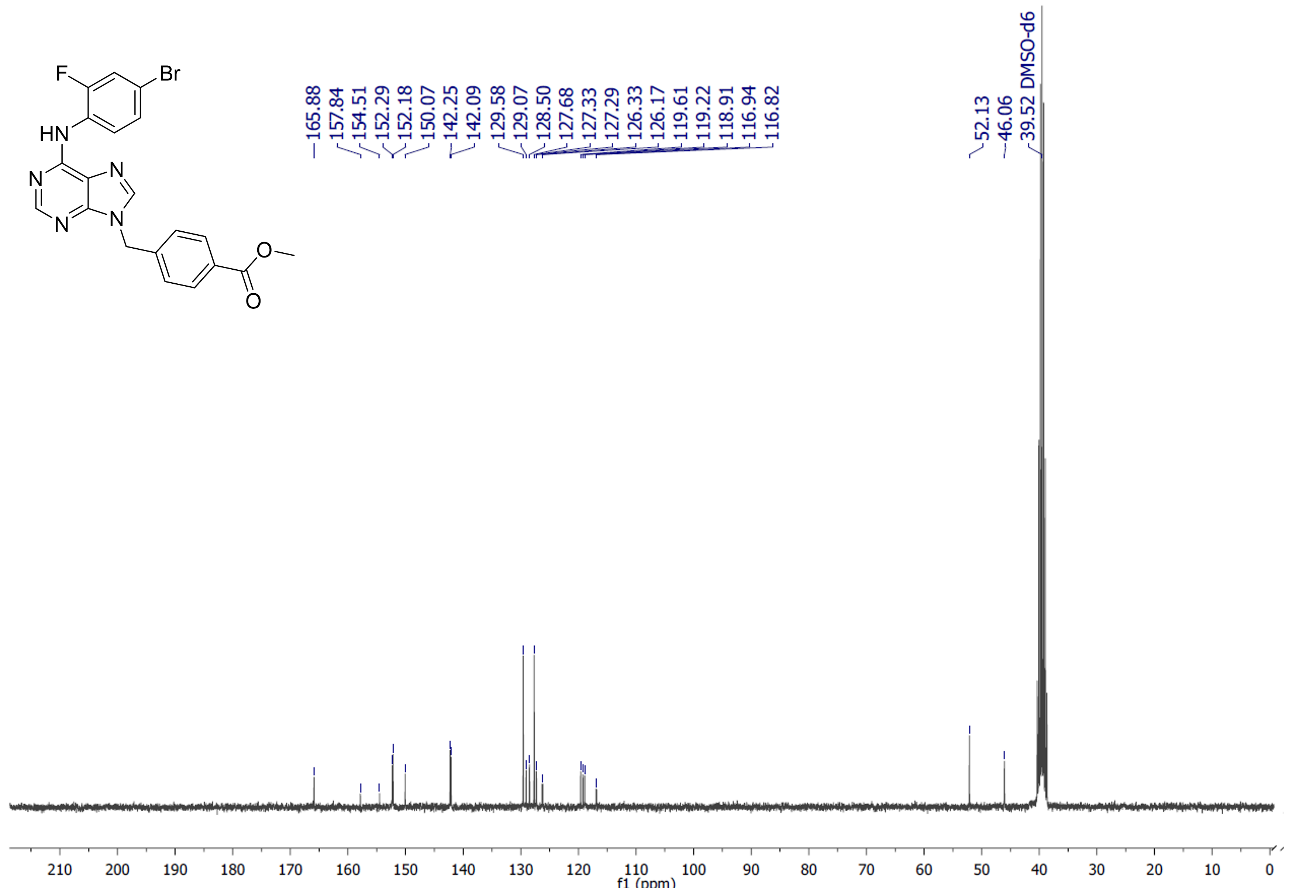

**Supplementary Figure S22:** ^1^H NMR spectra of the intermediate 3d (300 MHz, DMSO-d_6_, δ = ppm)

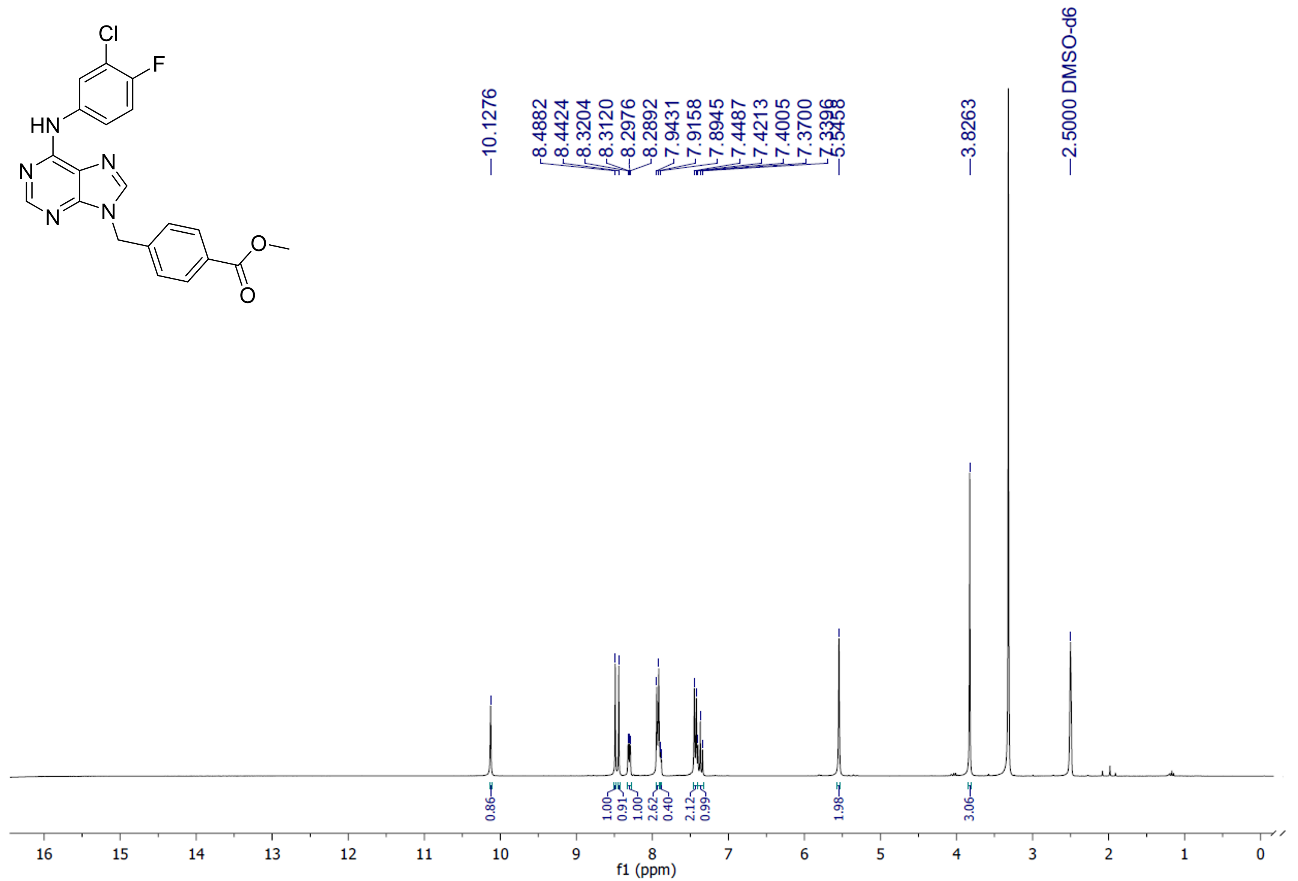

**Supplementary Figure S23:** ^13^C NMR spectra of the intermediate 3d (75 MHz, DMSO-d_6_, δ = ppm)

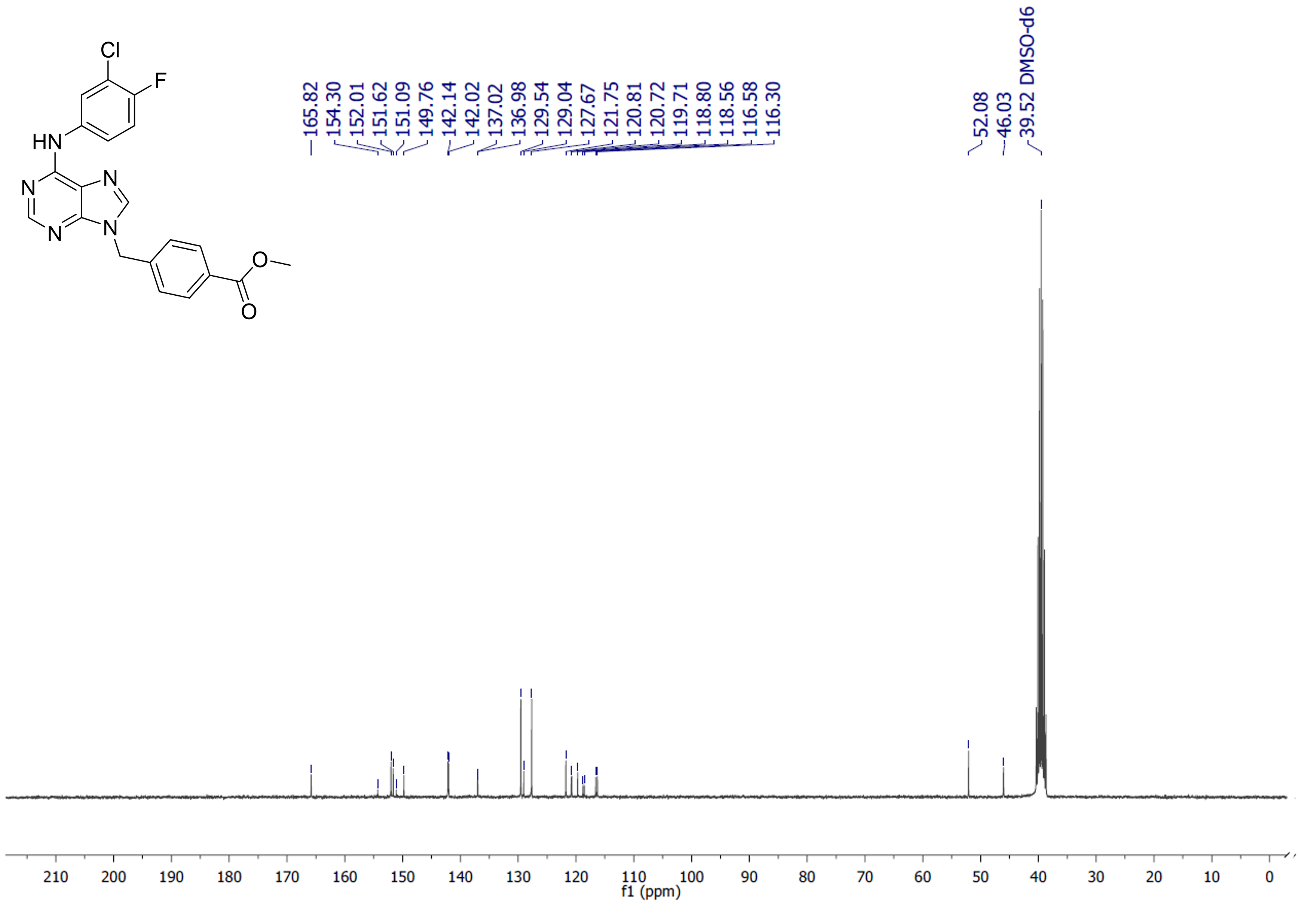

**Supplementary Figure S24:** ^1^H NMR spectra of the intermediate 3k (300 MHz, DMSO-d_6_, δ = ppm)

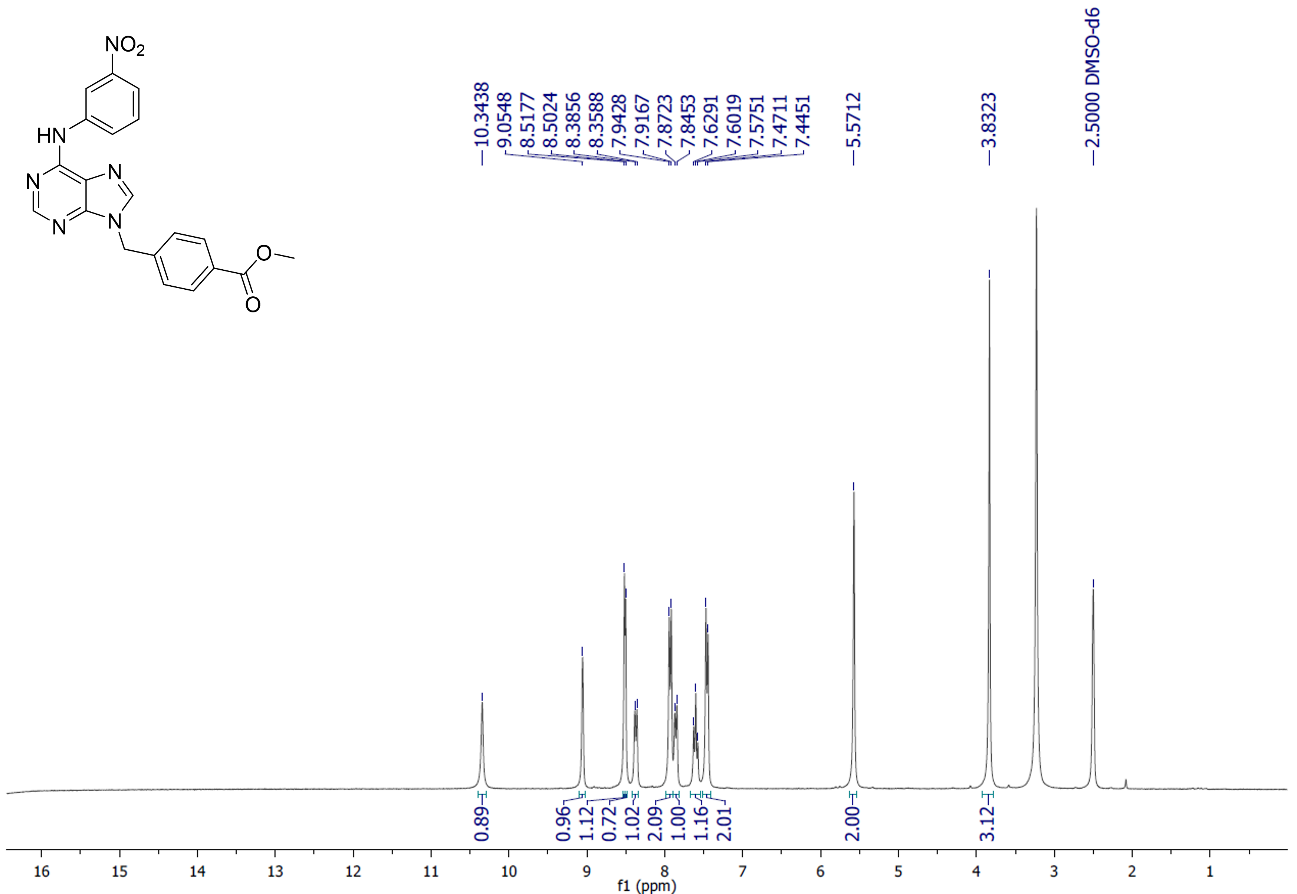

**Supplementary Figure S25:** ^13^C NMR spectra of the intermediate 3k (75 MHz, DMSO-d_6_, δ = ppm)

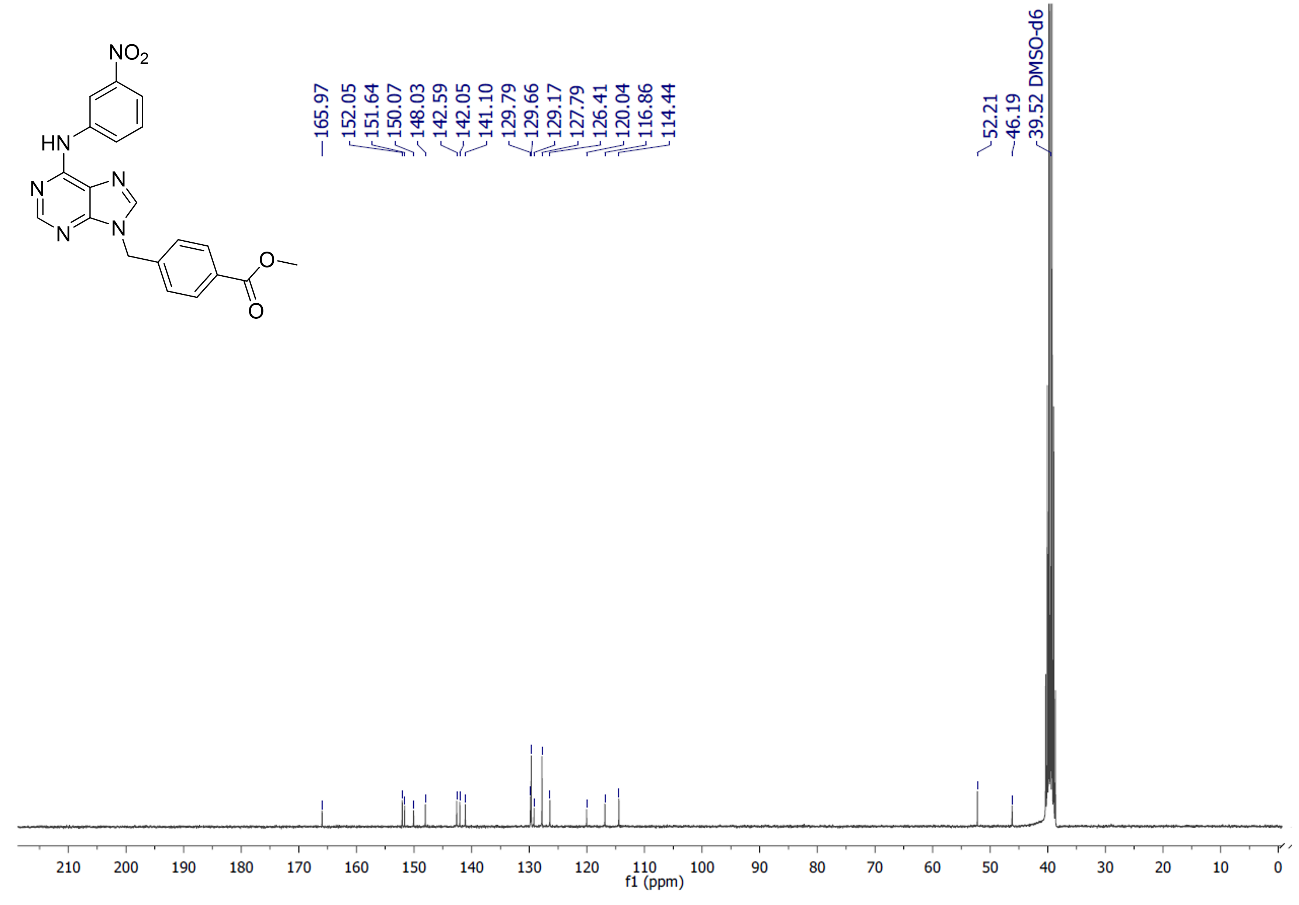

**Supplementary Figure S26:** ^1^H NMR spectra of the final compound 4a (300 MHz, DMSO-d_6_, δ = ppm)

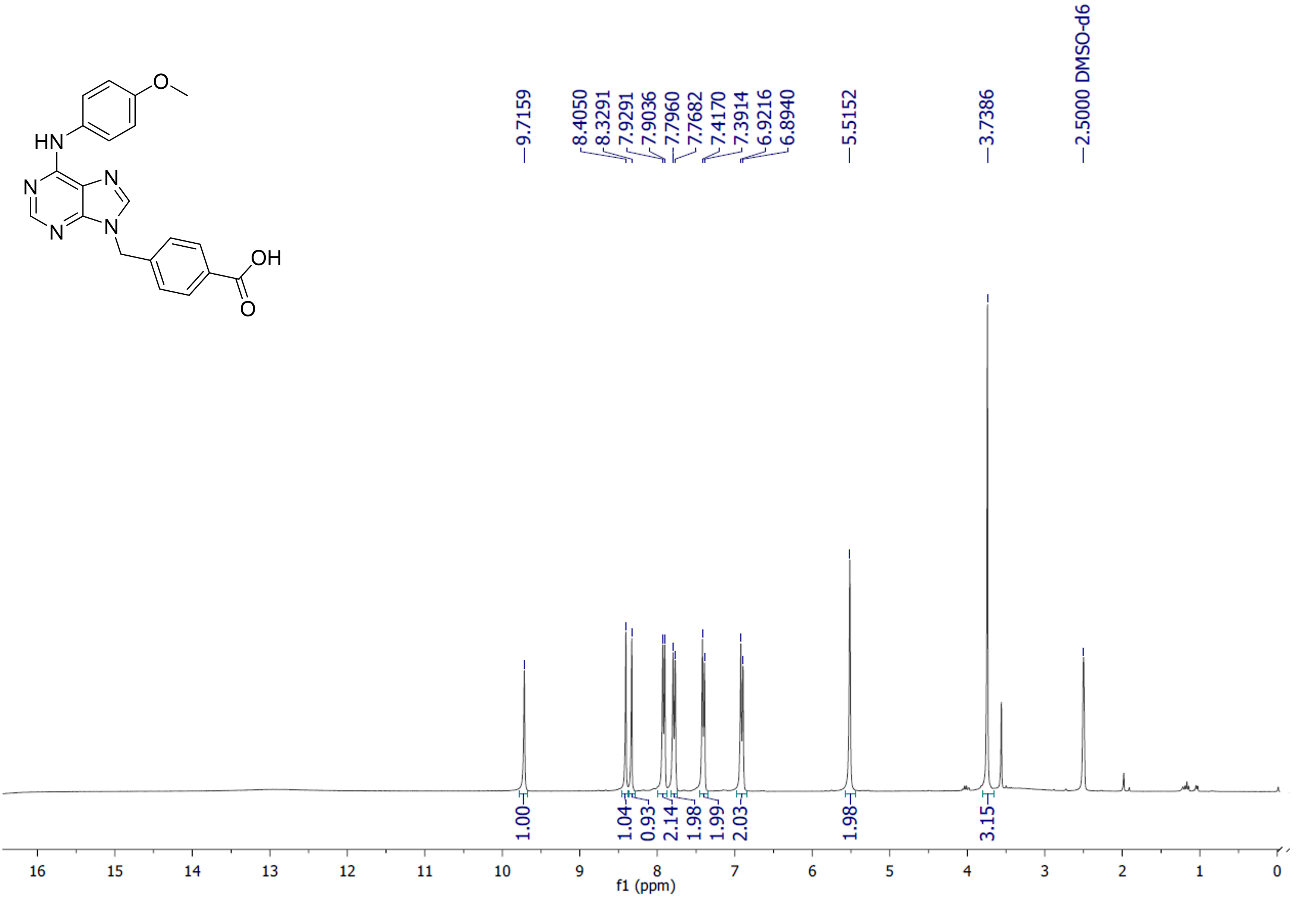

**Supplementary Figure S27:** ^13^C NMR spectra of the intermediate 4a (75 MHz, DMSO-d_6_, δ = ppm)

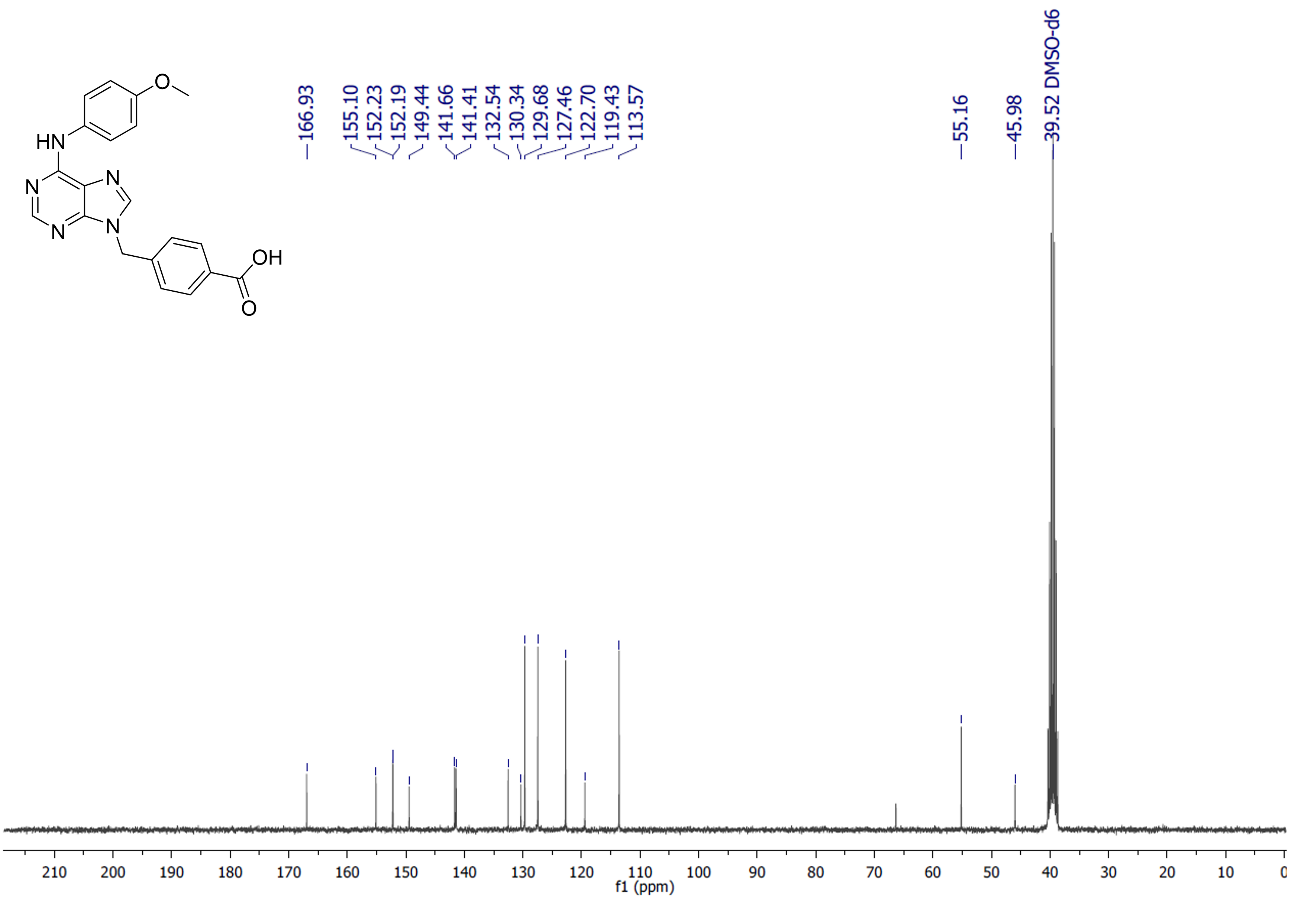

**Supplementary Figure S28:** ^1^H NMR spectra of the intermediate 4b (300 MHz, DMSO-d_6_, δ = ppm)

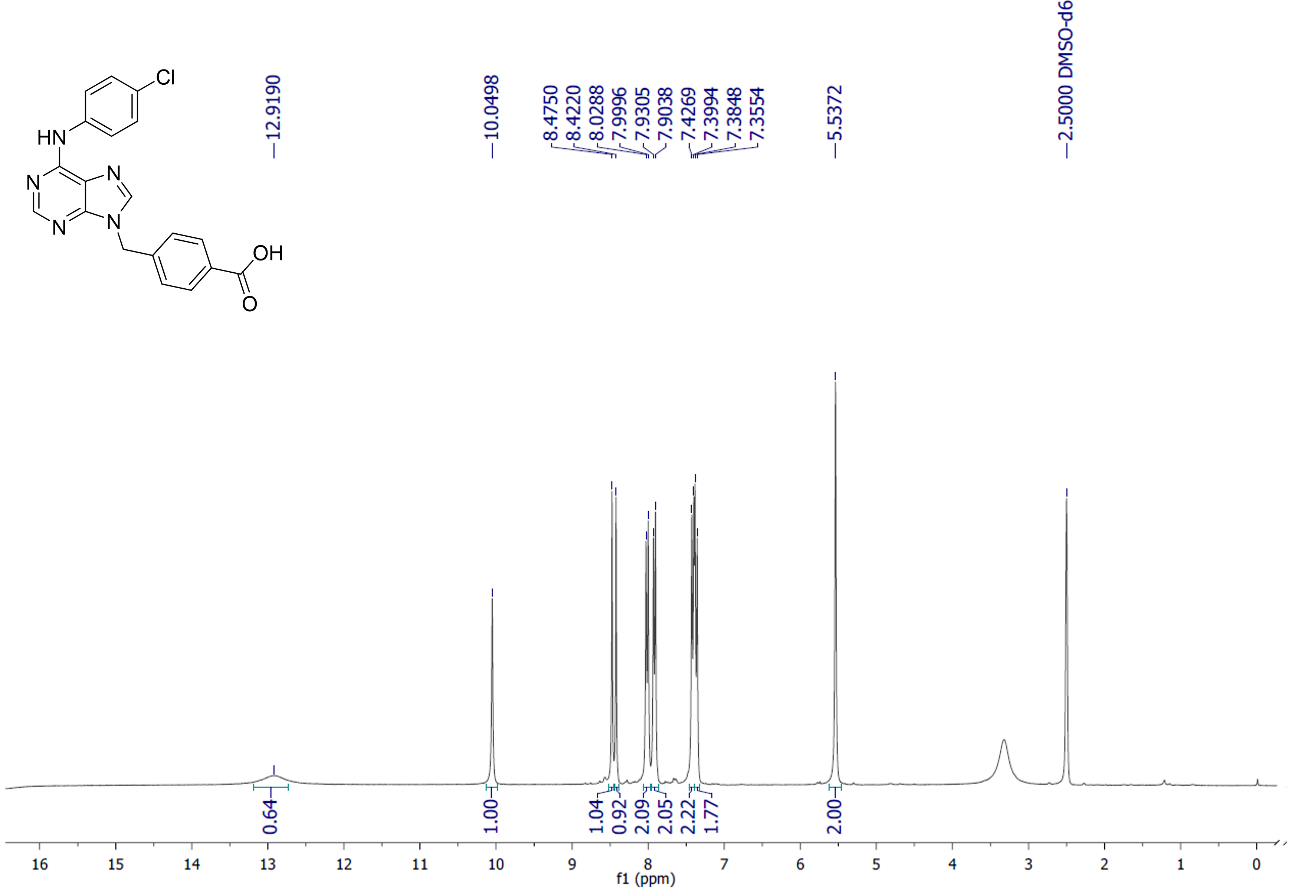

**Supplementary Figure S29:** ^13^C NMR spectra of the intermediate 4b (75 MHz, DMSO-d_6_, δ = ppm)

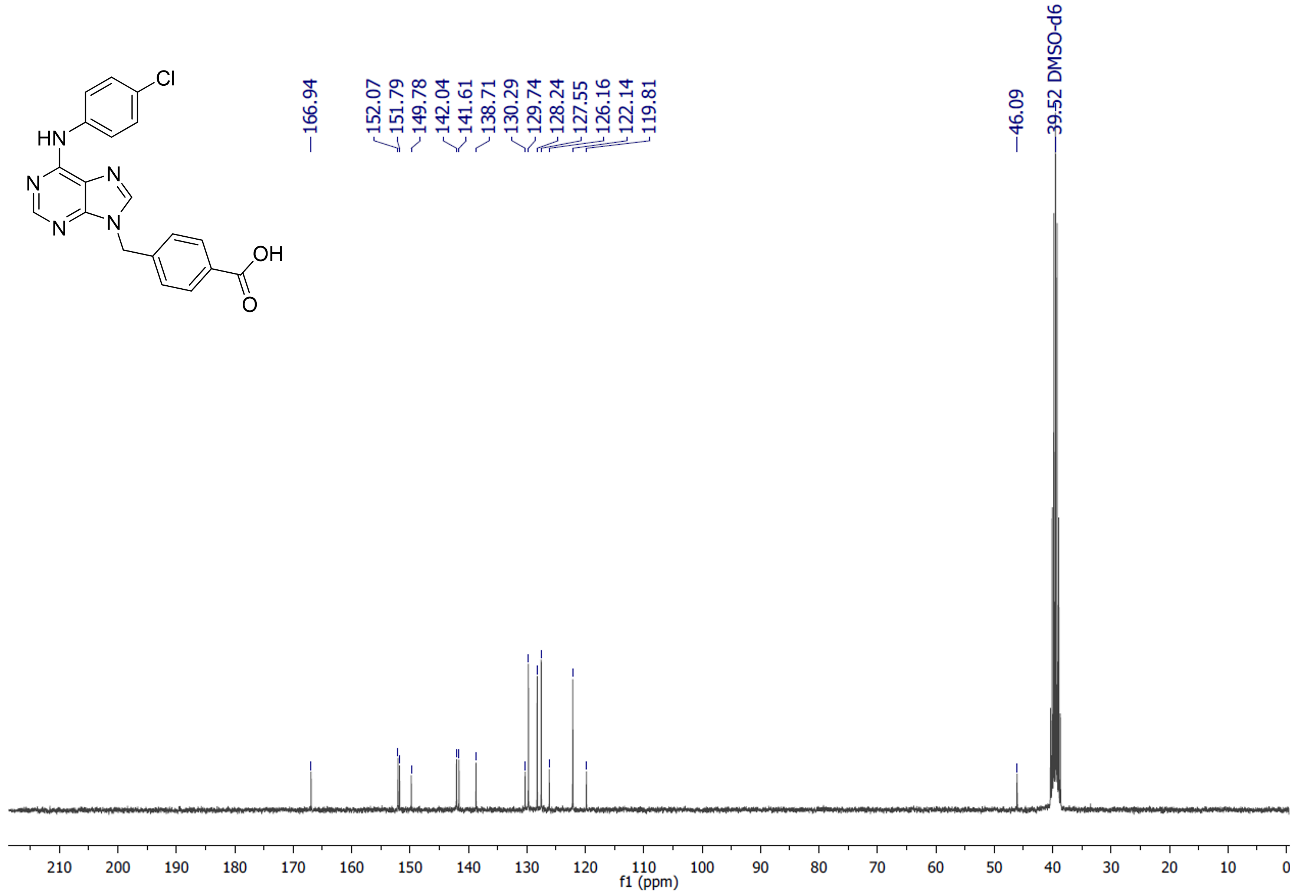

**Supplementary Figure S30:** ^1^H NMR spectra of the intermediate 4c (300 MHz, DMSO-d_6_, δ = ppm)

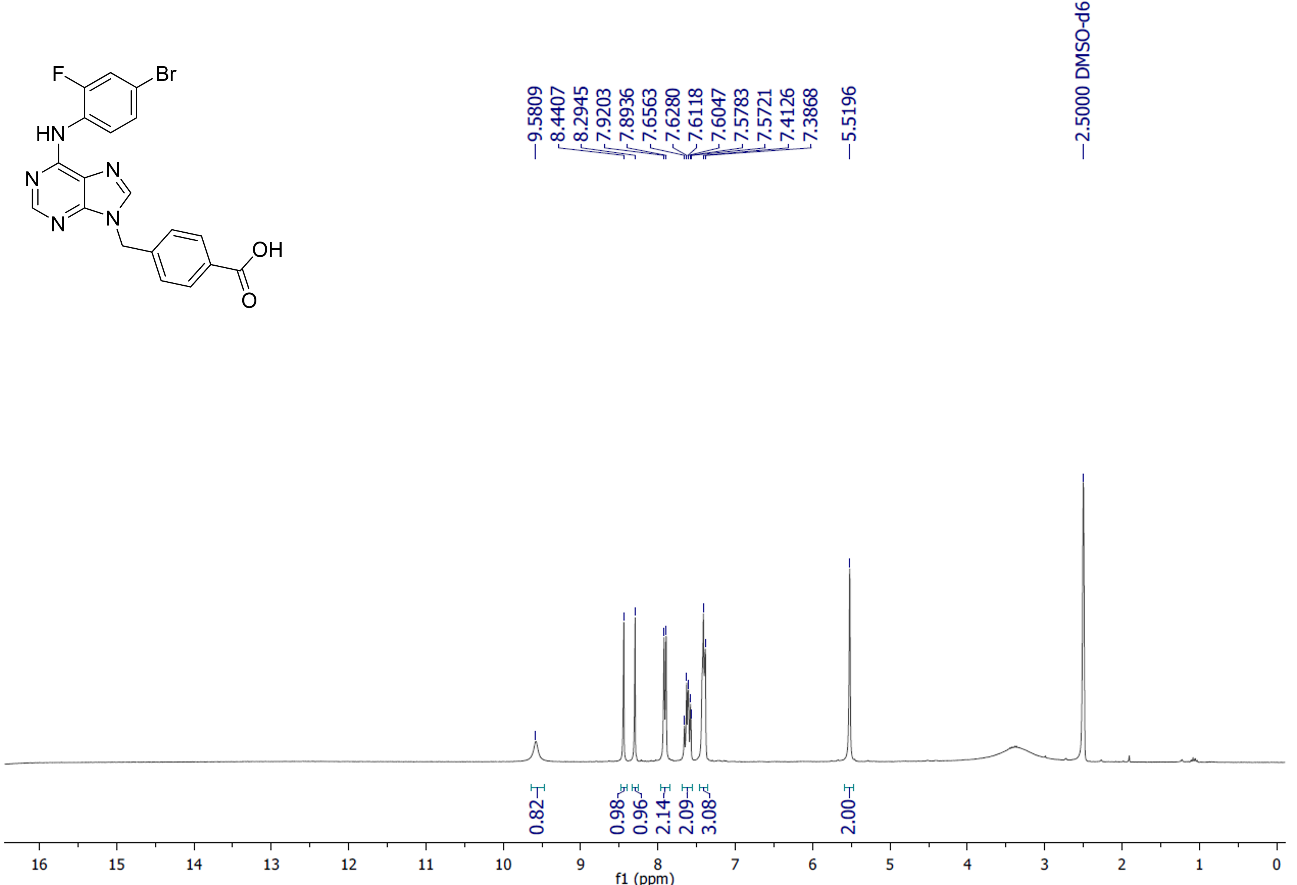

**Supplementary Figure S31:** ^13^C NMR spectra of the intermediate 4c (75 MHz, DMSO-d_6_, δ = ppm)

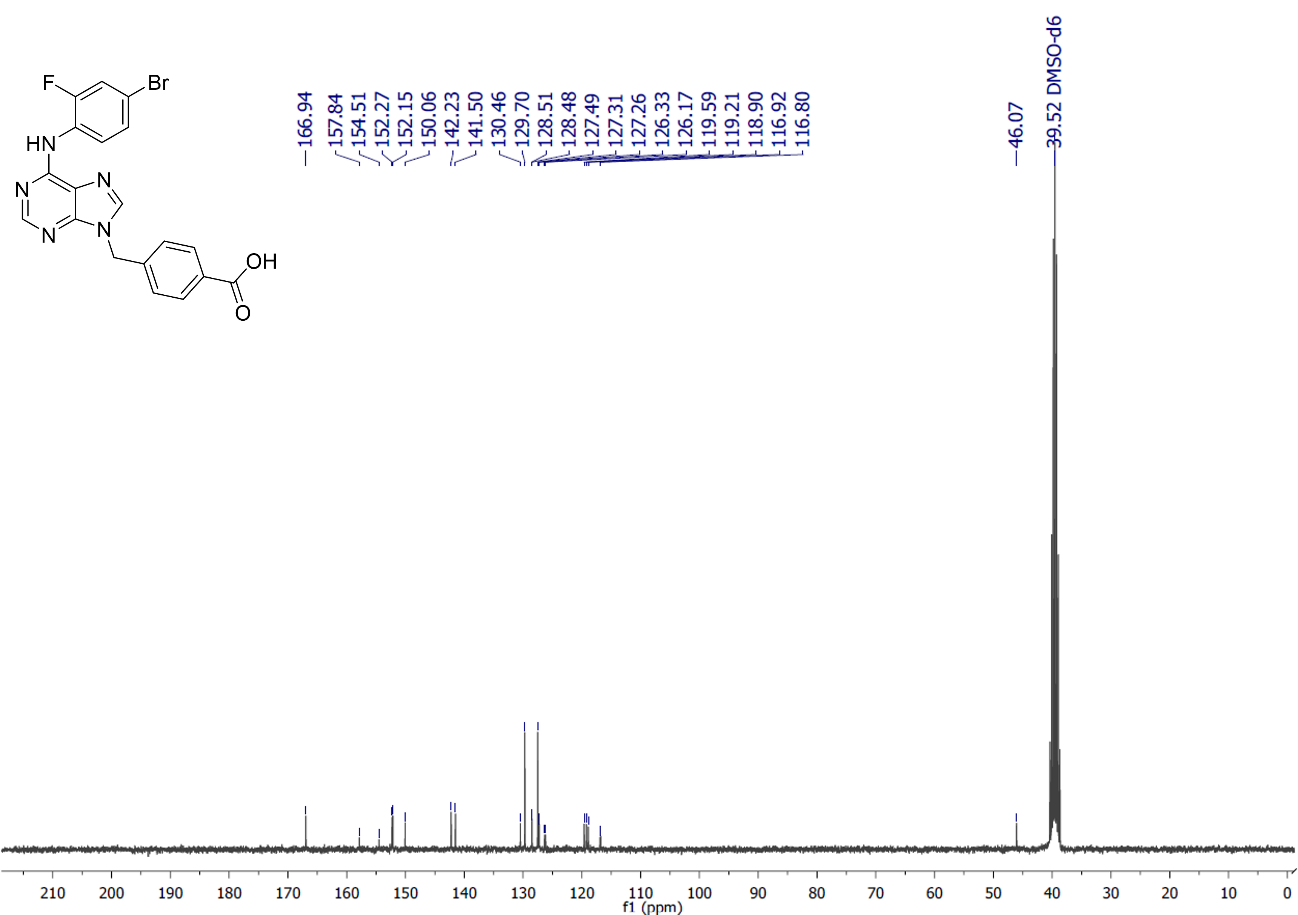

**Supplementary Figure S32:** ^1^H NMR spectra of the final compound 5a (300 MHz, DMSO-d_6_, δ = ppm)

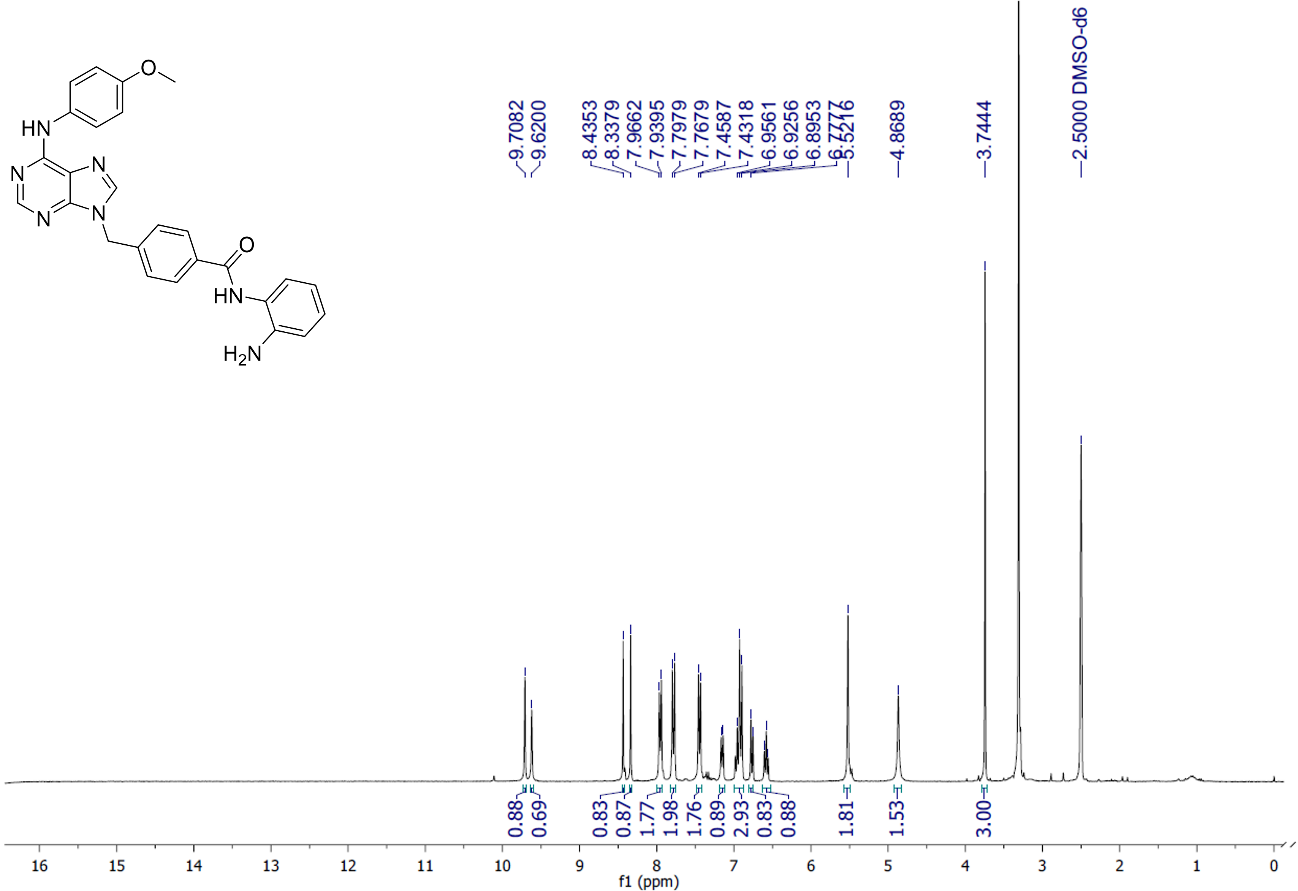

**Supplementary Figure S33:** ^13^C NMR spectra of the intermediate 5a (75 MHz, DMSO-d_6_, δ = ppm)

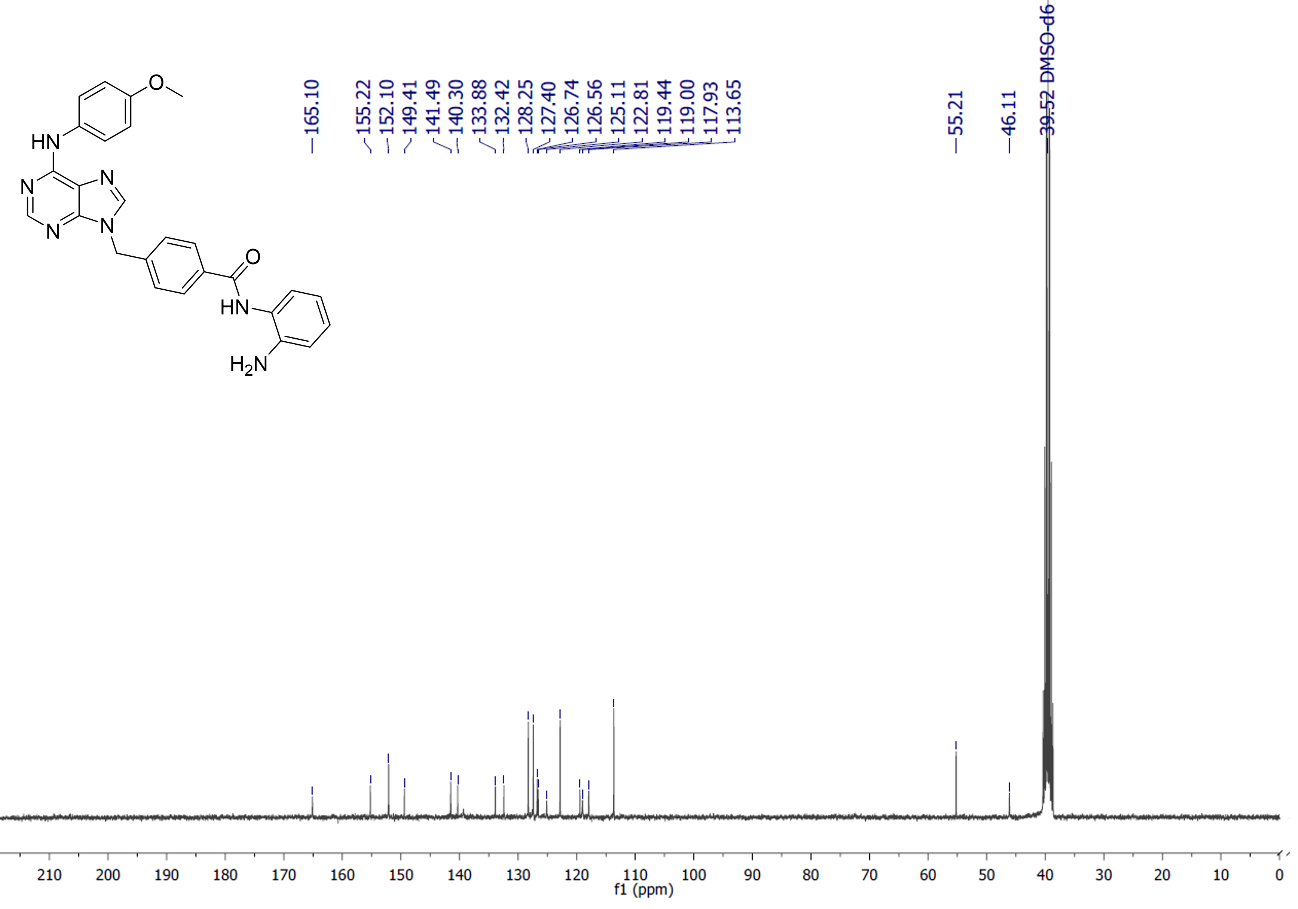

**Supplementary Figure S34:** ^1^H NMR spectra of the final compound 5b (300 MHz, DMSO-d_6_, δ = ppm)

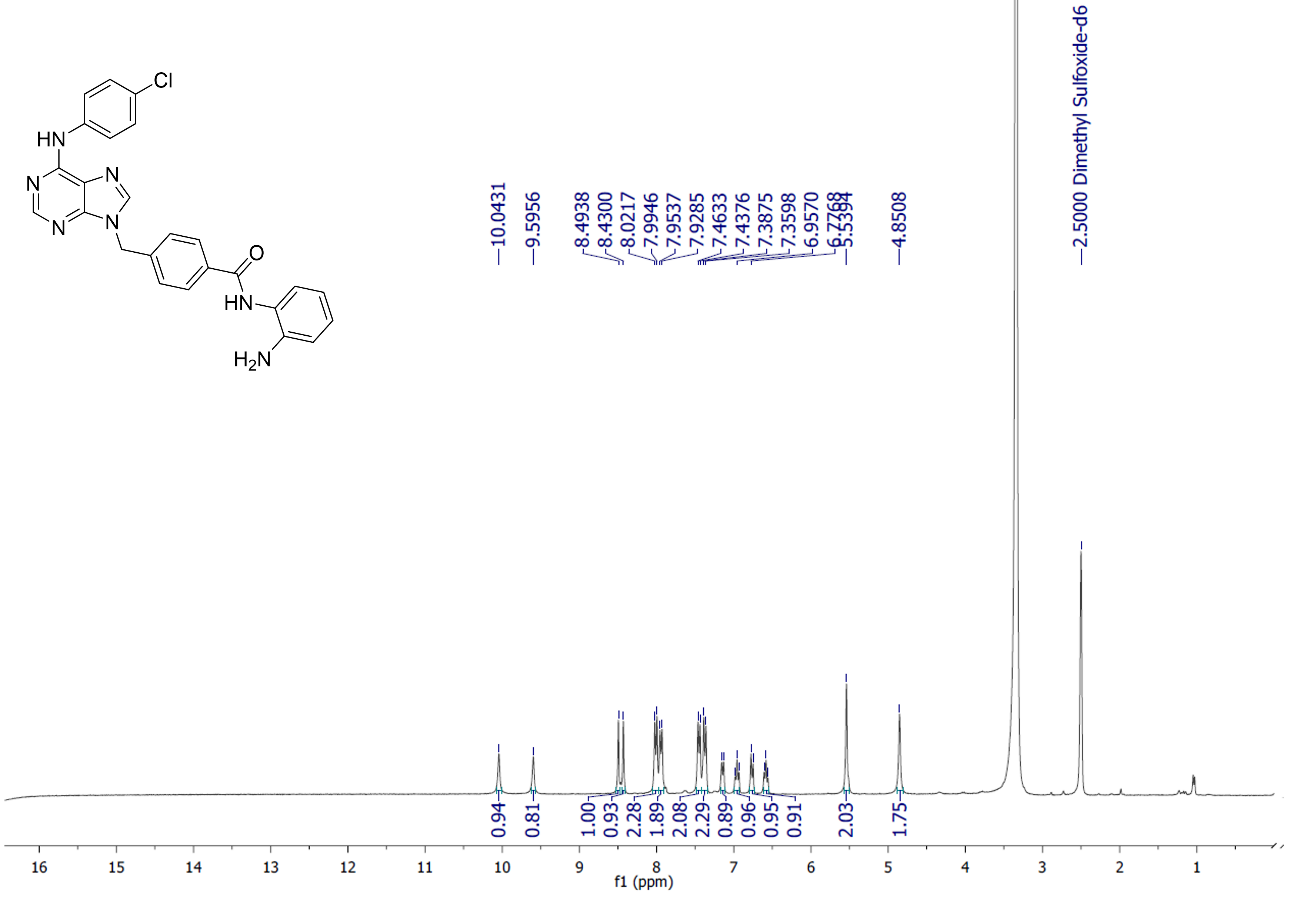

**Supplementary Figure S35:** ^13^C NMR spectra of the final compound 5b (75 MHz, DMSO-d_6_, δ = ppm)

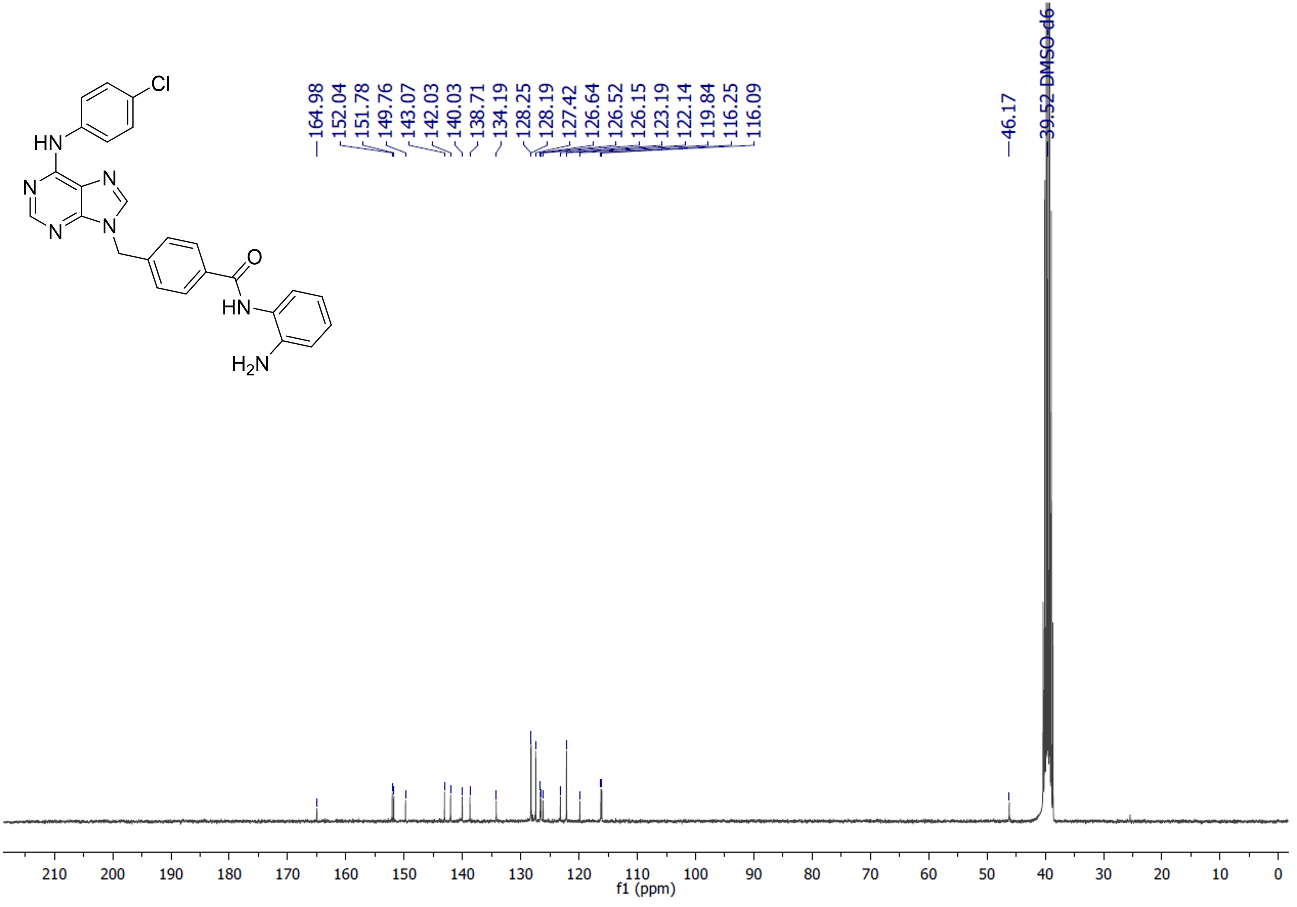

**Supplementary Figure S36:** ^1^H NMR spectra of the final compound 5c (300 MHz, DMSO-d_6_, δ = ppm)

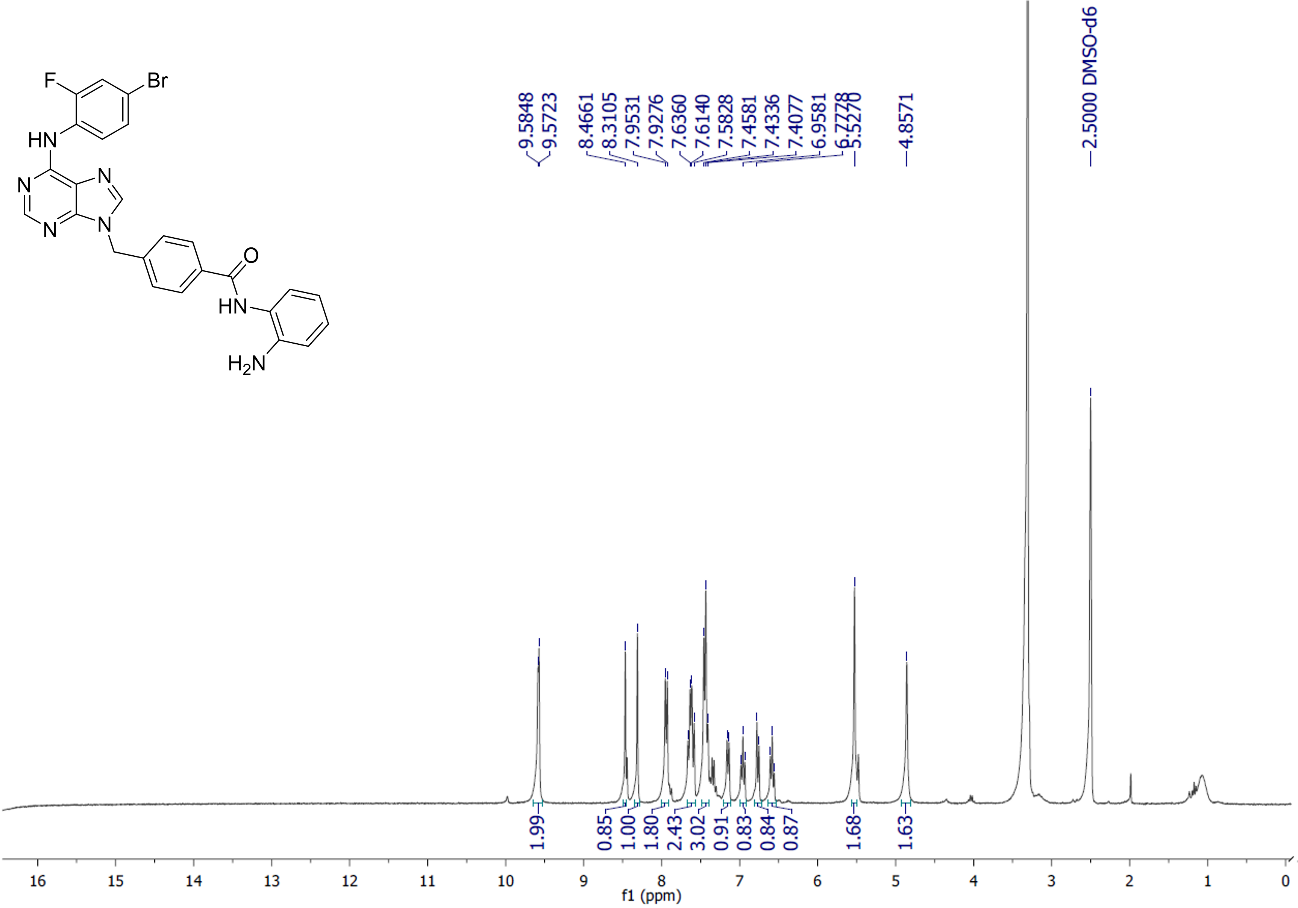

**Supplementary Figure S37:** ^13^C NMR spectra of the intermediate 5c (75 MHz, DMSO-d_6_, δ = ppm)

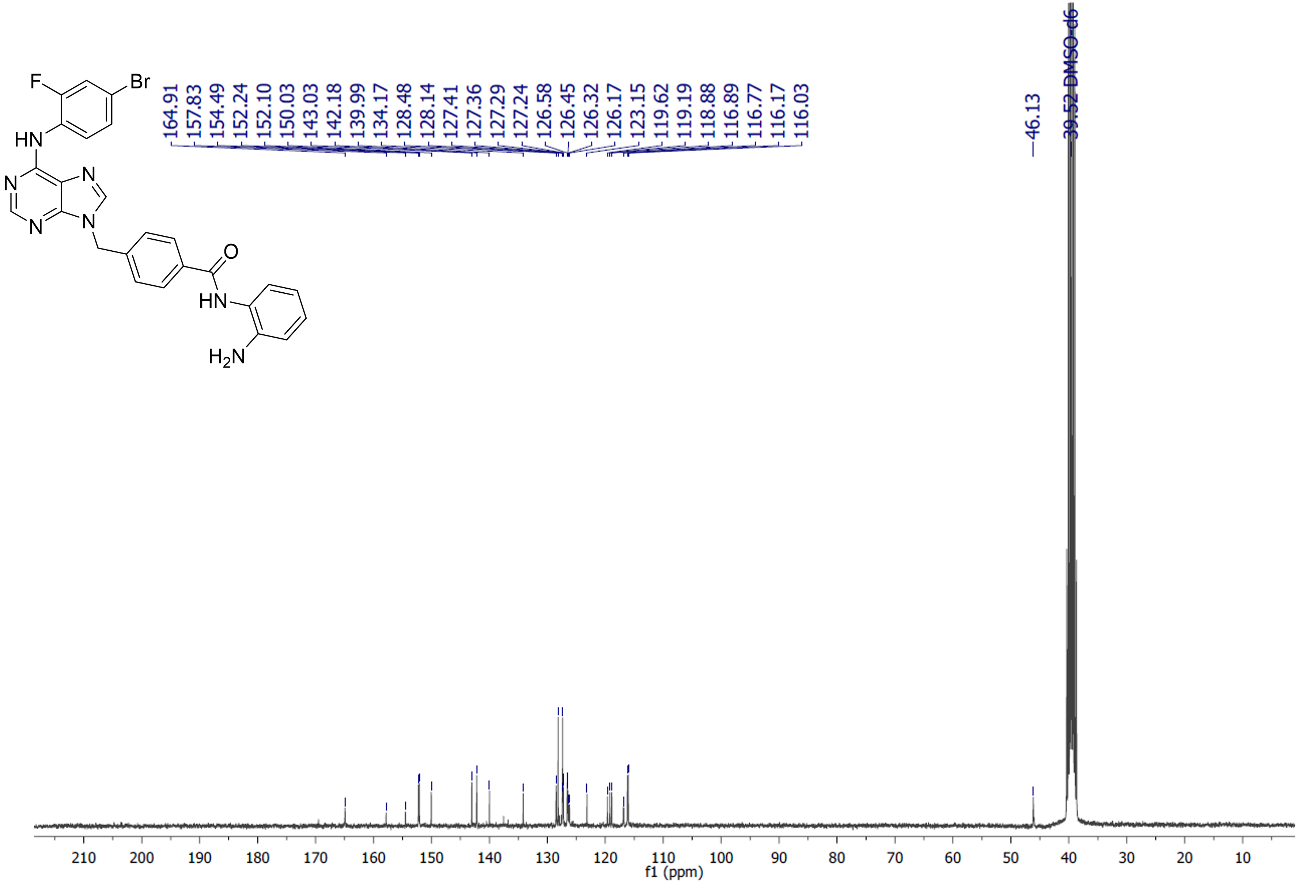

**Supplementary Figure S38:** ^1^H NMR spectra of the final compound 6a (300 MHz, DMSO-d_6_, δ = ppm)

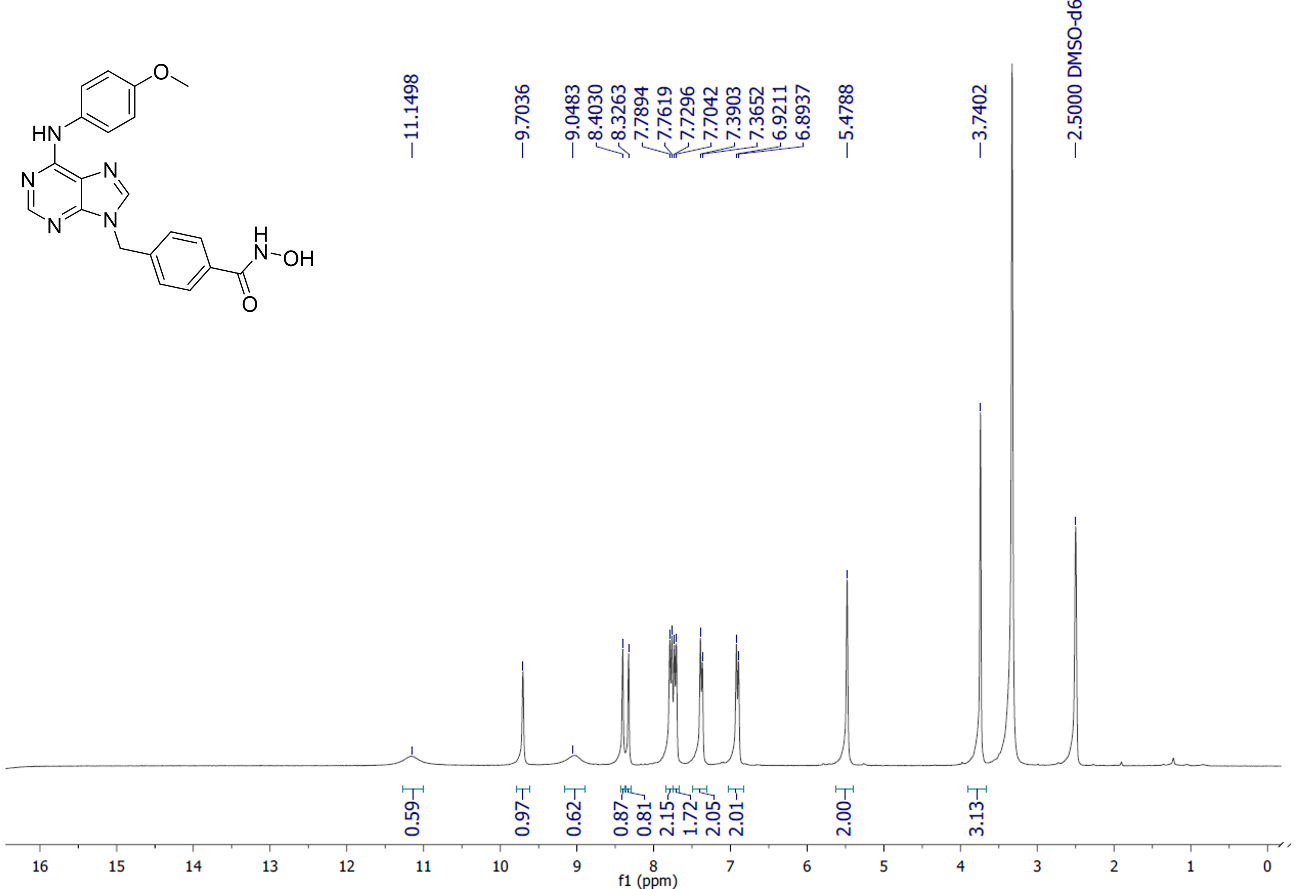

**Supplementary Figure S39:** ^13^C NMR spectra of the final compound 6a (75 MHz, DMSO-d_6_, δ = ppm)

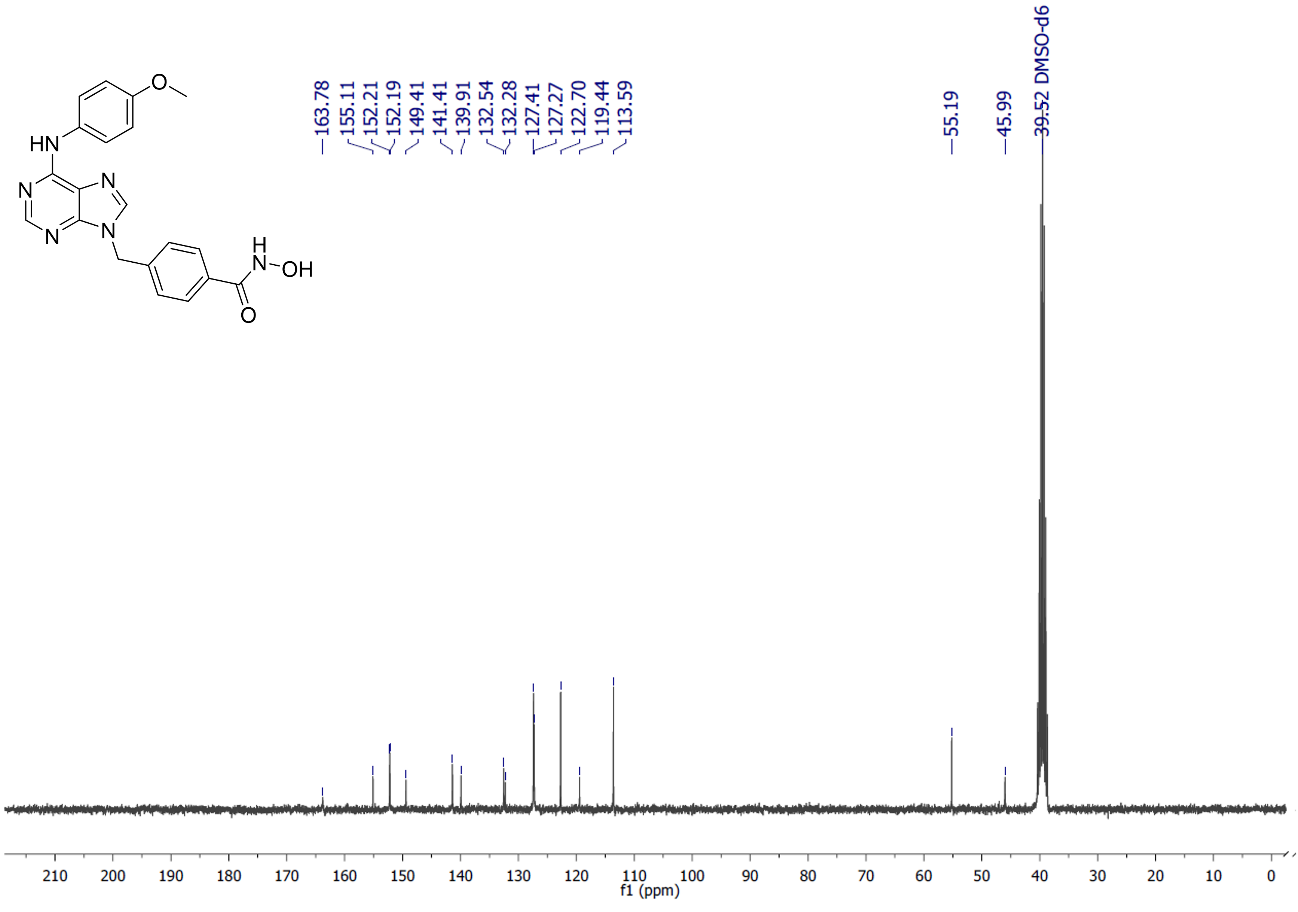

**Supplementary Figure S40:** ^1^H NMR spectra of the final compound 6b (300 MHz, DMSO-d_6_, δ = ppm)

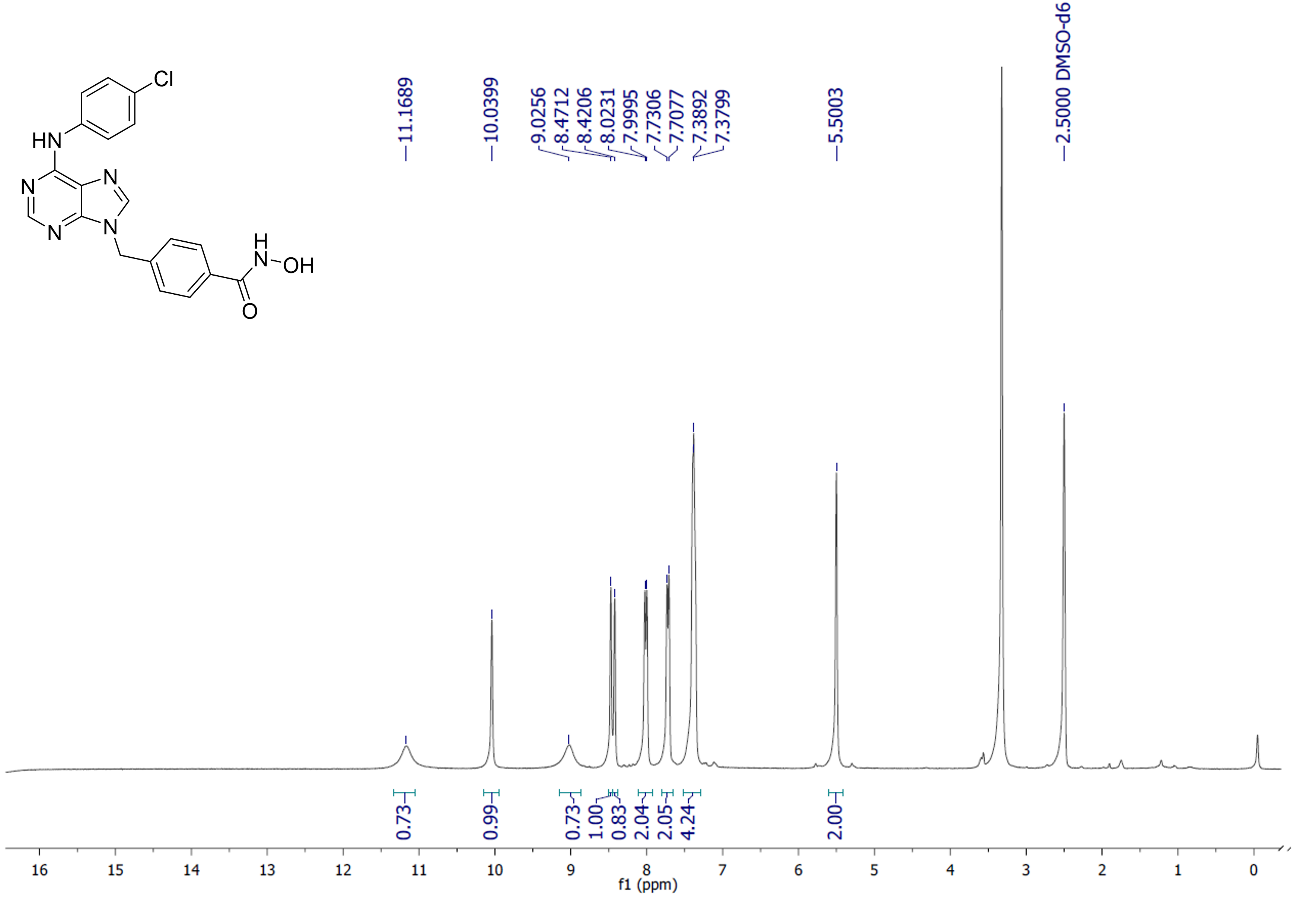

**Supplementary Figure S41:** ^13^C NMR spectra of the final compound 6b (75 MHz, DMSO-d_6_, δ = ppm)

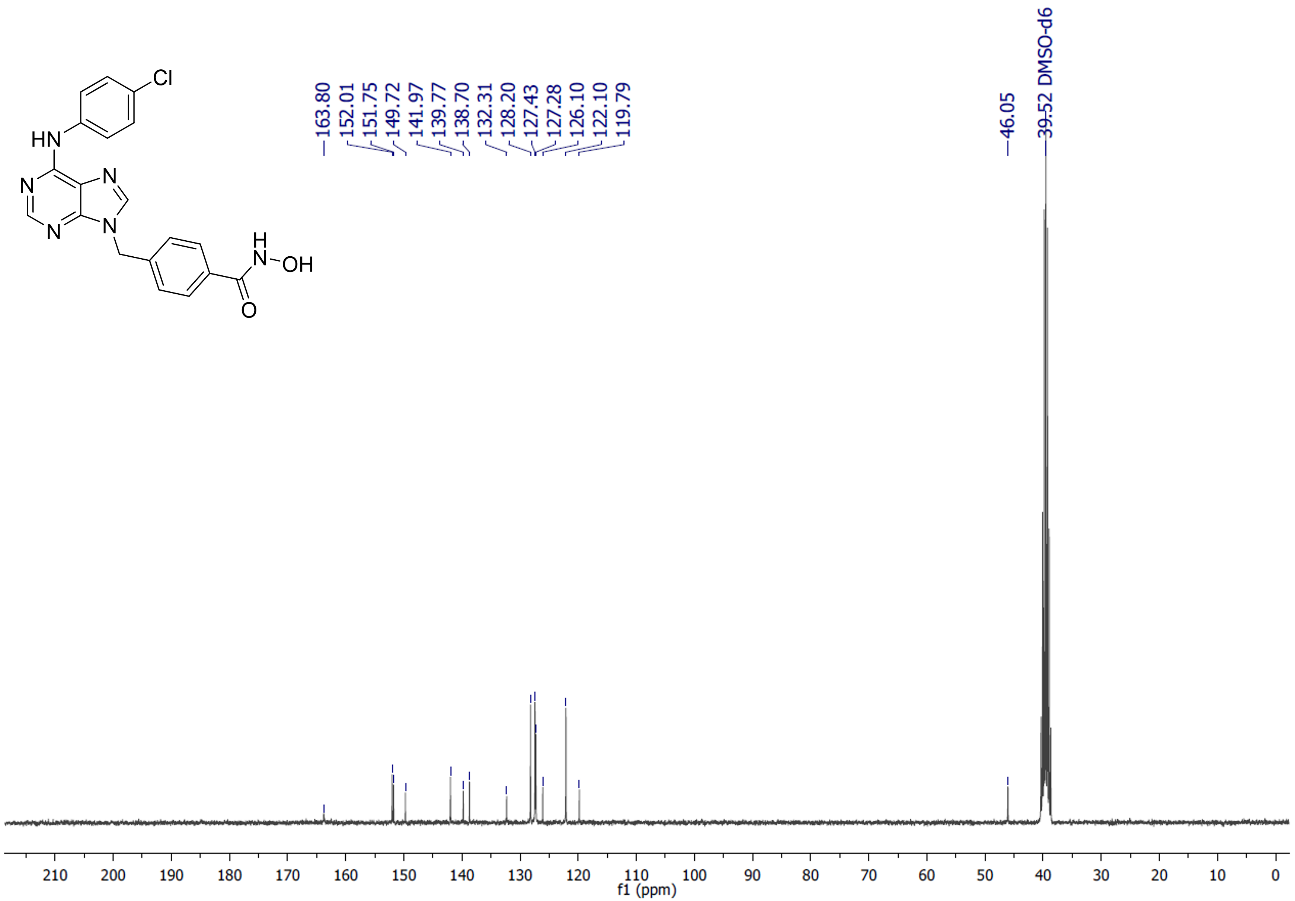

**Supplementary Figure S42:** ^1^H NMR spectra of the final compound 6c (300 MHz, DMSO-d_6_, δ = ppm)

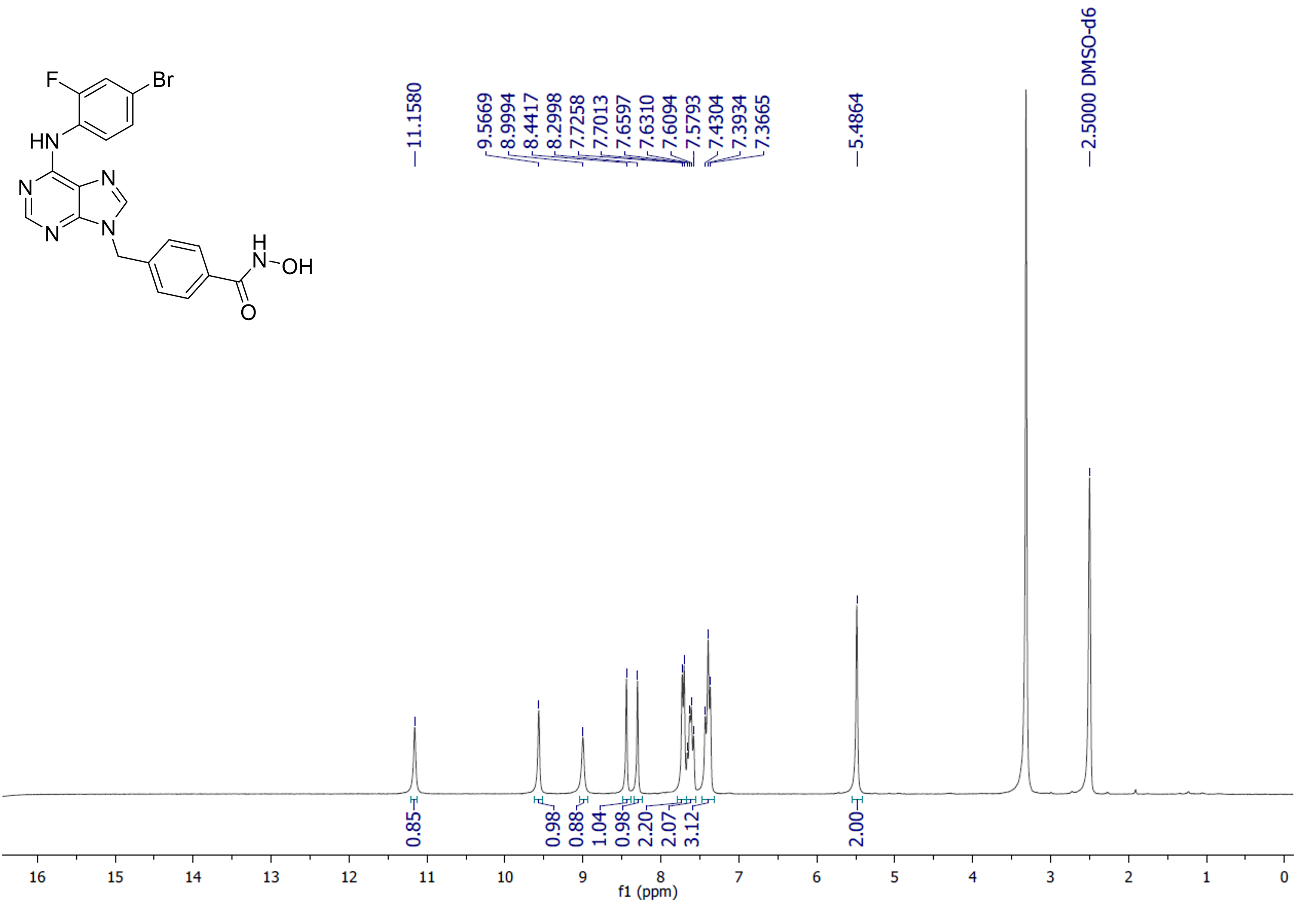

**Supplementary Figure S43:** ^13^C NMR spectra of the final compound 6c (75 MHz, DMSO-d_6_, δ = ppm)

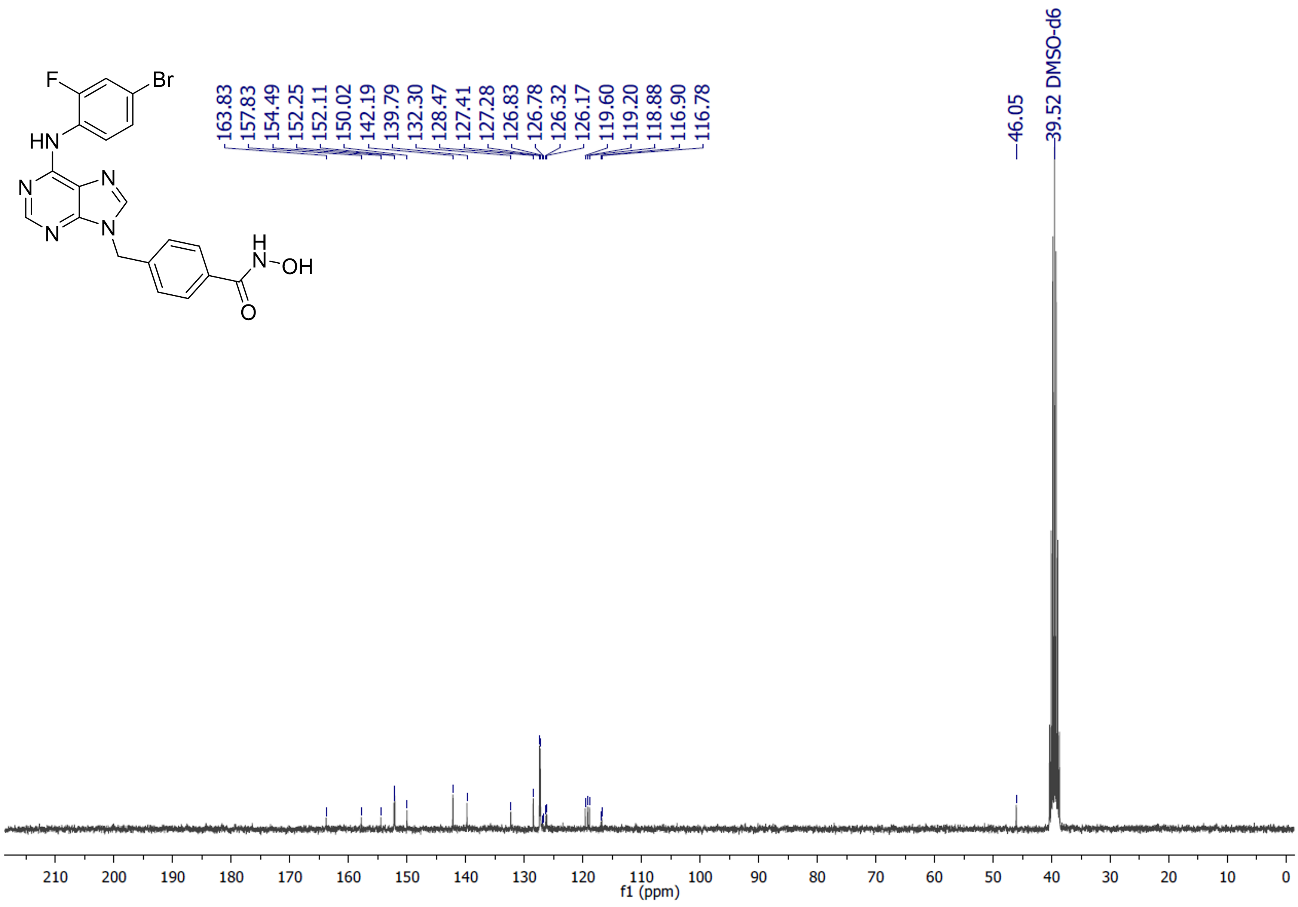

**Supplementary Figure S44:** ^1^H NMR spectra of the final compound 6d (300 MHz, DMSO-d_6_, δ = ppm)

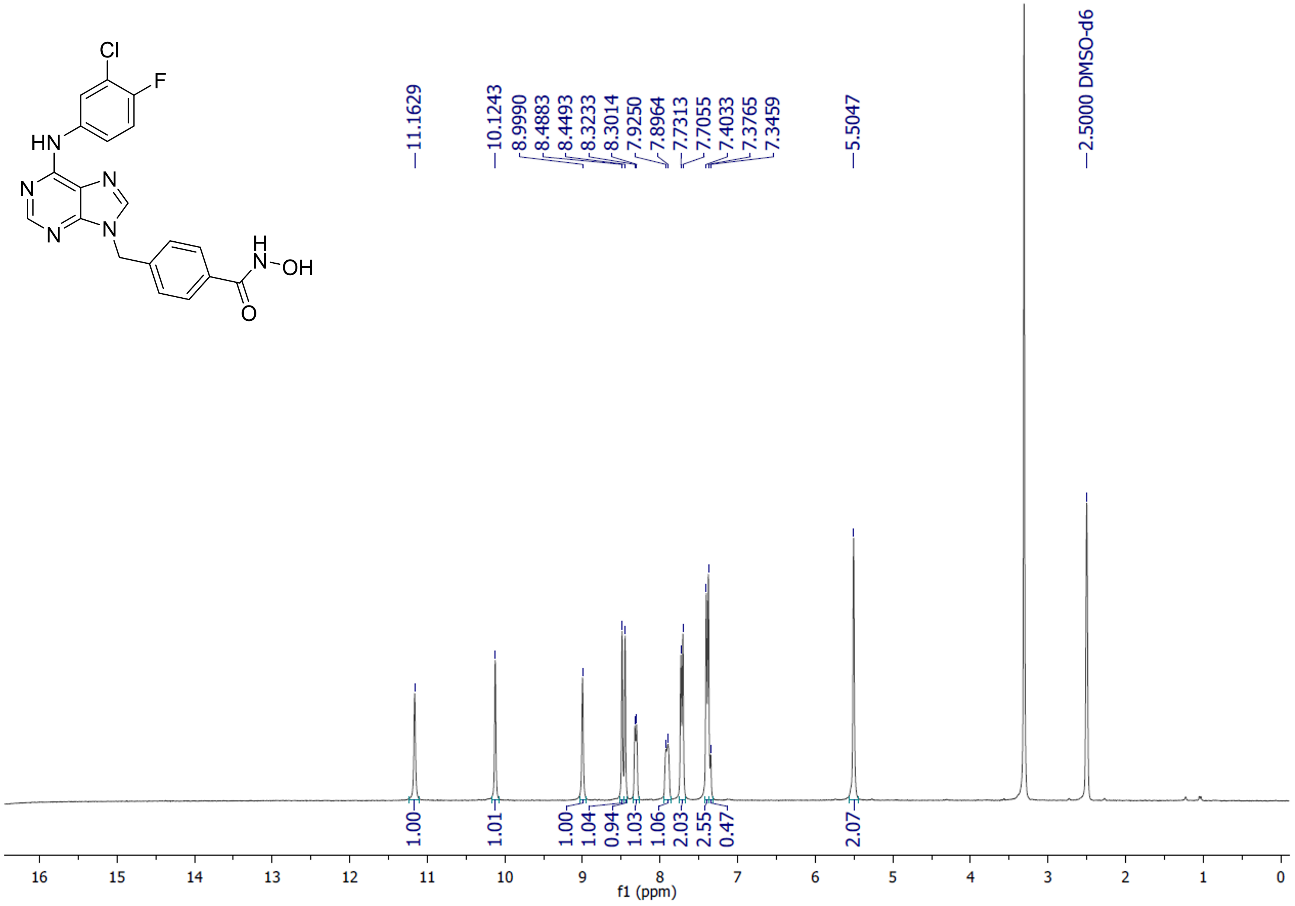

**Supplementary Figure S45:** ^13^C NMR spectra of the final compound 6d (75 MHz, DMSO-d_6_, δ = ppm)

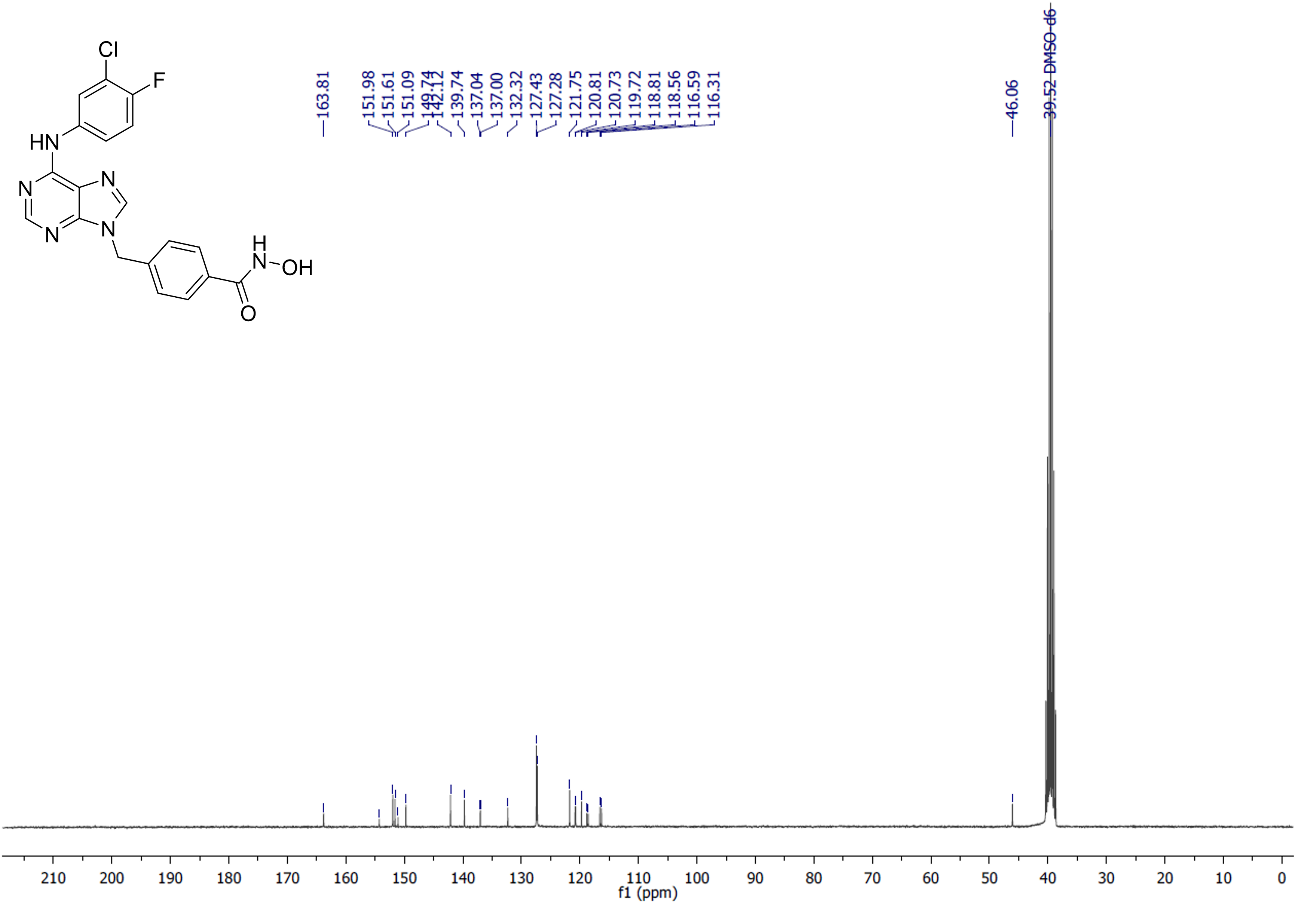

**Supplementary Figure S46:** ^1^H NMR spectra of the intermediate 6k (300 MHz, DMSO-d_6_, δ = ppm)

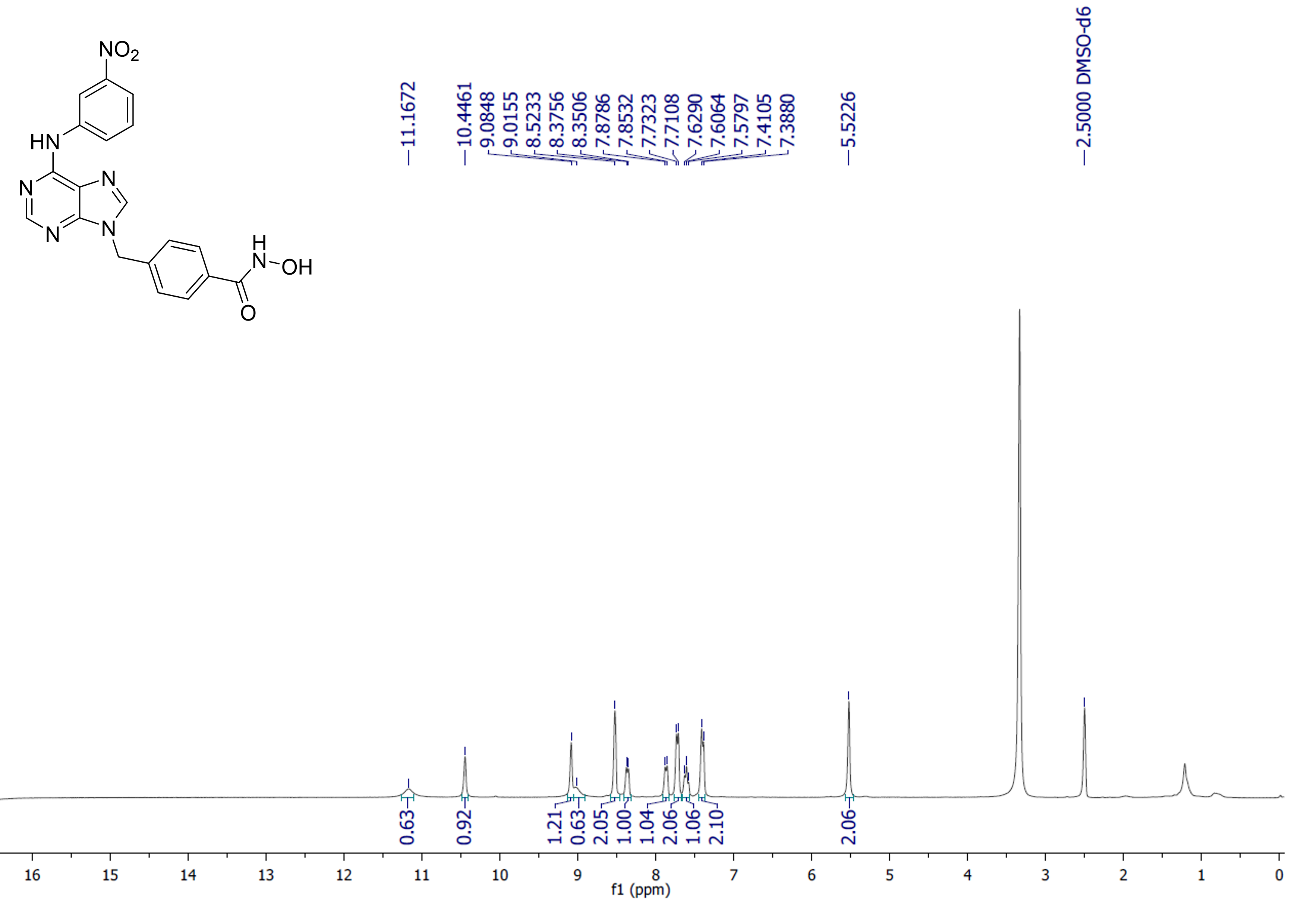

**Supplementary Figure S47:** ^13^C NMR spectra of the intermediate 6k (75 MHz, DMSO-d_6_, δ = ppm)

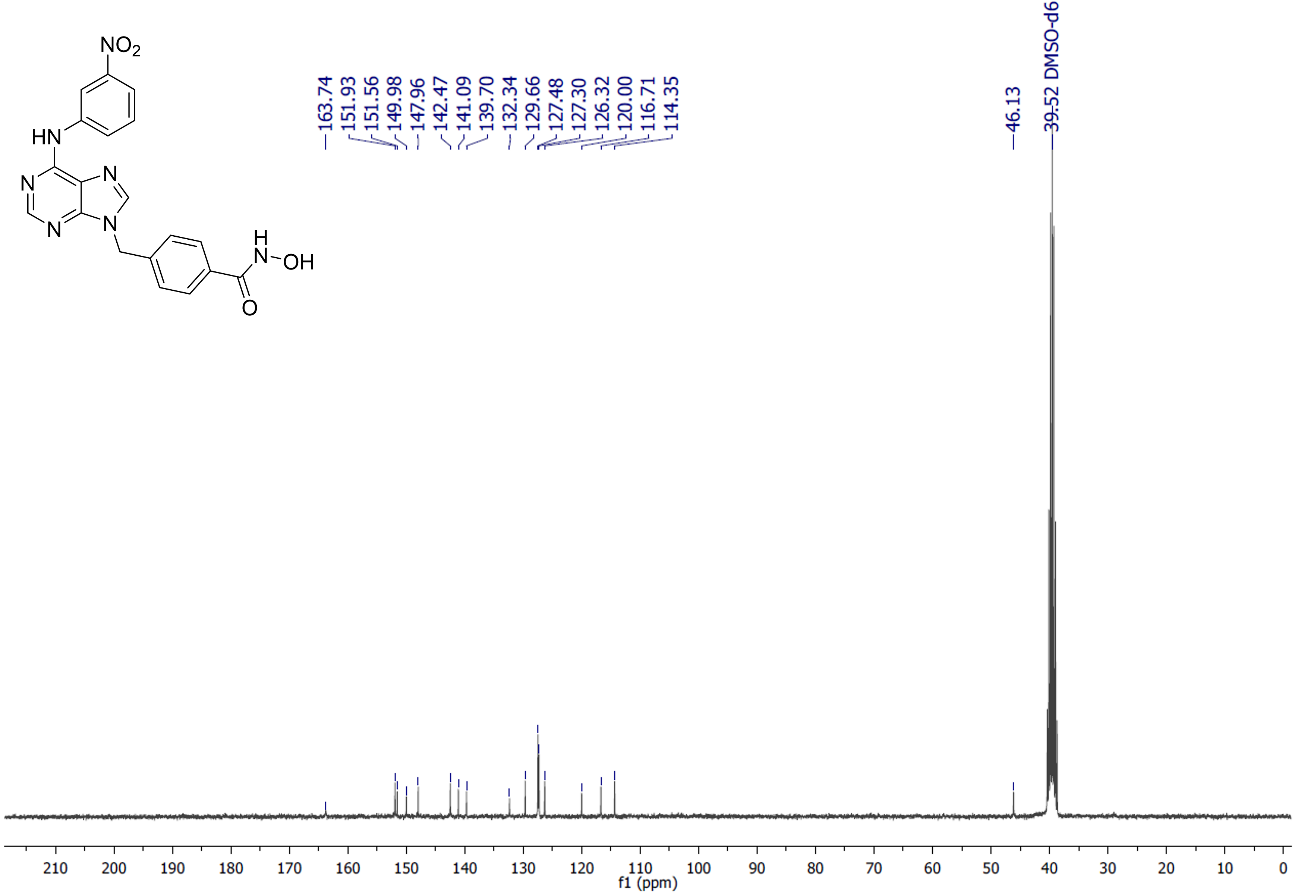

**Supplementary Figure S48:** HPLC spectra of the final compound 5a, page 1

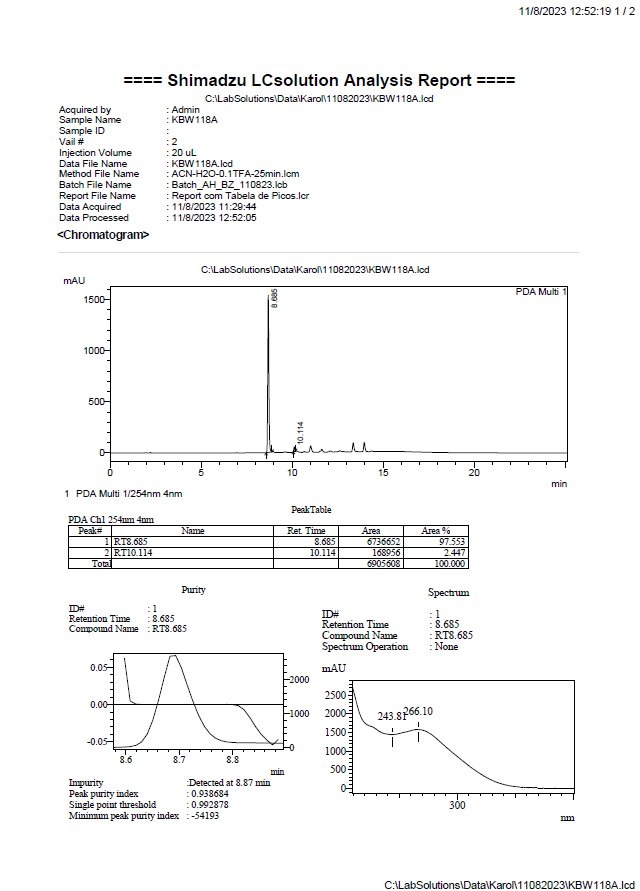

**Supplementary Figure S49:** HPLC spectra of the final compound 5a, page 2

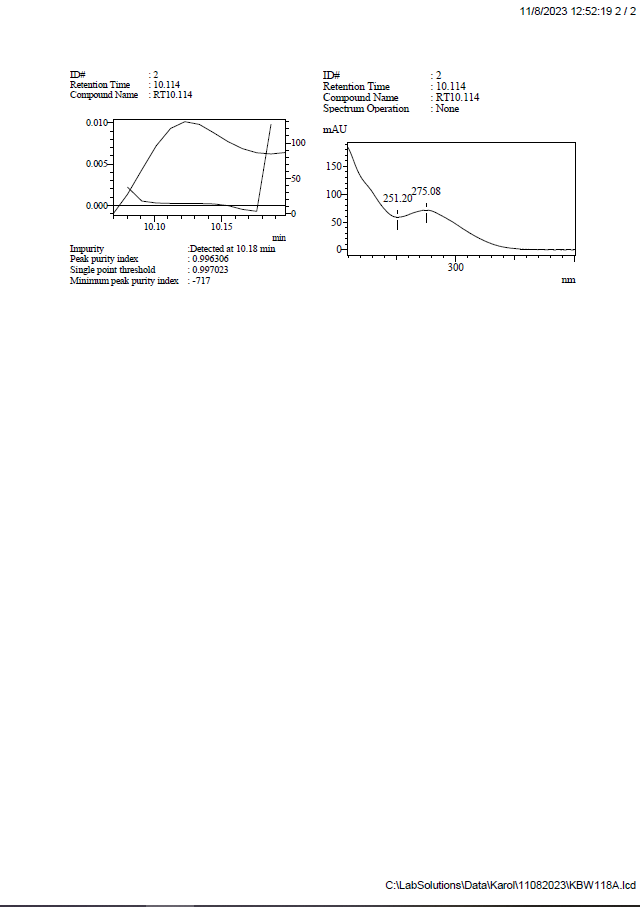

**Supplementary Figure S50:** HPLC spectra of the final compound 5b, page 1

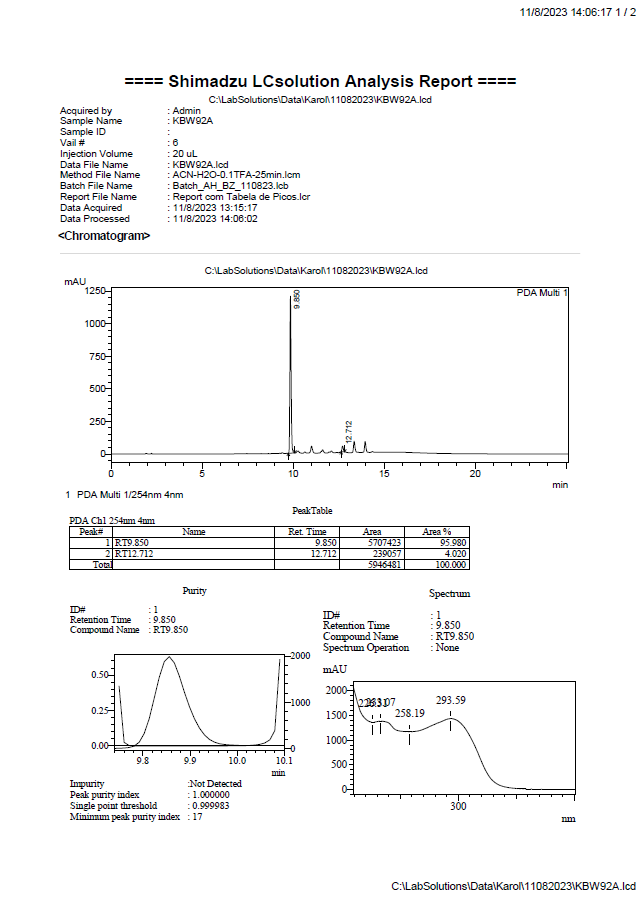

**Supplementary Figure S51:** HPLC spectra of the final compound 5b, page 2

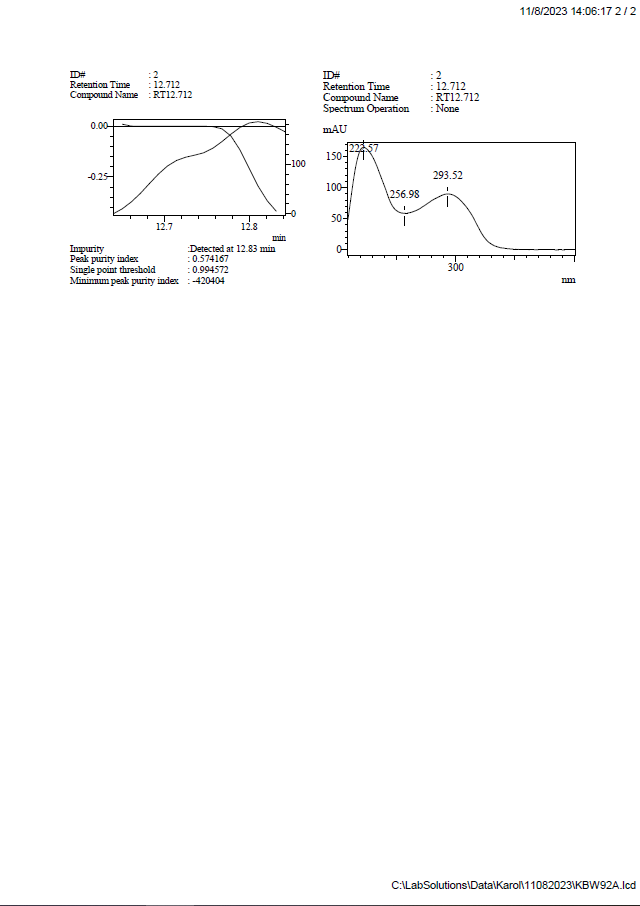

**Supplementary Figure S52:** HPLC spectra of the final compound 5c, page 1

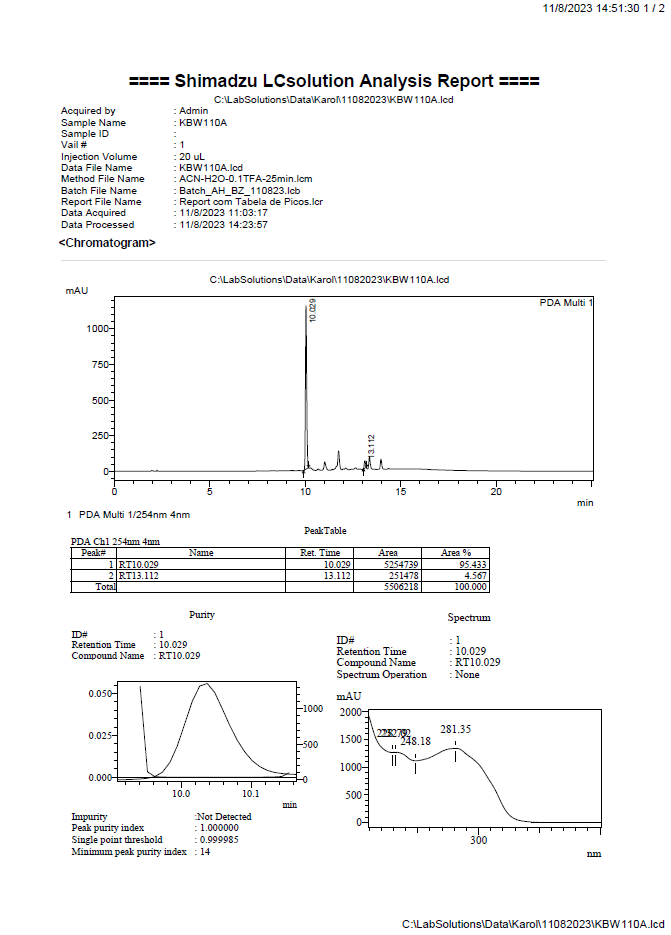

**Supplementary Figure S53:** HPLC spectra of the final compound 5c, page 2

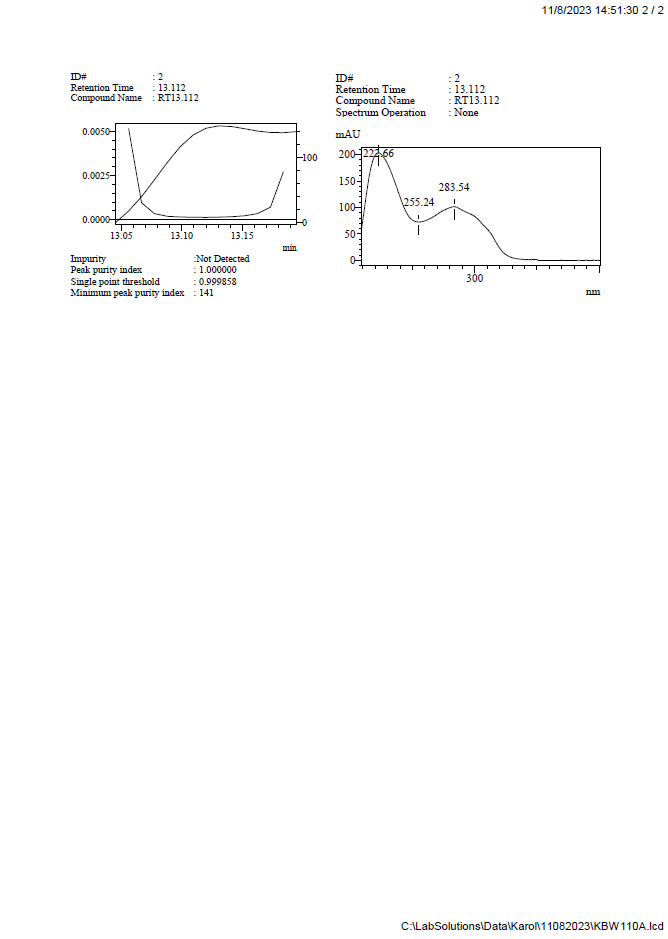

**Supplementary Figure S54:** HPLC spectra of the final compound 6a, page 1

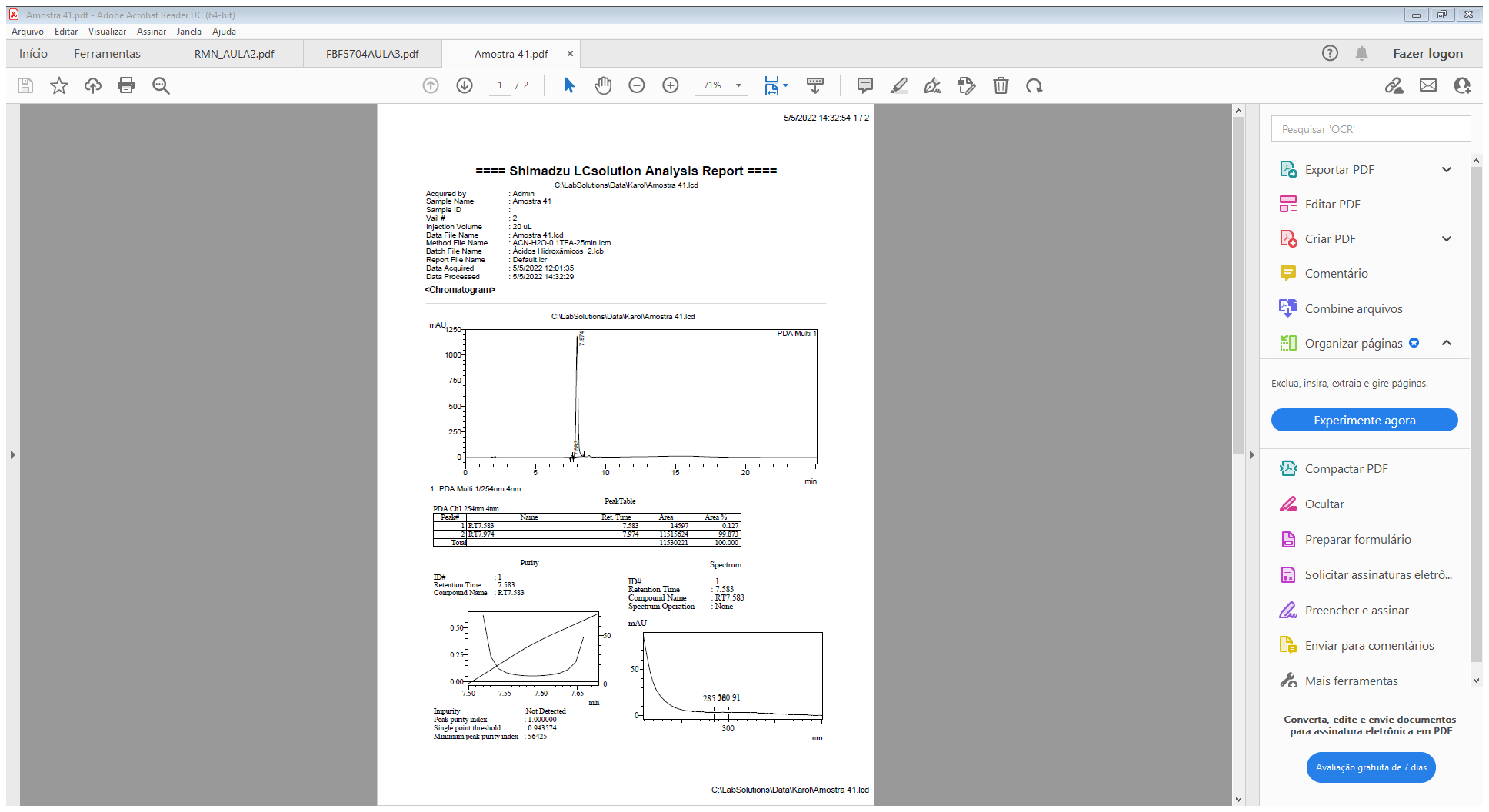

**Supplementary Figure S55:** HPLC spectra of the final compound 6a, page 2

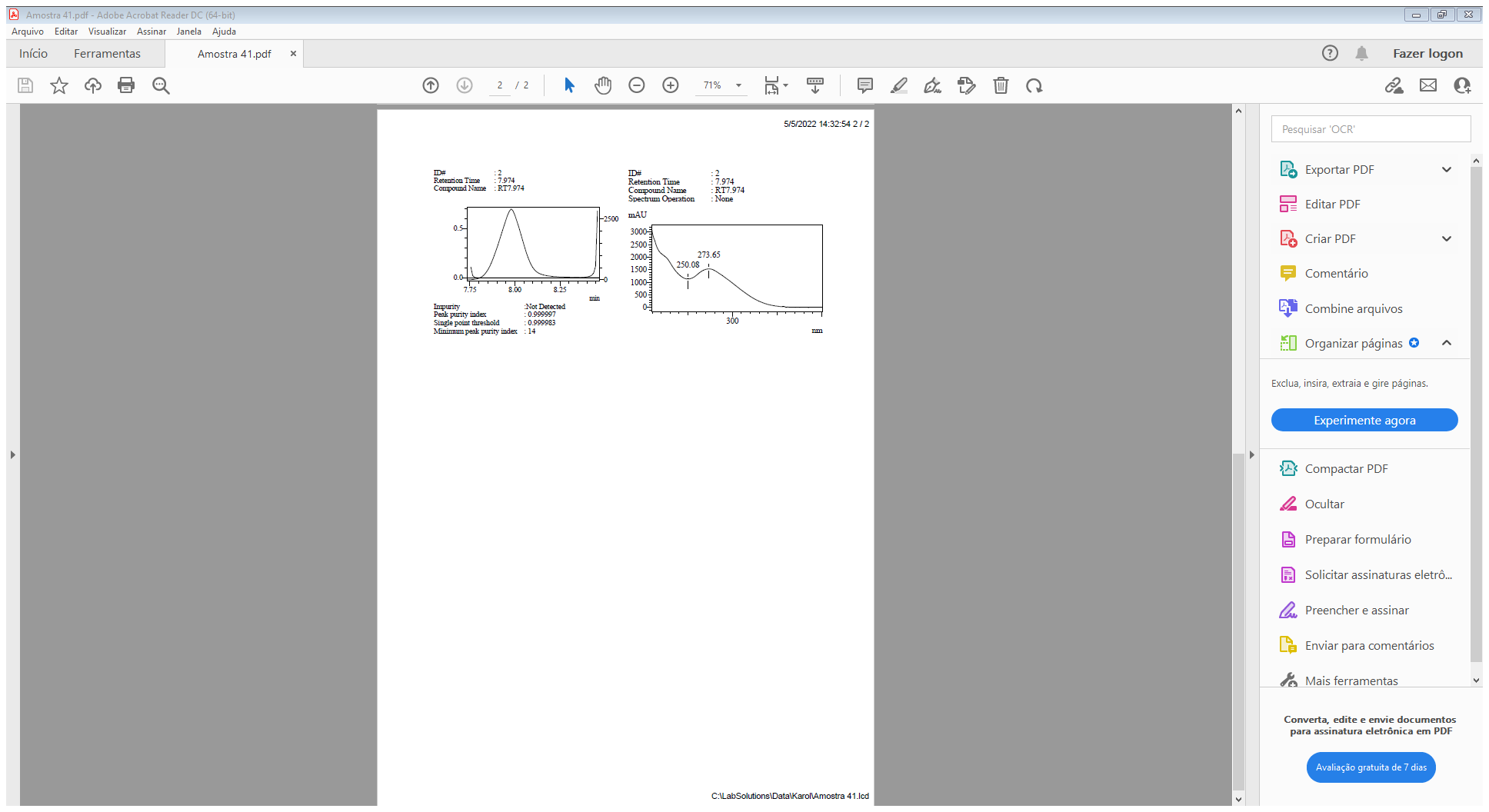

**Supplementary Figure S56:** HPLC spectra of the final compound 6b, page 1

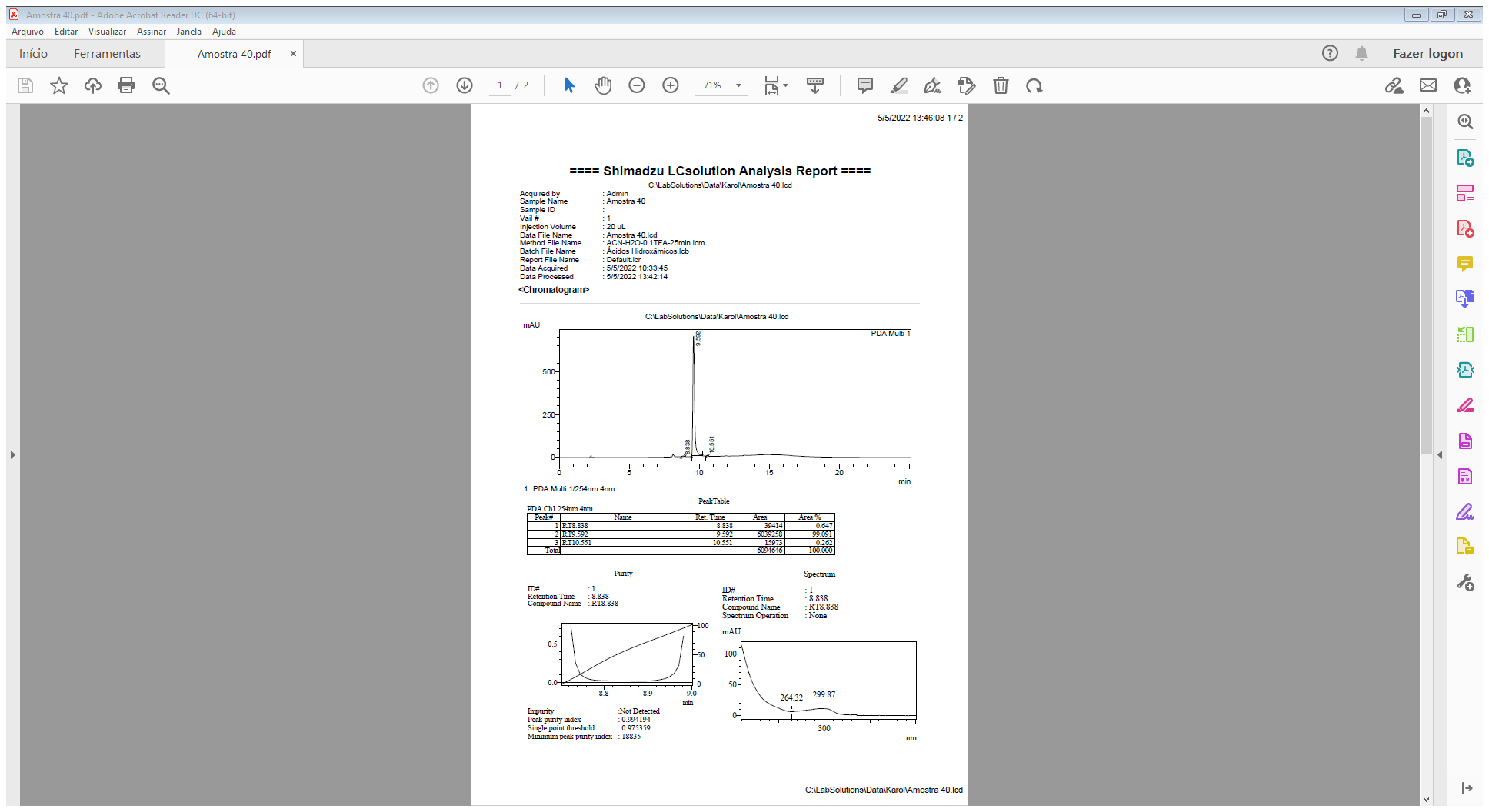

**Supplementary Figure S57:** HPLC spectra of the final compound 6b, page 2

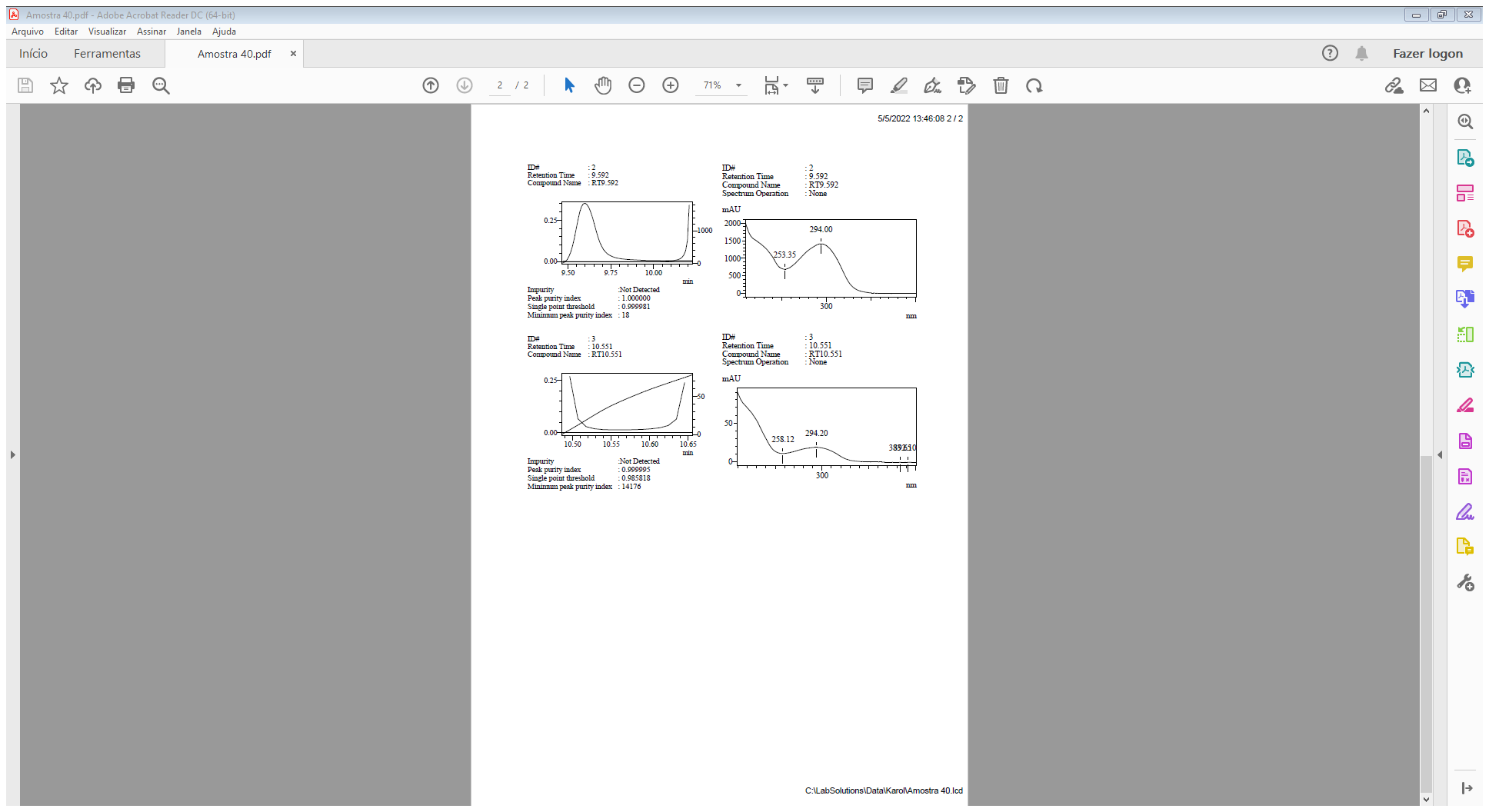

**Supplementary Figure S58:** HPLC spectra of the final compound 6c, page 1

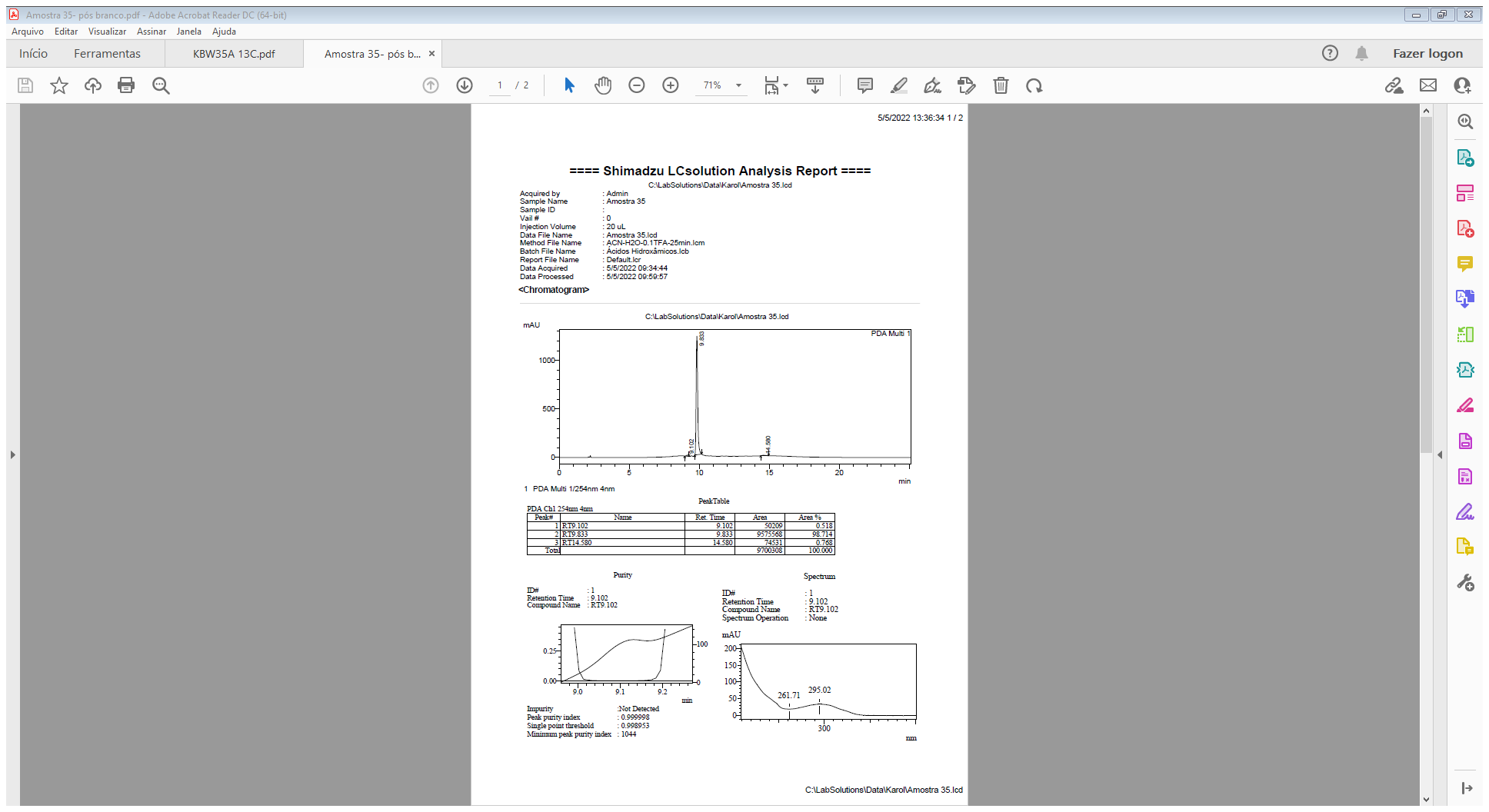

**Supplementary Figure S59:** HPLC spectra of the final compound 6c, page 2

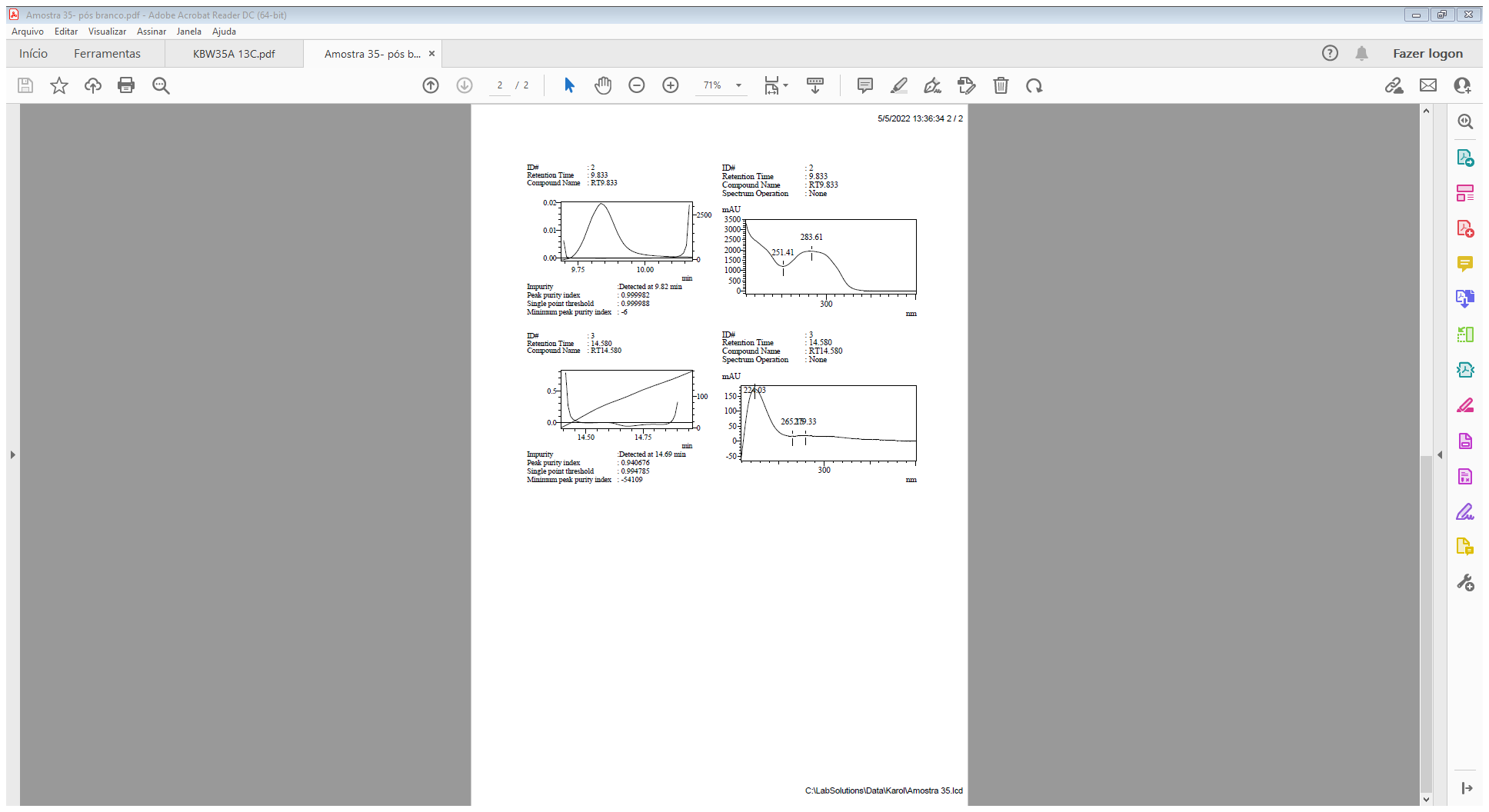

**Supplementary Figure S60:** HPLC spectra of the final compound 6d, page 1

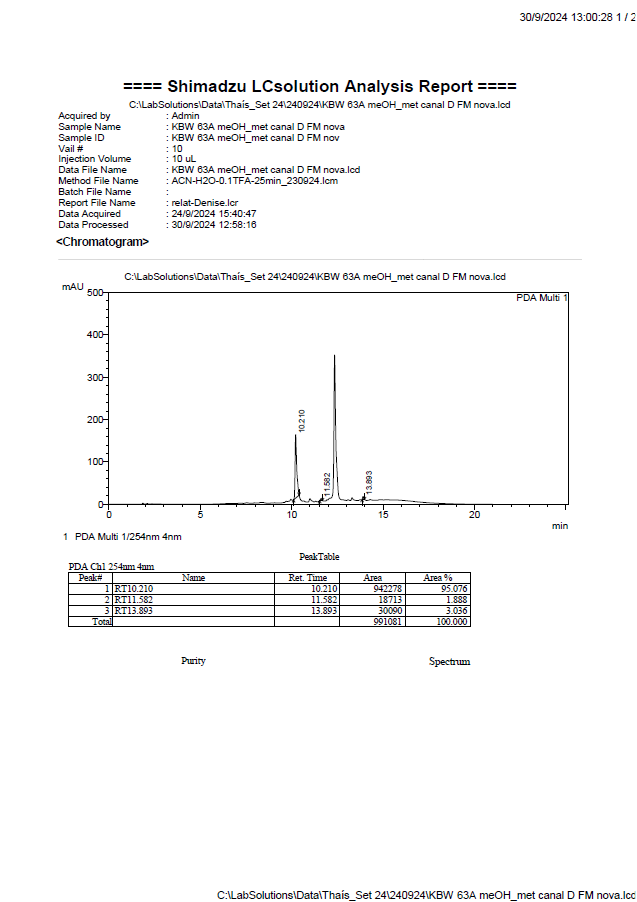

**Supplementary Figure S61:** HPLC spectra of the final compound 6d, page 2

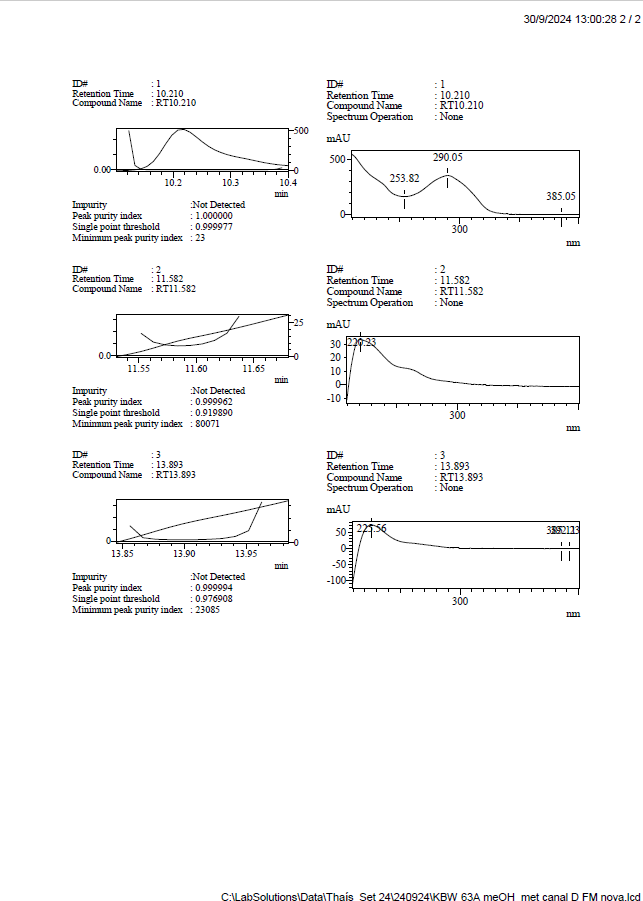

**Supplementary Figure S62:** HPLC spectra of the final compound 6k, page 1

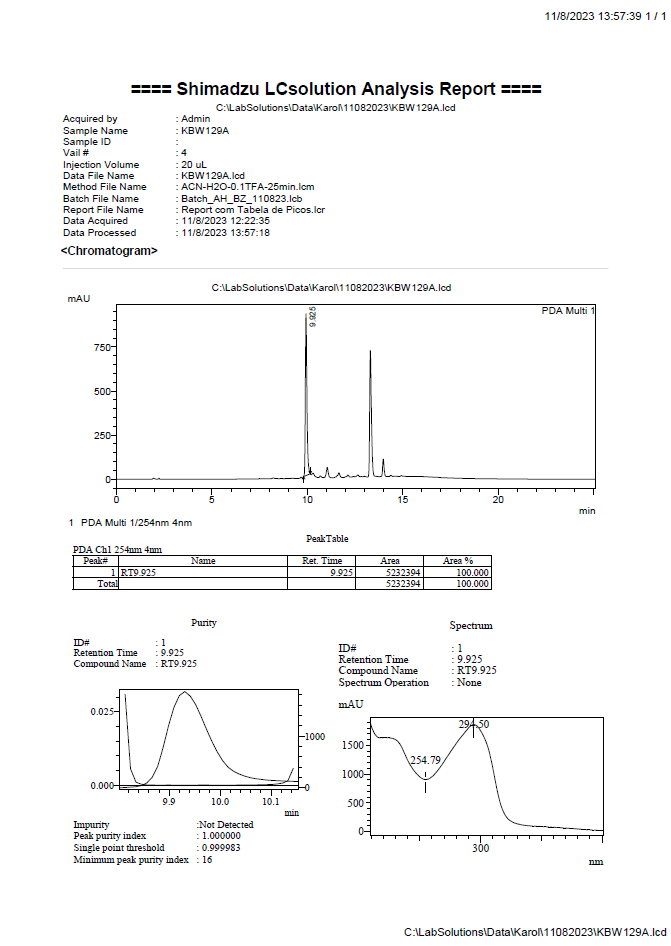
